# A familial, telomere-to-telomere reference for human *de novo* mutation and recombination from a four-generation pedigree

**DOI:** 10.1101/2024.08.05.606142

**Authors:** David Porubsky, Harriet Dashnow, Thomas A. Sasani, Glennis A. Logsdon, Pille Hallast, Michelle D. Noyes, Zev N. Kronenberg, Tom Mokveld, Nidhi Koundinya, Cillian Nolan, Cody J. Steely, Andrea Guarracino, Egor Dolzhenko, William T. Harvey, William J. Rowell, Kirill Grigorev, Thomas J. Nicholas, Keisuke K. Oshima, Jiadong Lin, Peter Ebert, W. Scott Watkins, Tiffany Y. Leung, Vincent C.T. Hanlon, Sean McGee, Brent S. Pedersen, Michael E. Goldberg, Hannah C. Happ, Hyeonsoo Jeong, Katherine M. Munson, Kendra Hoekzema, Daniel D. Chan, Yanni Wang, Jordan Knuth, Gage H. Garcia, Cairbre Fanslow, Christine Lambert, Charles Lee, Joshua D. Smith, Shawn Levy, Christopher E. Mason, Erik Garrison, Peter M. Lansdorp, Deborah W. Neklason, Lynn B. Jorde, Aaron R. Quinlan, Michael A. Eberle, Evan E. Eichler

## Abstract

Using five complementary short- and long-read sequencing technologies, we phased and assembled >95% of each diploid human genome in a four-generation, 28-member family (CEPH 1463) allowing us to systematically assess *de novo* mutations (DNMs) and recombination. From this family, we estimate an average of 192 DNMs per generation, including 75.5 *de novo* single-nucleotide variants (SNVs), 7.4 non-tandem repeat indels, 79.6 *de novo* indels or structural variants (SVs) originating from tandem repeats, 7.7 centromeric *de novo* SVs and SNVs, and 12.4 *de novo* Y chromosome events per generation. STRs and VNTRs are the most mutable with 32 loci exhibiting recurrent mutation through the generations. We accurately assemble 288 centromeres and six Y chromosomes across the generations, documenting *de novo* SVs, and demonstrate that the DNM rate varies by an order of magnitude depending on repeat content, length, and sequence identity. We show a strong paternal bias (75-81%) for all forms of germline DNM, yet we estimate that 17% of *de novo* SNVs are postzygotic in origin with no paternal bias. We place all this variation in the context of a high-resolution recombination map (∼3.5 kbp breakpoint resolution). We observe a strong maternal recombination bias (1.36 maternal:paternal ratio) with a consistent reduction in the number of crossovers with increasing paternal (r=0.85) and maternal (r=0.65) age. However, we observe no correlation between meiotic crossover locations and *de novo* SVs, arguing against non-allelic homologous recombination as a predominant mechanism. The use of multiple orthogonal technologies, near-telomere-to-telomere phased genome assemblies, and a multi-generation family to assess transmission has created the most comprehensive, publicly available “truth set” of all classes of genomic variants. The resource can be used to test and benchmark new algorithms and technologies to understand the most fundamental processes underlying human genetic variation.

## INTRODUCTION

The complete sequencing of a human genome was an important milestone in understanding some of the most complex regions of our genome^1^. Its completion added an estimated 8% of the most repeat-rich DNA, including regions typically excluded from studies of human genetic variation and recombination analyses, such as centromeres^2^, segmental duplications (SDs)^3^, and acrocentric regions^1,4^. Long-read sequencing has also driven assembly-based approaches to understand human genetic variation, revealing new insights into mutational mechanisms and access to regions previously considered intractable^5–7^. The ability to construct a phased genome assembly where the paternal and maternal complements are nearly fully resolved from telomere-to-telomere (T2T) opens up, in principle, the discovery of all forms of variation irrespective of class or complexity or the regions where they occur, placing them into the haplotypic context in which they immediately arose^8,9^. Direct comparison of parental genomes to their offspring increases the power to discover *de novo* mutation (DNM) as opposed to mapping reads to an intermediate reference, such as GRCh38 or T2T-CHM13^10^.

The goal of this study was to construct a high-quality T2T human pedigree resource where chromosomes were fully assembled and phased and their transmission studied intergenerationally to serve as a reference for understanding both recombination and DNM processes in the human species. We sought to eliminate three ascertainment biases with respect to discovery, including biases to specific genomic regions, classes of genetic variation, and reference genome effects. In addition to read-based approaches, we directly compare parent and child genomes to increase specificity and sensitivity of discovery in difficult regions of the genome, such as centromeres or chromosome Y. To achieve this, we focused on a four-generation, 28-member family, CEPH 1463, which has been intensively studied over the last three decades^11^, and sequenced members with five sequencing technologies having distinct and complementary error modalities. This particular pedigree has served as a benchmark for early linkage mapping studies^11,12^ and optimization of short-read sequencing data by Illumina^13^ and continues to serve as reference for understanding human variation, including patterns of mosaicism^14,15^.

Different from previous investigations, we focused our discovery on the sequencing and analysis of DNA obtained from primary tissue (i.e., peripheral blood leukocytes) as opposed to cell lines. We reconsented living family members (generations 2-3) and extended the sample collection to the fourth generation providing the opportunity to assess the transmission of DNMs. While all sequencing data and assemblies are available in dbGaP, 17 family members consented for their data to be publicly accessible similar to the 1000 Genomes Project samples. Just as the initial T2T genome^1^ served as a reference for understanding all regions of the genome, our objective was to create a reference truth set for both inherited and *de novo* variation. Our integration of multiple long- and short-read sequencing technologies across four generations allows us to understand the factors that affect the pattern and rates of DNMs in regions that were previously inaccessible.

## RESULTS

### Sequence and assembly of familial genomes

We generated PacBio high-fidelity (HiFi), ultra-long (UL) Oxford Nanopore Technologies (ONT), Strand-seq, Illumina, and Element AVITI Biosciences (Element) whole-genome sequencing (WGS) data for 28 members from the four-generation family (CEPH pedigree 1463) (**Fig. 1, Supplementary Table 1**). Individuals from the first to third generations (G1-G3) are members of the original CEPH pedigree^11^. The fourth generation (G4), as well as G3 spouses, are newly consented individuals. DNA for G2-G4 was extracted from peripheral whole blood leukocytes and is available both as primary material and cell lines. However, the great-grandparent generation (G1) are no longer living, thus DNA for G1 is only available as cell lines.

**Figure 1.**
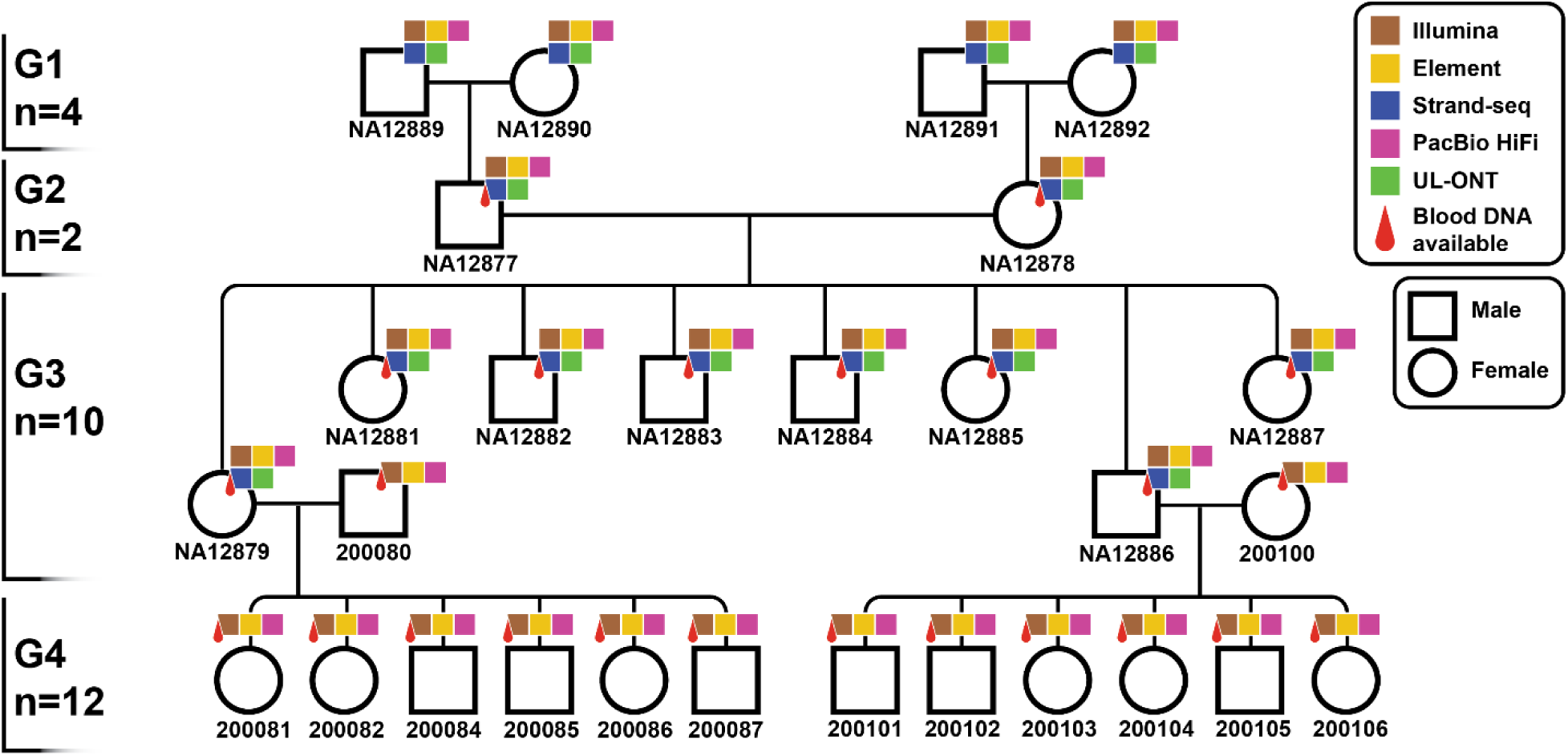
Sequencing the CEPH 1463 pedigree with five technologies. Twenty-eight members of the four-generation CEPH pedigree (1463) were sequenced using five orthogonal next-generation and long-read sequencing platforms: HiFi sequencing, Illumina, and Element sequencing for generations 2-4 (G2-G4) were performed on peripheral blood, while UL-ONT and Strand-seq were generated on available lymphoblastoid cell lines (G1-G3). The pedigree dataset has been expanded, for the first time, to include the fourth generation and G3 spouses (NA12879 and NA12886).

For the purpose of variant discovery, we focused on generating long-read PacBio, short-read Illumina, and Element data from blood-derived DNA to exclude DNA artifacts from EBV-transformed lymphoblasts. We also leveraged the corresponding cell lines to generate UL-ONT reads to construct near-T2T assemblies as well as Strand-seq data to detect large polymorphic inversions and evaluate assembly accuracy (**Methods**; **Supplementary Table 2**). In brief, we generated deep WGS data from multiple orthogonal sequencing platforms, focusing primarily on the first three generations (G1-G3) (**Extended Data Fig. 1a**), and used the fourth generation (G4) to validate *de novo* germline variants. We applied two hybrid genome assembly pipelines, Verkko^16^ and hifiasm^17^, to generate highly contiguous, phased genome assemblies for G1-G3, while G4 members were assembled using HiFi data only. We refer to hifiasm assemblies integrating UL-ONT data as ‘hifiasm (UL)’ while those that do not as ‘hifiasm’ only. Assemblies were phased using parental *k*-mers extracted from the high-coverage Illumina data for G2-G4 and with Strand-seq data for G1 samples (**Methods**).

Overall, Verkko assemblies are the most contiguous (contig AuN: 102 Mbp) followed by hifiasm (UL) and assemblies generated using HiFi reads alone (**Extended Data Fig. 1b**). Verkko-scaffolded contigs report even higher AuN value (134 Mbp) and contain a total of 896 gaps corresponding to an estimated gap size of 2.4 Mbp per assembled human genome haplotype (**Supplementary Fig. 1 and 2**). As expected, acrocentric chromosomes (13, 14, 15, 21 and 22) and chromosomes with secondary constrictions (chromosomes 1qh, 9qh and 16qh) composed of multiple megabase pairs of human satellite sequences (HSAT 1-3) were almost never completely assembled. Excluding acrocentric chromosomes, we estimate that 63.3% (319/504) of chromosomes across G1-G3 are spanned T2T in Verkko assemblies (**Extended Data Fig. 1c**). Telomere completeness was further evaluated showing 42.3% (213/504) of non-acrocentric chromosomes are spanned in a single contig and have canonical telomere repeats at each end (**Extended Data Fig. 1d, Supplementary Fig. 3, Supplementary Table 3, Methods**). Notably, we successfully sequenced and assembled 288 centromeres (44.7%) across the three generations, which required application of both Verkko and hifiasm (UL), as each assembler preferentially assembled different human centromeres (**Supplementary Fig. 4**). Verkko, for example, assembled 175 centromeres (27.2%) accurately, while hifiasm (UL) assembled 161 centromeres (25.0%) accurately. Only 48 centromeres (7.5%) were completely and accurately assembled by both Verkko and hifiasm (UL). Thus, by merging complete centromeres generated by both assemblers, we create a nonredundant list of 288 completely and accurately assembled centromeres (**Fig. 2e, Methods**).

**Figure 2.**
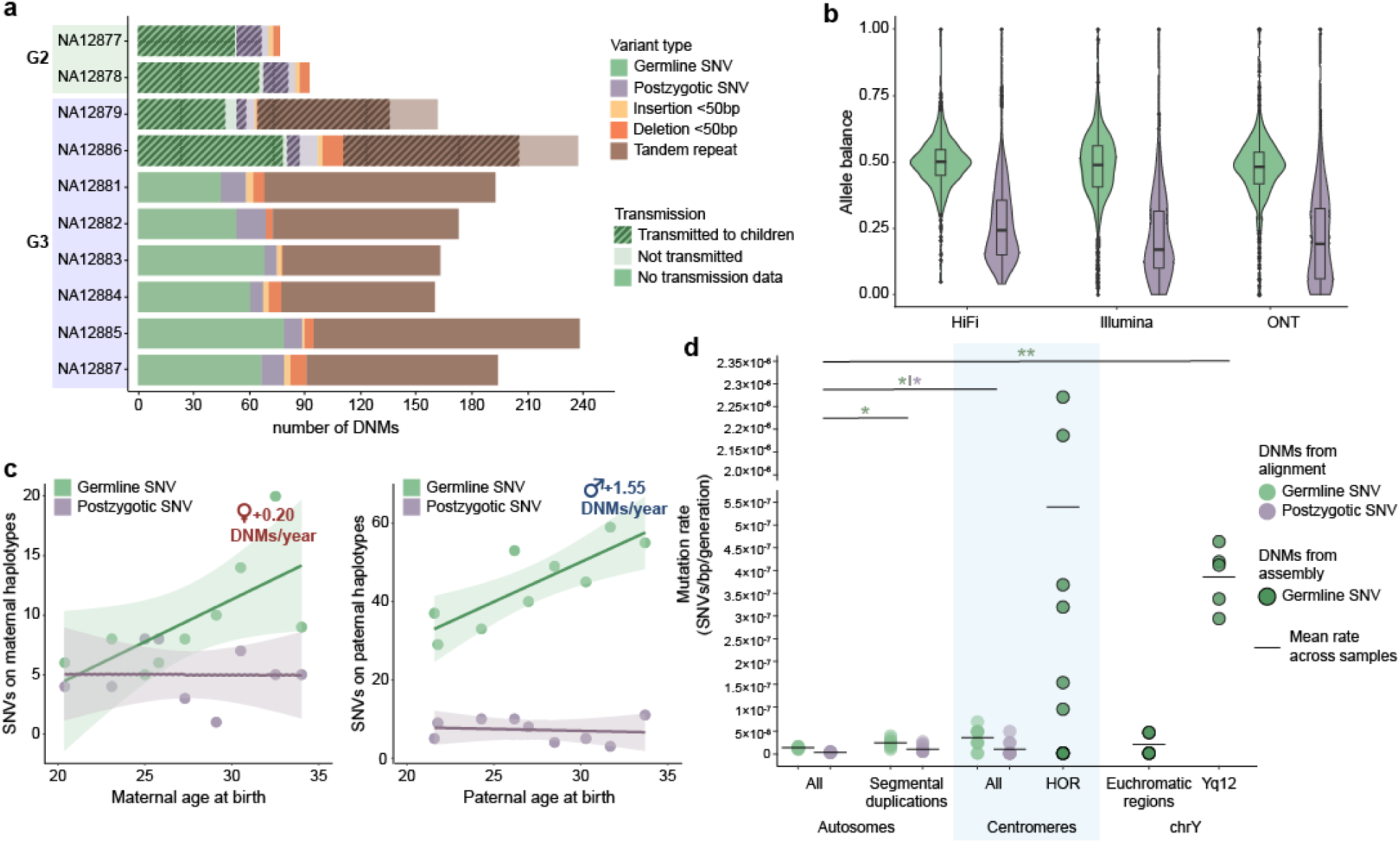
Summary of *de novo* mutation (DNM) rates. **a**) The number of *de novo* germline/postzygotic mutations (PZMs) and indels (<50 bp) for the parents (G2) and 8 children in CEPH 1463. Tandem repeat de novo mutations (TR DNMs) (<50 bp) are shown for G3 only because they have greater parental sequencing depth and we can assess transmission (**Methods**). Crosshatch bars are the number of SNVs confirmed as transmitting to the next generation. **b**) Germline SNVs have a mean allele balance near 0.50 across sequencing platforms, while the mean postzygotic allele balance is less than 0.25. **c**) A strong paternal age effect is observed for germline *de novo* SNVs but not for PZMs. **d**) Estimated SNV DNM rate by region of the genome shows a significant excess of DNM for large repeat regions, including centromeres and segmental duplications. Assembly-based DNM calls on the centromeres and Y chromosome show an excess of DNM in the satellite DNA.

Using Illumina WGS data (**Methods**), we estimate the accuracy of the Verkko assemblies at quality value 54 on average (range: 47-58) (**Supplementary Fig. 5**). In addition, we tested structural and phasing accuracy of our Verkko assemblies. Our Strand-seq data confirms a low misorientation rate (<0.022%) (**Supplementary Fig. 6**) and a high phasing accuracy with Hamming error rates <2%, which is further supported by pedigree-based phasing of G2-G3 (**Supplementary Fig. 7**). We detected, however, four extended haplotype switch errors (from ∼500 kbp to 3.7 Mbp in size) that have been corrected in our assembly-based variant callsets to avoid biases in subsequent analysis. Lastly, we note a single chimeric contig in the Verkko assembly of G3-NA12886 (**Supplementary Figs. 8 and 9, Supplementary Table 4**).

We systematically assessed the assembled chromosomes for other collapses and misjoins using Flagger^9^ and NucFreq^18^. Flagger reports on average >98% (5.91 Gbp) of each phased assembly being assembled at the correct copy number with one outlier sample (G3-NA12879) with an excess of potential collapses (**Supplementary Fig. 10, Methods**). However, this observation is not supported by the alternate validation tool NucFreq, suggesting that this particular sample may be subject to a less uniform sequence coverage (**Supplementary Fig. 11**). When considering alignment of phased assemblies to the T2T-CHM13 reference, we report that on average ∼97% (2.88 Gbp) of each phased assembly is fully alignable to the reference at expected diploid copy number (**Supplementary Fig. 12, Methods**). Last, we identify a relatively small number of misjoins in our assemblies (n=47, median: 2 per haploid assembly) (**Supplementary Fig. 13**) along with >98% completeness of single-copy genes (**Supplementary Fig. 14, Methods**).

**Extended Data Figure 1.**
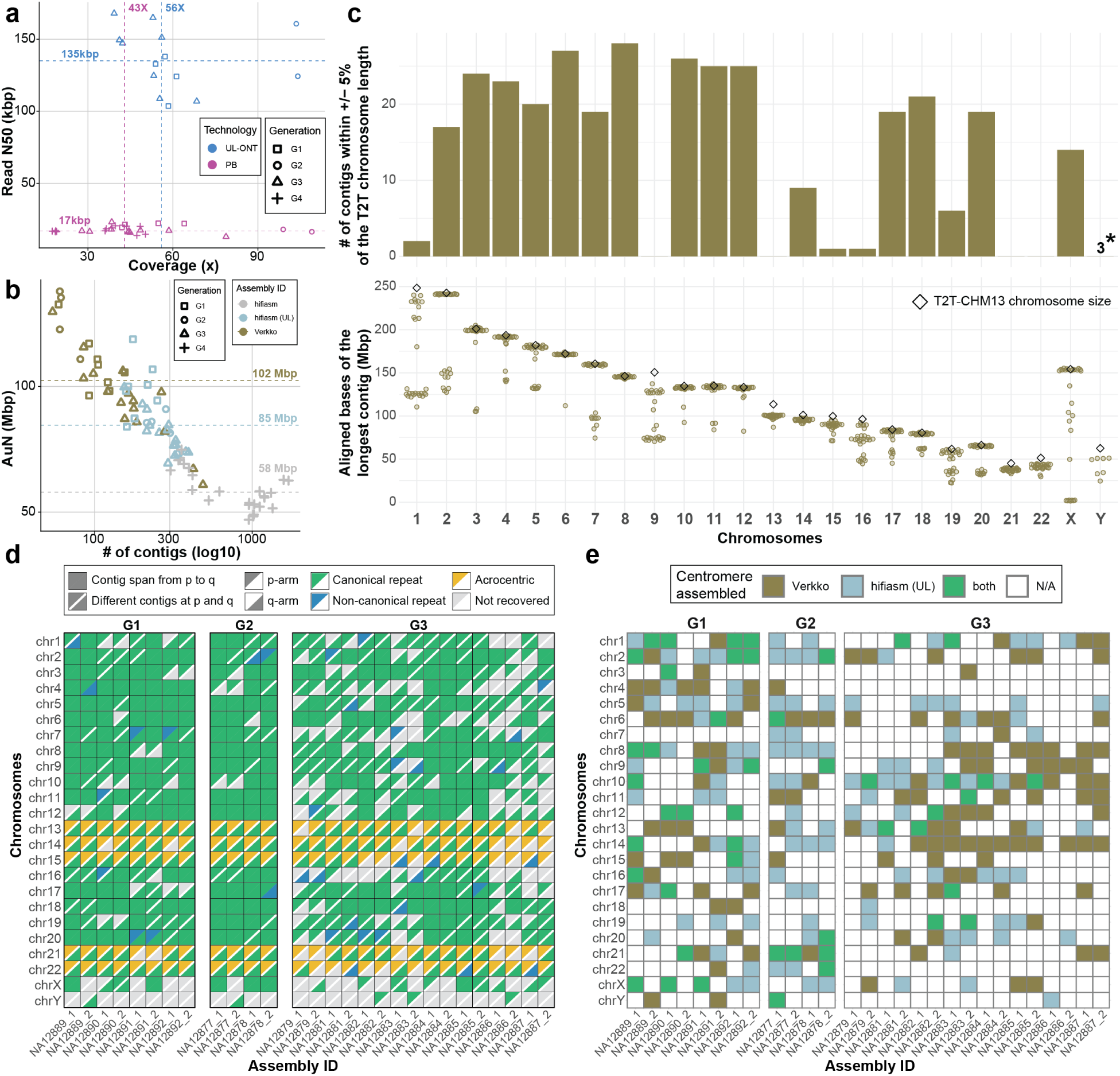
Long-read sequencing and assembly contiguity. **a**) Scatterplot of sequence read depth and read length N50 for ONT (blue) and PacBio (PB; magenta) with median coverage (dashed line) and different generations indicated (point shape). **b**) Scatterplot of the assembly contiguity measured in AuN values for Verkko (brown), hifiasm (UL) (light blue), and hifiasm (light gray) assemblies of G1-G4. Note: G4 samples were assembled using PacBio HiFi data (hifiasm) only. **c**) Top: Total number of Verkko contigs whose maximum aligned bases are within +/-5% of the total T2T-CHM13 chromosome length. **\***Due to substantial size differences between the T2T-CHM13 Y (haplogroup J1a-L816) and the Y chromosome of this pedigree (haplogroup R1b1a-Z302), three contigs are shown that span the entire male-specific Y region without breaks (i.e., excluding the pseudoautosomal regions). Bottom: Each dot represents a single Verkko contig with the highest number of aligned bases in a given chromosome. **d**) Chromosomes containing complete telomeres and being spanned by a single contig are annotated as solid squares. In instances where the p- and q-arms are not continuously assembled and for acrocentric chromosomes, we plot diagonally divided and color-coded triangles. **e**) Evaluation of centromere completeness across G1-G3 assemblies and across all chromosomes. We mark centromeres assembled by Verkko (brown), hifiasm (UL) (light blue), or both (green).

### A multigenerational variant callset

Having contiguous assemblies as well as read-based data from multiple technologies allows us, in principle, to track the inheritance of any genomic segment and associated variants across all four generations with high specificity (**Extended Data Fig. 2a**). We used the sequence reads (PacBio, Illumina and ONT) as well as the assemblies to create a union of all genetic variants. We consider three classes of variants: single-nucleotide variants (SNVs; single-base-pair variants), indels (1-49 bp insertion/deletions) and structural variants (SVs ≥50 bp), including inversions, and leverage the multigenerational nature of the pedigree^13^ to directly validate genetic variation through haplotype transmission. We establish a variant truth set to be used for subsequent analyses and identify a total of 8.8 million SNVs/indels and 35,685 SVs of which 95% and 70%, respectively, show evidence of transmission from G2-G3 (**Supplementary Table 5**). In total, we identify 6.05 million pedigree-consistent small variant alleles against GRCh38, of which 4.6 million (76.0%) are supported by all three technologies and callers. Leveraging long-read sequence data in the context of a pedigree provides access to an additional ∼244 Mbp of the human genome (2.76 Gbp high-confidence regions) when compared to Genome in a Bottle (GIAB) (2.51 Gbp)^19^ or Illumina WGS data (2.58 Gbp)^13^, including 194 Mbp not present in either study. Some of the largest gains occur among SDs and the genes associated with them. In this analysis, for example, we classified 83.7% of the SDs (coverage >95%) as high-confidence regions compared to a previous GIAB analysis, which called variants only in 25.6% of these regions. Among the SVs, we identify 2,161 *Alu* insertions, 398 LINE-1 insertions, and 152 SINE-VNTR-Alu (SVA) retrotransposon insertions (**Supplementary Fig. 15**). Only *Alu* elements >260 bp were included in this analysis. We identify 112 LINE-1 insertions of either full-length or near full-length (at least 5,500 bp) and 124 SVA insertions of at least 2,000 bp (**Supplementary Table 6, Supplementary Notes**). Using Strand-seq data, we also detect a total of 120 segregating simple inversions and 17 inverted duplications with median size of ∼53 kbp and ∼41 kbp, respectively (**Supplementary Table 7, Supplementary Figs. 16-19, Methods**). This includes a rare inversion (∼703 kbp) overlapping a morbid copy number variant region at 15q25.2^20^ (**Supplementary Fig. 20**) and an inverted duplication (∼295 kbp) at 16q11.2. We find that the region (∼63 kbp) between this inverted duplication is specifically inverted in this family, which changes the orientation of *UBE2MP1*—a pseudogene whose expression was recently linked to negative outcomes in hepatocellular carcinoma patients^21^ (**Supplementary Fig. 21**).

### Sequence-resolved recombination map

Using three different approaches (**Methods**), including transmission of assembled chromosomes (**Extended Data Fig. 2b**), we construct a high-resolution recombination map and identify 539 meiotic breakpoints in G3 (n=8) with respect to T2T-CHM13, with 99.8% supported by more than one approach (**Supplementary Fig. 22**). Strand-seq analysis of G1 assemblies allows us to phase and determine parent-of-origin for G2 chromosomes adding 139 breakpoints^22^. In total across the two generations (G2-G3; 10 transmissions), we identify 678 meiotic breakpoints (**Extended Data Fig. 2c, Supplementary Fig. 23, Supplementary Table 8**), including 15 recombination “hotspots”, 10 of which are in line with previously reported increased recombination rates^23^ (**Supplementary Fig. 24**). We also characterize 78 smaller haplotype segment “switches” (median size of ∼1 kbp) that would be consistent with either a double crossover or allelic gene conversion event, although many (n=17) overlap with low-complexity DNA. We validate eight events based on visual inspection of HiFi sequence data, including an event at chromosome 8p22 that overlaps the two protein-coding genes: *VPS37A* and *MTMR7* (**Supplementary Table 9, Supplementary Fig. 25, Methods**).

We find that ∼19% of paternal and maternal homologs are transmitted without a detectable meiotic breakpoint (i.e., nonrecombinant chromosomes) while the remainder (∼81%) contain at least one recombination breakpoint (**Supplementary Fig. 26**). We observe five regions ranging from ∼200 kbp to ∼19.4 Mbp that are inherited from a single grandparental homolog while the other homolog is essentially lost in the subsequent generations of this family (**Supplementary Figs. 27 and 28, Methods**). In line with previous research we observe a significant excess (two-sided t-test, p=0.0031) of maternal recombination events (1.36 maternal:paternal ratio) with chromosomes 8 and 10 showing the most significant maternal excess (z-score > 2.3; p < 0.02)^24^ (**Supplementary Fig. 29**). Paternal recombination is significantly biased towards the ends of human chromosomes with 31 paternal recombination events mapping within 2 Mbp of the telomere compared to none in females creating a bimodal paternal distribution of inherited segment lengths (**Extended Data Fig. 2d, Supplementary Fig. 30, Methods**). We observe a significant decrease in crossover events with advancing parental age for both male (R = −0.86; p = 0.0014; Pearson correlation) and female (R = −0.66; p = 0.039; Pearson correlation) germlines^25,26^ (**Extended Data Fig. 2e**).

We initially narrowed recombination breakpoint regions to ∼3.5 kbp; however, using the phased genome assemblies, including direct comparisons between parent and child (**Methods**), we further refined 90.4% (487/539) of the recombination events to a median size of ∼2.5 kbp (**Supplementary Fig. 31**). Surprisingly, only about half of the intervals are reduced (n=248) while 191 breakpoints actually increase when compared to T2T-CHM13 likely pointing to reference biases artificially truncating recombination intervals (**Supplementary Fig. 32**). We observe two types of recombination intervals: those with a very sharp transition between parental haplotypes and those with an extended region of homology at both parental haplotypes (**Extended Data Fig. 2b, Supplementary Fig. 33**). Last, we characterized the *PRDM9* genotypes for all individuals in the pedigree (**Supplementary Table 10**), comparing the results obtained from Verkko and hifiasm (UL) assemblies across the G1-G3 samples. Across the entire family, we define the alleles A, B, M10, and M19—all four from the PRDM9-A-type predicted binding site group^27^.

**Extended Data Figure 2.**
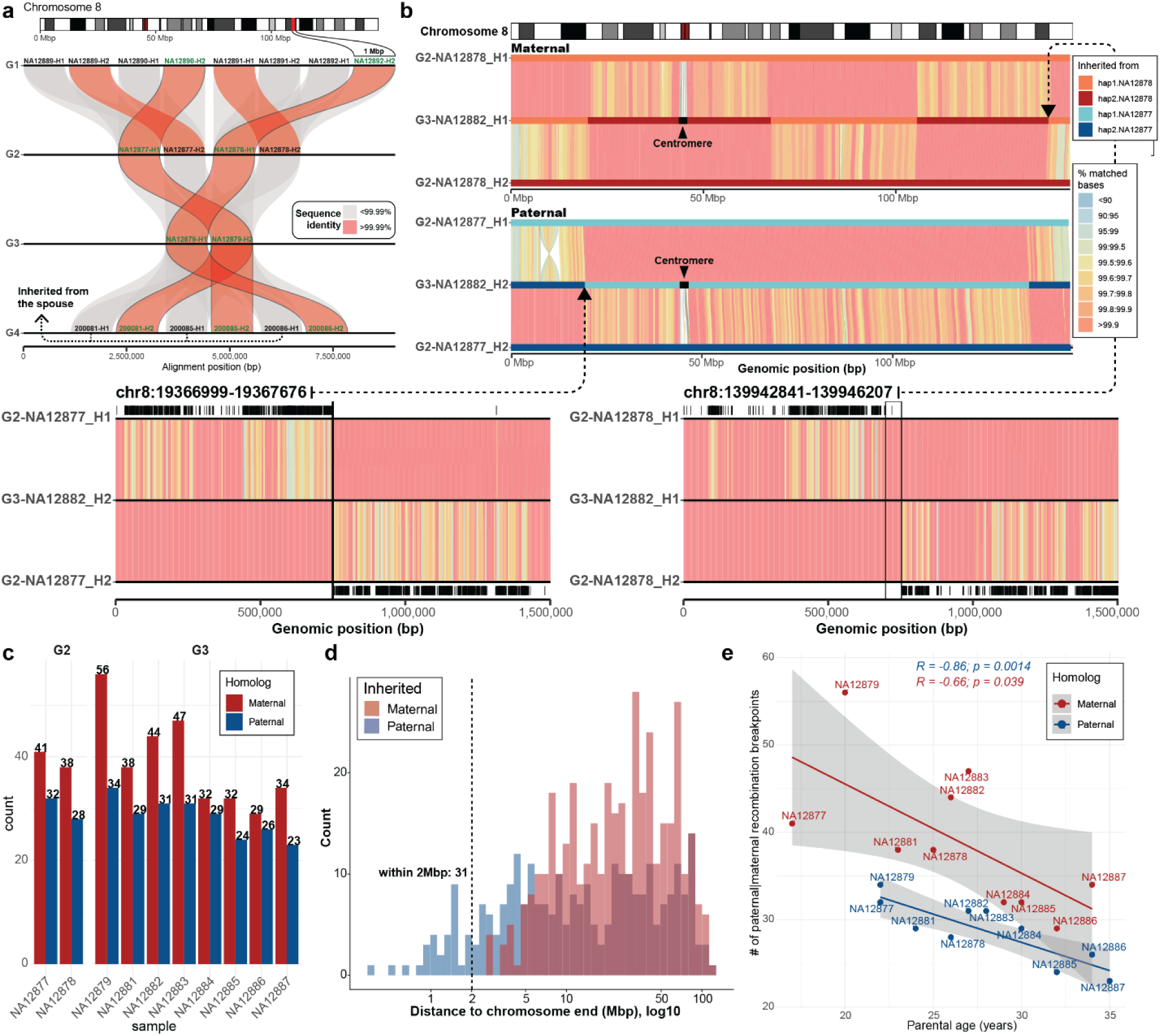
Recombination breakpoint map of CEPH 1463. **a**) Depiction of intergenerational (G1->G4) inheritance of a 1 Mbp assembled contig. Alignments transmitted between generations that are >99.99% identical (red) are contrasted with non-transmitted with lower sequence identity (gray). **b**) T2T recombination between child and parental haplotypes for chromosome 8. Alignments between parental and a child’s haplotypes are binned into 500 kbp long bins and colored based on the percentage of matched bases. Inherited maternal (shades of red) and paternal (shades of blue) segments are marked on top. Dashed arrows show zoom-in of the two recombination breakpoints that differ in size of the region of homology at the recombination breakpoint. Black tick marks show positions of mismatches between parental and child haplotypes. **c**) Summary of recombination breakpoints detected in inherited maternal (red) and paternal (blue) homologs with respect to T2T-CHM13. **d**) Distribution of distances of maternal (red) and paternal (blue) recombination breakpoints to chromosome ends. **e**) Correlation between the number of recombination breaks (y-axis) and parental age (x-axis) shown separately for maternal (red) and paternal (blue) recombination breakpoints.

### De novo SNVs and small indels

To discover small DNMs outside of tandem repeats (TRs), we initially map HiFi reads to T2T-CHM13 for every sample in the pedigree, selecting all variants that are not observed in parents (**Supplementary Table 11, Methods**). We partition variants into SNVs or indels based on length and then validate each variant by requiring orthogonal support with ONT and/or Illumina (i.e., present in child and absent in parents). By this criterion, we discover 755 *de novo* SNVs and 73 *de novo* indels across the autosomes of 10 individuals (n=2 G2; n=8 G3 individuals, Fig. 2a), as well as 27 *de novo* SNVs and 1 indel on the X chromosome. Using flanking SNVs from long-read sequencing data to construct haplotypes as well as allele balance for unphased variants, we categorize *de novo* variants as either germline DNMs or postzygotic mutations (PZMs) (**Fig. 2b, Methods**). We classify 17.1% (129/755) of *de novo* SNVs as PZMs, defined here as somatic mutations occurring very early in development. Of the 311 *de novo* SNVs in G2 and G3 individuals with offspring, 97.1% of germline events transmit to the next generation, compared with 64.5% of postzygotic events (**Extended Data Fig. 3**). All 28 indels in individuals with offspring are transmitted to the next generation. A previous Illumina-based study of this family^14^ identified a total of 605 *de novo* SNVs of either germline (G2 and G3) or postzygotic (only G2) origin, for an average of 59.0 DNMs and 7.5 PZMs per sample. We recover 92.4% (n=559) of these events in our final callset, and only four of the remaining variants pass our validation filters when considering HiFi and ONT data. Conversely, we identify an additional 196 DNMs, including 76 postzygotic events in G3 for the first time. Thus, our approach increases germline SNV discovery by 6.1% and indel discovery by 21%.

We find that 81.4% of germline small DNMs originate on paternal haplotypes (4.38:1 paternal:maternal ratio, Wilcoxon signed-rank test, p<2×10^−16^), whereas PZMs show no significant difference with respect to parental origin (1.35:1 paternal:maternal ratio, Wilcoxon signed-rank test, p = 0.123). In addition, we observe a significant parental age effect of 1.55 germline DNMs per additional year of paternal age when fitting with linear regression (p=0.013)—a signal absent from *de novo* SNVs designated as PZMs (Fig. 2c). The mutational spectra of DNM and PZM differ from each other, with a depletion of CpG>TpG PZMs, although this difference does not yet reach statistical significance based on sample size (**Supplementary Fig. 34a**). Using this approach, we successfully assay 91.9% of the autosomal genome (2.66 Gbp) with an overall SNV mutation rate of 1.39×10^−8^ SNVs/bp/generation (95% CI: 1.22 - 1.56×10^−8^) (**Supplementary Fig. 34b**). Based on our postzygotic and germline classification, we determined the germline contributes 1.17×10^−8^ SNVs/bp/generation (95% CI: 1.02 - 1.32×10^−8^). *De novo* SNVs are significantly enriched in repetitive sequences, as much as 2.8-fold in centromeres (95% CI: 1.65 - 4.88×10^−8^ SNVs/bp/generation, two-sided t-test, p=0.017) and 1.9-fold in SDs (95% CI: 1.53 - 2.86×10^−8^ SNVs/bp/generation, two-sided t-test, p=0.0066) (**Fig. 2d, Supplementary Fig. 34c**). We observe a lower PZM rate of 2.23×10^−9^ SNVs/bp/generation (95% CI: 1.74 - 2.37×10^−9^) across the autosomes, yet we see 4.5-fold enrichment of PZMs in SDs (95% CI: 4.46×10^−9^ - 1.57×10^−8^ SNVs/bp/generation, two-sided t-test, p=0.011).

**Extended Data Figure 3.**
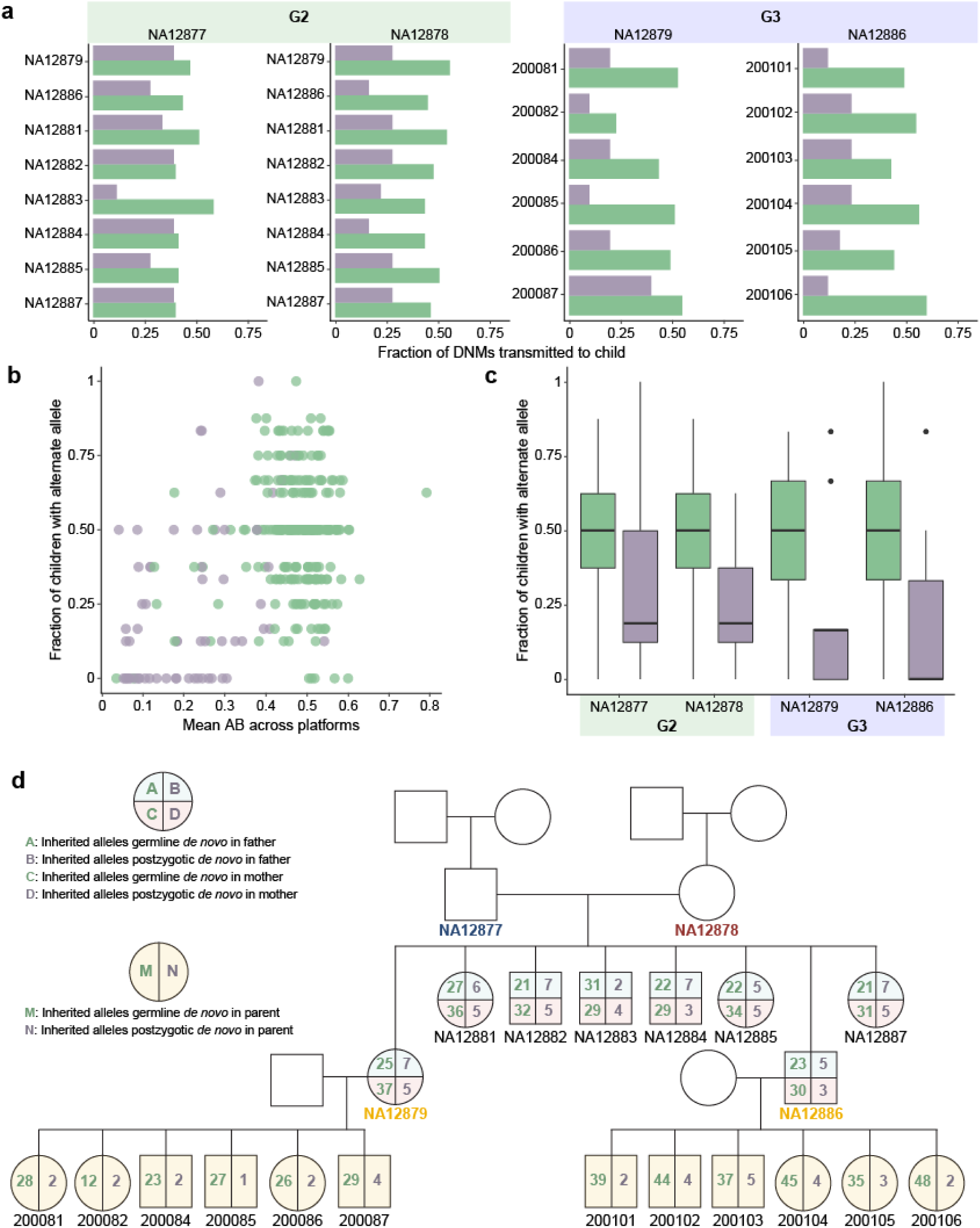
Number of germline and postzygotic SNVs transmitted to children. **a**) The fraction of a parent’s germline SNVs (green, DNMs) and postzygotic SNVs (purple, PZMs) transferred to each child. **b**) The mean allele balance (AB) of DNMs and PZMs across HiFi, Illumina, and ONT data plotted against the fraction of children who inherited a variant reveals that about half the PZMs with AB < 0.25 get transmitted to at least one child. **c**) On average, DNMs are transmitted to 50% of children, while PZMs are transmitted to less than 25% of children. **d**) Number of DNMs and PZMs transmitted to each child in the pedigree.

### *De novo* tandem repeats and recurrent mutation

Given the challenges associated with assaying mutations in short tandem repeats (STRs, 1-6 bp motifs) and variable number of tandem repeats (VNTRs, >6 bp motifs), we applied a targeted HiFi genotyping strategy coupled with validation by transmission and orthogonal sequencing. First, we identified 7.82 million TR loci in the T2T-CHM13 reference genome ranging from 10-10,000 bp (**Methods**). We performed TR genotyping at these loci on HiFi data using the Tandem Repeat Genotyping Tool (TRGT)^28^, across all members of the pedigree. We were able to genotype 7.68 million of these loci in every member of the pedigree and, of those, 7.17 million (93.4%) loci were completely Mendelian concordant across all trios (**Methods**). We investigated all TRGT calls to identify loci at which we could confidently call DNMs by using TRGT-denovo^29^ to annotate them with additional read evidence and applied custom filters (**Methods**). On average, 7.58 million loci were covered by at least 10 HiFi reads in all members of a trio, our threshold to be examined for evidence of TR DNMs.

We refined these putative DNMs through orthogonal sequencing and transmission. Element sequencing exhibits substantially lower error rates following homopolymer tracts^30^, so we tested if it could more accurately measure the length of homopolymers and other TR alleles. We generated Element sequencing from blood DNA for all family members. We observed low “stutter” in the Element data at homopolymers; across a random sample of 1,000 homozygous homopolymer loci called by TRGT, an average of 99.5% of Element reads perfectly support the TRGT-genotyped allele size in GRCh38, compared to 93.5% of Illumina sequencing reads (**Supplementary Figs. 35 and 36**).

We used the Element data to further validate *de novo* TR alleles called by TRGT-denovo. As Element reads are 150 bp in length, we only validated DNMs at STRs with TRGT allele lengths less than 120 bp and spanned by at least 10 Element reads in all members of a trio; using these criteria, we were able to assess 90/671 (13.4%) of *de novo* alleles comprising STRs (average of 11.3 STRs per sample). We considered a DNM validated if Element reads supported the TRGT allele size in the child and did not support it in either parent (allowing for off-by-one base-pair errors, **Methods**). Of the 90 *de novo* STRs we could assess using Element sequencing reads, 56 (62.2%) passed our strict consistency criteria. The validation rate was lower at homopolymers (5/20; 25%) than at non-homopolymers (51/70; 72.9%), indicating that homopolymers still pose a challenge for long-read genotyping, and that our estimates of mutation rates at these loci may be less precise. TR DNMs that failed consistency analysis are significantly shorter than those that passed (Mann Whitney U-test for TR allele length change: p = 1.84 x 10^−10^) and are enriched for *de novo* expansions and contractions of 1 bp; TRGT is known to exhibit higher off-by-one genotyping error rates^28^. We leveraged additional information from the 1463 pedigree to further refine our DNM rate estimates. We required that candidate *de novo* TR alleles observed in the two G3 individuals with sequenced children (NA12879 and NA12886) be transmitted to at least one child in the subsequent generation (G4). Of the 189 *de novo* TR alleles observed in the two G3 individuals, 144 (76.2%) were transmitted to the next generation.

After Element and transmission validation, we found an average of 79.6 TR DNMs (including STRs, VNTRs, and complex loci) per sample and estimated a TR mutation rate across all passing TR loci genome-wide of 5.25×10^−6^ per locus per haplotype per generation (95% CI = 4.42 - 6.01×10^−6^), with substantial variation across repeat motif sizes (**Fig. 3a**). Collectively, TR DNMs inserted or deleted a mean of 1427.1 bp per sample or 15.0 bp per event (**Supplementary Table 11**). An average of 62.3 mutations were expansions or contractions of STR motifs, 7.4 affected VNTR motifs, and 10.0 affected “complex” loci comprising both STR and VNTR motifs. The STR (1-6 bp motif) mutation rate was 5.45×10^−6^ *de novo* events per locus per haplotype per generation (95% CI = 5.0 – 5.95×10^−6^). The VNTR (7+ bp motif) mutation rate was 2×10^−6^ (95% CI = 1.8 – 3.1×10^−6^), predominantly comprising loci that could not be assessed in short-read studies. Several prior estimates of the genome-wide STR mutation rate only considered polymorphic STR loci; when we limited our analysis to STR loci that were polymorphic in the 1463 pedigree, we found 4.01×10^−5^ *de novo* events per locus per generation (95% CI = 3.49 – 4.58×10^−5^), which is broadly consistent with prior estimates of 4.95 – 5.6×10^−5 31–33^. TR DNMs were more common in the paternal germline; 73.9% of phased *de novo* TR alleles were paternal in origin (**Fig. 3b**). The mutation rate for dinucleotide motifs was higher than for homopolymers, and we observed increasing mutation rate with motif size for motifs greater than 6 bp in length (**Fig. 3a**).

**Figure 3.**
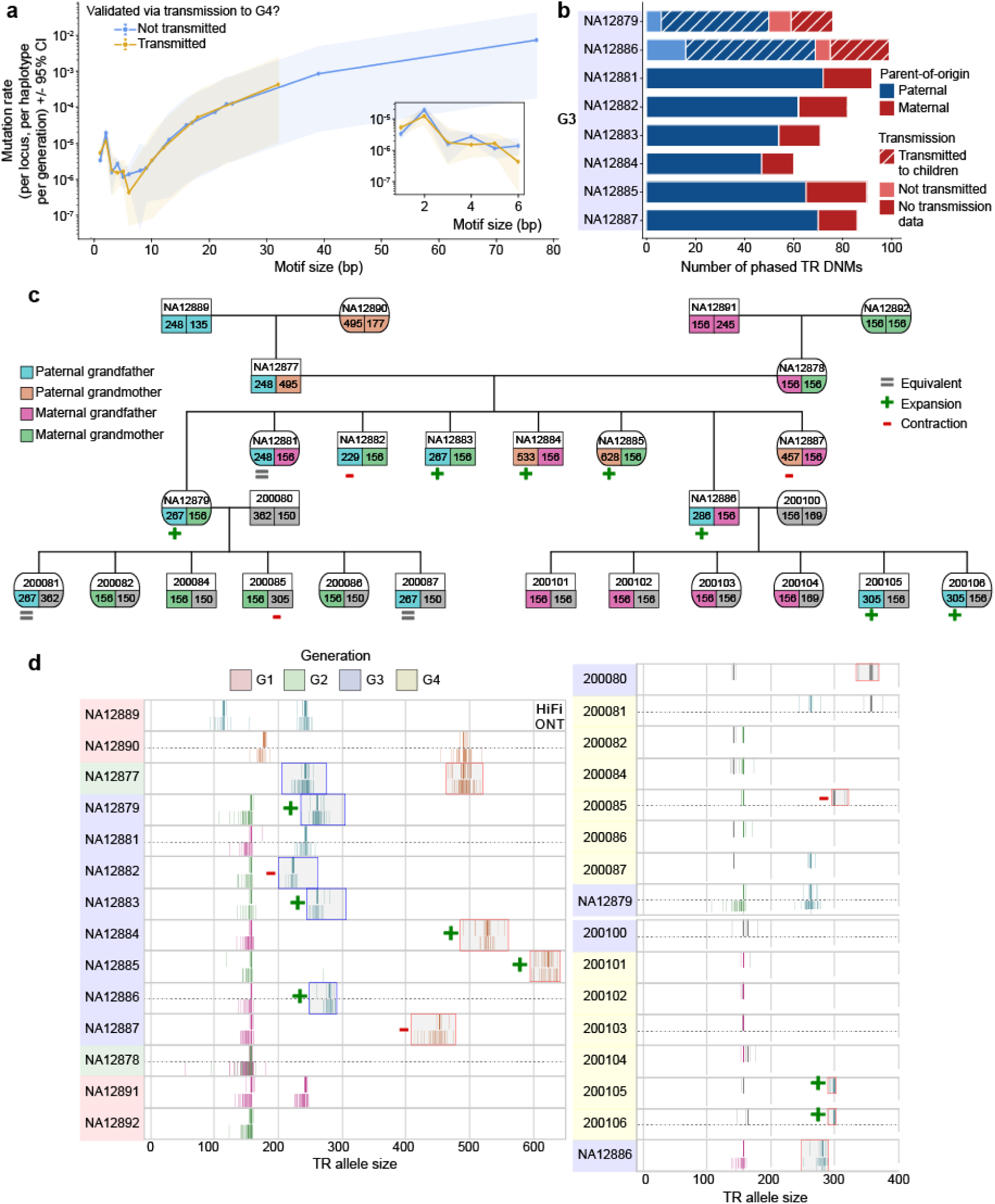
Tandem repeat *de novo* mutations (TR DNMs) show motif size dependent mutation rates, paternal bias, and are highly recurrent at specific loci. **a**) TR DNM rates (mutations per haplotype, per locus, per generation) as a function of TR motif size in the T2T-CHM13 reference genome. Complex TR loci that comprise more than one unique motif were excluded. Error bars denote 95% Poisson confidence intervals around the mean mutation rate estimate. Mutation rates include all calls that pass TRGT-denovo filtering criteria but are not adjusted for Element validation. **b**) Inferred parent-of-origin for confidently phased TR DNMs in G3. Crosshatches indicate transmission to at least one G4 child, where available. **c**) Pedigree overview of a recurrent VNTR locus at chr8:2376919-2377075 (T2T-CHM13) with motif composition GAGGCGCCAGGAGAGAGCGCT(n)ACGGG(n). Allele coloring indicates inheritance patterns as determined by inheritance vectors, gray representing unavailable data. Symbols denote inheritance type relative to the inherited parental allele: “+” for de novo expansion, “-” for de novo contraction, and “=” for regular inheritance, shown only for the mutating alleles, and numbers indicate allele lengths in bp. **d**) Read-level evidence for the recurrent DNM in (c), represented as vertical lines, obtained from individual sequencing reads, shown per sample. Where available, both HiFi (top) and ONT (bottom) sequencing reads are displayed. Coloring is consistent with inheritance patterns in (c); outlined boxes with +/- markers highlight DNMs.

We identified a subset of TR loci that were recurrently mutated amongst members of the 1463 pedigree. After both strict filtering and visual inspection of reads (**Methods**), we identified a high-confidence set of 32 loci (**Supplementary Table 12**): five showing intragenerational recurrence (observed DNMs in at least two G3 individuals) and 27 loci with intergenerational recurrence (observed DNMs in at least two generations). Notably, we observed three or more unique *de novo* expansions or contractions at 16 of the loci that exhibited recurrence (**Table 1**). As an example, we highlight an intergenerational recurrently mutated TR locus with 10 unique *de novo* expansions and/or contractions (**Fig. 3c,d**). *De novo* TR alleles are present at this locus in seven of eight G3 individuals; these *de novo* alleles transmit to four G4 individuals, with two expanding further upon transmission. Additionally, the spouse of a G3 individual (sample 200080) carries a distinct TR allele that undergoes a *de novo* contraction in subsequent transmissions. This recurrent *de novo* was supported by both HiFi and ONT reads.

**Table 1.**
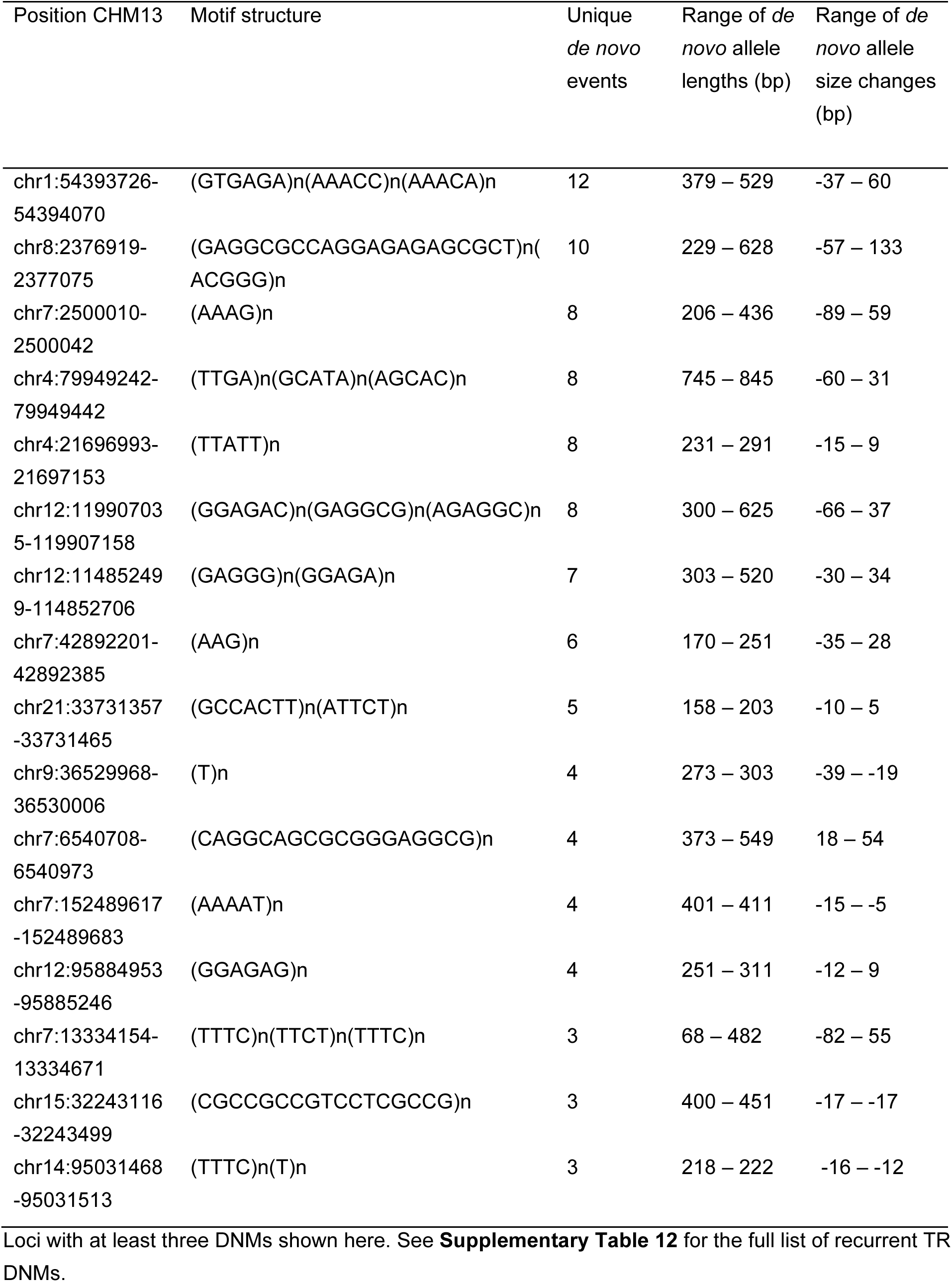
Recurrently mutated tandem repeat loci.

### Centromere familial transmission and *de novo* SVs

Among the 288 completely sequenced and assembled centromeres, we were able to assess 150 transmissions (33 from G1 to G2 and another 117 transmissions from G2 to G3) (**Fig. 4a**). Comparing these assembled centromeres between parent and child (**Methods**), we identify 18 (12%) *de novo* SVs validated by both ONT and HiFi data with roughly equivalent number of insertions and deletions (**Fig. 4b**). We find that 72.2% (13/18) of the structural changes map to ɑ-satellite higher order repeat (HOR) arrays with the remainder (5/18 or 27.7%) corresponding to various pericentromeric flanking sequences but none within flanking monomeric alpha satellite. All ɑ-satellite HOR *de novo* SV events involve integer changes of the basic ɑ-satellite HOR cassettes specific to each centromere and range in size from 680 bp (one 4-mer ɑ-satellite HOR on chr9) to 12,228 bp (4×18-mer ɑ-satellite HORs on chr6) (**Fig. 4c, Extended Data Fig. 4a**). One transmission from chromosome 9 involves both a gain of 2,052 bp (six dimer ɑ-satellite HOR units) and a loss of 1,710 bp (one 4-mer ɑ-satellite HOR and three ɑ-satellite dimer units) in a single G2 to G3 transmission (**Fig. 4d-f**). The chromosome 6 centromere harbors the most recurrent structural events with three being observed across three generations (**Fig. 4a**). This enrichment of centromeric events on chromosome 6 is notable, as it is also the centromere that has the greatest number of nearly perfectly identical (>99.9%) ɑ-satellite HORs (**Extended Data Fig. 4b**). Although the data are still preliminary, we also assess 18 SV events for their potential effect on the hypomethylation pocket associated with the centromere dip region (CDR)—a marker of the site of kinetochore attachment^34,35^. We find that 11 SVs mapping outside of the CDR have a marginal effect on changing the center point of the CDR (<100 kbp) from one generation to another (**Extended Data Fig. 4c,d**), while SVs mapping within the CDR have a more dramatic effect (average shift ∼260 kbp) and/or they completely alter the distribution of the CDR (**Fig. 4g, Extended Data Fig. 4e,f**). Although follow-up experiments using CENP-A ChIP-seq are needed to confirm the actual binding site of the kinetochore, these findings suggest that structural mutations may have epigenetic consequences in changing the position of kinetochore on at least three occasions in this family. Finally, using 31 parent–child transmissions of centromeres (150.5 Mbp), we used the assemblies to reassess the SNV DNMs. We identify 48 SNV DNMs within the ɑ-satellite HOR DNA, suggesting a significantly higher rate of DNM at 7.4×10^−7^ mutations/bp/generation (95% C.I. = 0 - 1.18×10^−6^) when compared to the 4.21×10^−8^ (95% C.I. = 1.98×10^−8^ - 6.44×10^−8^) rate calculated from 18 centromeric SNVs identified from read-based mapping (**Fig. 2a, Supplementary Table 11**).

**Figure 4.**
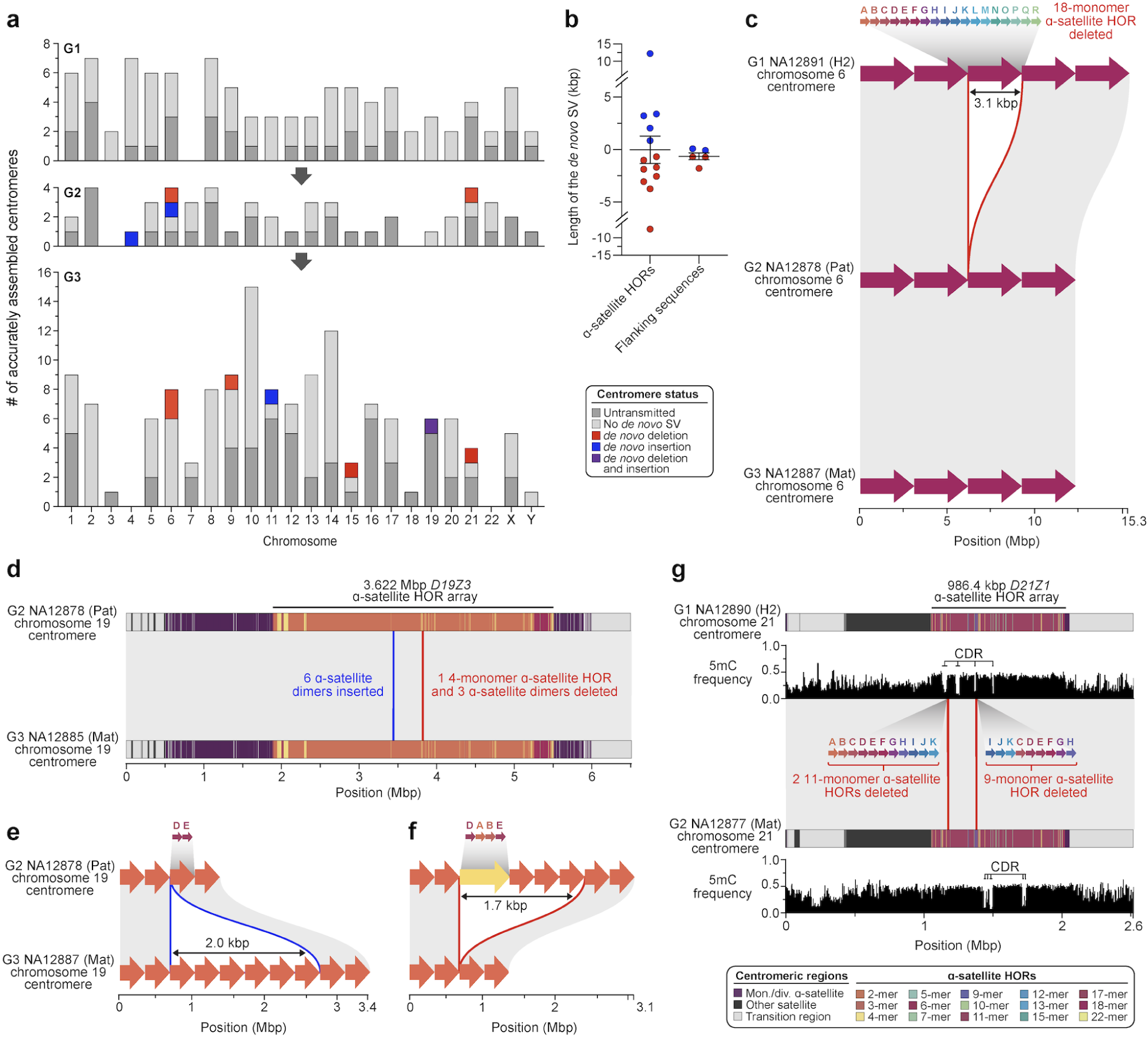
*De novo* SVs among centromeres transmitted across generations. **a**) Plot summarizing the number of correctly assembled centromeres (dark gray) as well as those transmitted to the next generation (light gray). Transmitted centromeres that carry a *de novo* deletion, insertion, or both are colored (see legend). **b**) Lengths of the *de novo* SVs within α-satellite HOR arrays and flanking regions.**c)** An example of a *de novo* deletion in the chromosome 6 α-satellite HOR array in G2-NA12878 that was inherited in G3-NA12887. Red arrows over each haplotype show the α-satellite HOR structure, while gray blocks between haplotypes show syntenic regions. The deleted region is highlighted by a red outline. **d)** An example of a *de novo* insertion and deletion in the chromosome 19 α-satellite HOR array of G3-NA12887. **e-f**) Zoom-in of the α-satellite HOR structure of the inserted (blue outline) and deleted (red outline) α-satellite HORs from (d). Again, colored arrows on top of each haplotype show the α-satellite HOR structure. **g**) Example of two *de novo* deletions in the chromosome 21 centromere of G2-NA12877. The deletions reside within a hypomethylated region of the centromeric α-satellite HOR array, known as the “centromere dip region” (CDR), which is thought to be the site of kinetochore assembly. The deletion of three α-satellite HORs within the CDR results in a shift of the CDR by ∼260 kbp in G2-NA12877.

**Extended Data Figure 4.**
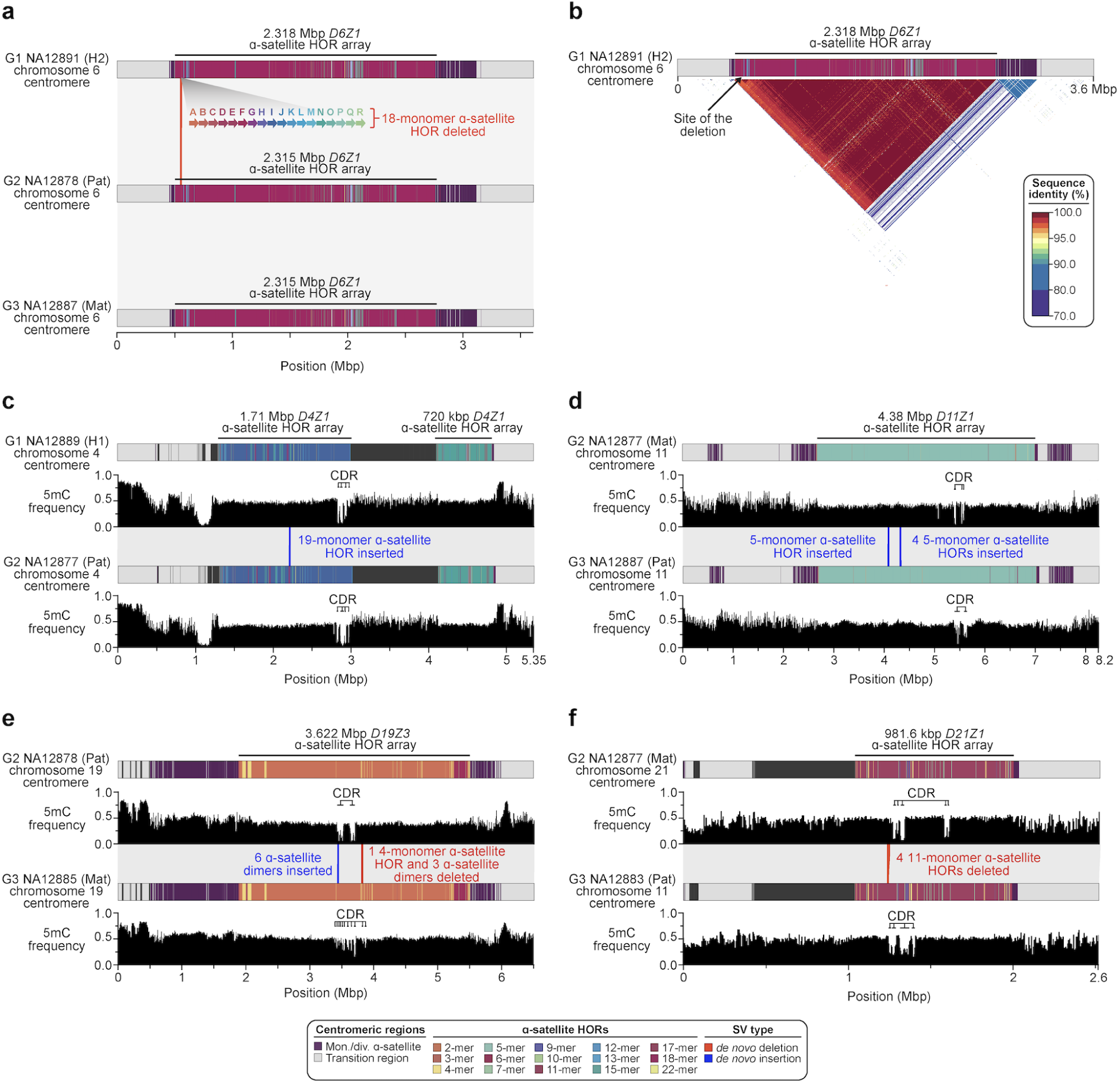
Changes in centromere sequence, structure, and DNA methylation patterns across generations. **a**) Deletion of an 18-monomer α-satellite HOR within the chromosome 6 centromere of G2-NA12878 is inherited in G3-NA12887, shortening the length of the α-satellite HOR array by ∼3 kbp. **b**) Sequence identity heatmap of the chromosome 6 centromere in G1-NA128991 shows the high (∼100%) sequence identity of α-satellite HORs along the entire centromeric array and at the site of the *de novo* deletion. **c,d**) Deletions of α-satellite HORs in regions outside of the centromere dip region (CDR) in the **c**) chromosome 4 and **d**) chromosome 11 centromeres does not affect the position of the CDR. **e,f**) Deletions and insertions of α-satellite HORs within the CDR in the **e**) chromosome 19 and **f**) chromosome 21 centromeres alter the distribution of the CDR.

### Y chromosome mutations

Here, we focus on the ∼59.7 Mbp male-specific Y-chromosomal region (MSY, i.e., excluding pseudoautosomal regions) considering both read-based as well as assembly-based approaches to discover DNMs (**Methods, Supplementary Notes**). There are nine male members who carry the R1b1a-Z302 Y haplogroup across the four generations (**Fig. 5a, Supplementary Table 13**) and we use the great-grandfather (G1-NA12889, **Fig. 1**) chromosome Y assembly as a reference for DNM detection across the 48.8 Mbp MSY. The *de novo* assembly-based approach increases by >2-fold the number of accessible base pairs when compared to HiFi read-based calling but increases by >7-fold the discovery of *de novo* SNVs (**Methods**). In total, we identify 48 *de novo* SNVs in the MSY across the five G2-G3 males, ranging from 7-11 SNVs per Y transmission (mean 9.6, median 10) (**Supplementary Table 14**). Only two SNVs map to the Y euchromatic regions, one to the pericentromeric with the remaining 45/48 to the Yq12 heterochromatic satellite regions (**Fig. 5b**). We thus estimate the *de novo* SNV rate of 1.99×10^−7^ (95% CI = 1.59 - 2.39×10^−7^) for the MSY combining both read- and assembly-based approaches. It is important to note that 13/45 (29%) of the DNMs had 100% identical matches elsewhere in the Yq12 region (but not at orthologous positions) and could, therefore, result from interlocus gene conversion events (**Methods**) consistent with the *DYZ1/HSat3A6* and *DYZ2/HSat1B* organization of the region^36^. We also identify a total of nine *de novo* indels (<50 bp, homopolymers excluded) ranging from 1-3 indels/sample (mean 1.8 events/Y transmission) and five *de novo* SVs (≥50 bp) (**Fig. 5b, Supplementary Table 14**). The latter range from 2,416 to 4,839 bp in size, each affecting an entire *DYZ2* repeat unit(s), with an average of one SV per Y transmission. Variants detected in the G3 parents of G4 are confirmed by both transmission and read data, supporting the high quality of the variant calls. Overall, 83% (52/63) of the DNMs identified on chrY (42/48 SNVs, 4/9 of indels and 5/5 SVs) are located in regions where short reads cannot be reliably mapped (mapping quality = 0).

**Figure 5.**
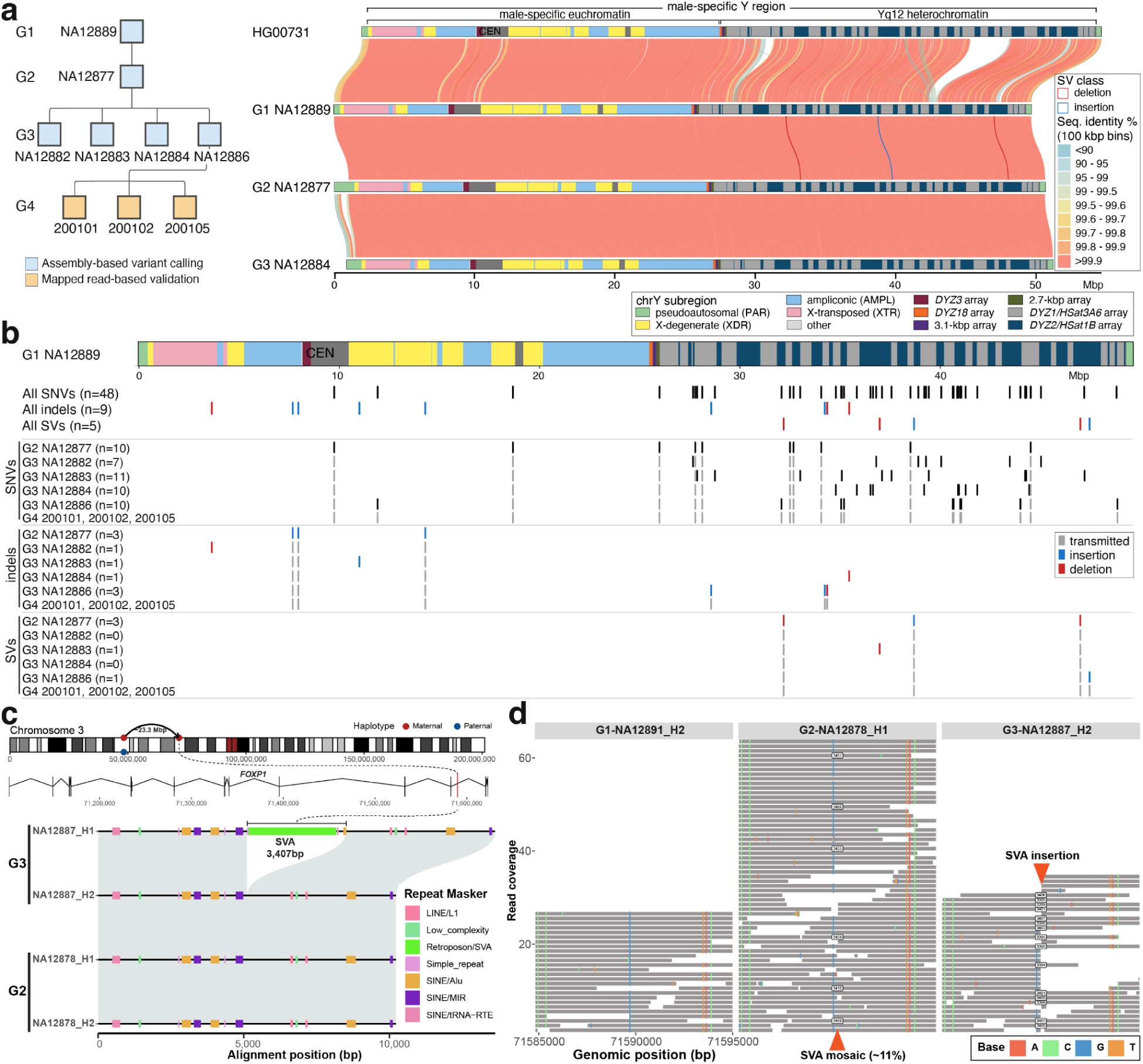
Chromosome Y and an example of a *de novo* mobile element. **a**) Pedigree of the nine males carrying the R1b1a-Z302 Y chromosomes (left) and pairwise comparison of Y assemblies: closely related Y from HG00731 (R1b1a-Z225) and the most contiguous R1b1a-Z302 Y assemblies from three generations. Y-chromosomal sequence classes are shown with pairwise sequence identity between samples in 100 kbp bins, with QC-passed SVs identified in the pedigree males shown. **b**) Summary of chrY DNMs. Top - Y structure of G1-NA12889. Below the Y structure - all identified DNMs across G1-G3 Y assemblies. Bottom - breakout by mutation class and by sample. In light gray are DNMs that show evidence of transmission from G2 to G3-G4, and from G3-NA12886 to his male descendants in G4. **c**) *De novo* SVA insertion in G3-NA12887. **d**) HiFi read support for the *de novo* SVA insertion in G3-NA12887.

### *de novo* SVs

Using the operational definition of (≥50 bp) and accumulating across the above analyses, we validate a total of 41 *de novo* SVs across eight individuals (G3), including 16 insertions and 25 deletions. This set of *de novo* SVs was vetted by visual inspection of read evidence and assembly support and as such likely represents a lower bound. Overall, 68% (28/41) of events originate in the paternal germline with evidence of an increase in SVs with paternal age (**Supplementary Fig. 37**). Almost all of the validated SVs (40/41) correspond to contractions and expansions of TRs described above, including mutation in centromeres, Y chromosome satellites, and clustered SDs. We report two TR events subject to recurrent mutations (**Supplementary Table 11**). We estimate ∼5 SVs (95% CI: 3-7) per transmission affecting approximately ∼4.4 kbp of DNA (median: 4875 bp). If we exclude *de novo* SVs mapping to the centromere and Y chromosomes (n=14), the median size of the events drops by an order of magnitude (median: 362 bp). Non-allelic homologous recombination (NAHR) has frequently been invoked as a mechanism to underlie TR expansions and contractions^37,38^. We find, however, that none of the 27 euchromatic *de novo* SVs coincide with detected crossover positions (**Supplementary Fig. 37d**). In fact, the average distance was often many megabase pairs apart arguing against NAHR as the primary mechanism for their origin. Although most *de novo* SVs involve TR changes, we identify one exception: a full-length (3,407 bp long) *de novo* insertion of an SVA element (SVAF subfamily) in sample G3-NA12887^39^. We define target site duplication to be “GATTATGTTTC” and the length of the poly-A tail to be 43 bp long. We predict the donor site of this element to be on the same homolog ∼23 Mbp upstream from the insertion site (**Fig. 5c, Supplementary Fig. 38**). We also find this insertion present at a low frequency (∼11% of overlapping reads) in the parent (G2-NA12878) but not in the grandparental transmitting haplotype consistent with a germline mosaic event arising in G2 (**Fig. 5d, Supplementary Fig. 39**).

## DISCUSSION

The origin, rate, and distribution of DNMs is arguably one of the most important aspects of human genetics and key to our understanding of genetic disease, phenotypic variation, and the evolution of our species^40^. Most studies^41–45^ that establish DNM rates utilize short reads amongst large groups of trios and generally agree on 60-70 DNMs per generation; however, this largely excludes highly mutable regions of the genome, e.g., long TRs, SDs, and satellite sequences^7^. Our approach differs in that we generated a comprehensive multi-generational assembly-based resource with five orthogonal short- and long-read sequencing technologies with the aim to catalog transmitting and *de novo* variation of all classes—establishing a truth set for human genetic variation and all subsequent sequencing technologies. The multiplatform and multigenerational, assembly-based approach provides access to some of the most difficult regions of the genome, such as centromeres and heterochromatic regions on the Y chromosome. The use of parental references in addition to the standard GRCh38 and T2T-CHM13 references and the ability to confirm transmissions across subsequent generations improves both sensitivity and specificity. In this multigenerational pedigree, we estimate 128-259 DNMs per generation, including 75.5 *de novo* SNVs and 7.4 non-TR indels, indels or SVs originating from TRs (79.6 *de novo*), centromeric changes (7.7 *de novo* events per generation), and the Y chromosome (12.4 *de novo* events per generation). We observe a strong paternal *de novo* bias (70-80%) and an increase with advancing paternal age not only for SNVs but also for indels and SVs, including TRs.

We find that the rate of *de novo* SNVs varies by more than an order of magnitude depending on the genomic context. In particular, we observe elevated rates of *de novo* SNVs in repetitive regions both for germline and postzygotic events, consistent with recent human population-based analyses^7,46^ and theoretical predictions^47^. SD regions show an 88% increase (2.2 vs. 1.17×10^−8^), which is driven by SDs with >95% identity. Although the number of validated SNV DNMs is still rather modest, we currently estimate that satellite DNA constituting centromeres and the Yq12 heterochromatic region is at least 30 times more mutable (3.68×10^−7^ - 7.41×10^−7^ mutations per bp per generation) than autosomal euchromatin. The Yq12 region in particular has never been studied at base-pair resolution as it is largely missing from the GRCh38 reference and its complete assembly has only recently become possible^36,48^. It is composed of thousands of short satellite DNA repeats (*DYZ1/Hsat3A6* and *DYZ2/Hsat1B*) organized into blocks that are >98% identical, in total tens of megabase pairs in size^36,48^. This, along with the fact that 29% of mutational changes match to non-orthologous sites in Yq12, is consistent with “interlocus gene conversion” driving this excess potentially as a result of increased sister chromatid exchange events^36^. While our DNM SNV rate estimate for Y euchromatic regions is comparable to previous pedigree-based work (∼22 Mbp, 1.81×10^−8^ mutations per bp per generation in this study compared to 2.87×10^−8^ mutations per bp per generation from^43^), the SNV estimate for Yq12 is >20x higher.

In addition to germline events, we classified nearly twice the number of *de novo* SNVs as PZMs (12.9 PZM per transmission or 17%) compared to even the highest previous estimate (6-10%)^14,49^. Previous studies have distinguished between *de novo* post-zygotic and germline SNVs using allele balance thresholds^49^ or by identifying incomplete linkage to nearby SNVs across three generations^14^. Long-read sequencing provides a third approach, allowing us to assign nearly every *de novo* SNV to a parental haplotype and distinguish mosaic events by the presence of three distinct long-range haplotypes. Early cell divisions of human embryos are frequently error prone^50^ with an accelerated rate of cell division and these properties may contribute to the high fraction of PZMs with high (>25%) allele balance (38% are estimated to have high allele balance and 83% of these (n=20/24) are transmitted to the next generation). Such events would previously have been classified as germline but, consistent with PZM expectations, we find no paternal bias associated with these *de novo* variants (**Fig. 2b**).

As has long been observed^32,51,52^, TRs are among the most mutable loci of our genome with the number of such *de novo* events comparable to germline SNVs^53^ but affecting more than an order of magnitude more base pairs per generation. Unlike previous studies^31,32^, long-read sequencing and assembly allows the sequence characterization of *de novo* events many kilobase pairs in length and in regions where it is difficult to map short reads. We find a threefold differential in TR DNM rate with increasing repeat number and motif length generally correlating with mutation rate. We, however, observe an apparent mutation rate “trough” between dinucleotides and larger motif lengths (>10 bp) (**Fig. 3b**), which may reflect different mutational mechanisms based on locus size, motif length, and complexity. For example, larger TR motifs may be more likely to mutate via NAHR, synthesis-dependent strand annealing, or interlocus gene conversion while mutational events at STRs may be biased toward traditional replication-based slippage mutational mechanisms^37,38^. Consistent with some earlier genome-wide analyses of minisatellites^54^, we did not find evidence that TR changes are mediated by unequal crossover during meiosis since none of our TR DNMs coincided with recombination breakpoints. Of particular interest, in this regard, is the discovery of 32 recurrently mutated TRs—loci rarely discovered out of the context of unstable disease alleles^55^. At five of these recurrent loci, we discovered multiple DNMs within a single generation (G3); these DNMs may be the outcome of germline mosaicism in a G2 parent or the activity of hypermutable TRs. The remaining loci are recurrently mutated in multiple generations, and likely represent a collection of highly mutable TR loci. Nearly all of these highly recurrent *de novo* events produced TR alleles that are significantly longer than the average short-read length and were only detectable using long-read sequencing. This includes changes in the length of ∼7% of human centromeres where insertions and deletions all occur as multiples of the predominant HOR unit^51^. Not surprisingly, the rate of *de novo* SVs increased from previous estimates of 0.2-0.3 per generation^15,56^ to 3-4 *de novo* SVs per generation reported in this study.

There are several limitations to this study. First, as discussed, homopolymers still remain challenging even with the use of Element data since longer alleles and motifs embedded in larger repeats are still not reliably assayed with short reads. Second, we were unable to successfully characterize *de novo* variation in the acrocentric regions due to the repetitive nature of the regions and rampant heterologous recombination. An important next step will be to assign acrocentric contigs to their respective chromosomes and assess patterns of mutation and ectopic recombination in regions predicted to be the most dynamic^4^. Third, we examined only one multigenerational family and familial variation is expected depending on the genetic background^14,32,57^. Many more families will be required to establish a reliable estimate of the mutation rate especially for complex regions of the genome. In that regard, it is perhaps noteworthy that efforts are underway to characterize an additional 10 CEPH pedigrees. Notwithstanding, this study highlights two facts that a single sequencing technology and a single human genome reference are insufficient to comprehensively estimate mutation rates. This is especially problematic in complex regions such as centromeres and heterochromatic regions of chromosome Y where assembly-based parent-to-offspring comparisons are required to catalog DNMs. More variation, including *de novo* variation, remains to be discovered—we were conservative in our callset requiring multiple technologies supporting the discovery of DNM, assessing transmission, where possible, to the next generation for all variants, and being careful to consider DNA from primary tissue as opposed to cell lines. It is noteworthy that several studies with long-read sequencing technologies have claimed higher DNM rates^10,58^. The multigenerational resource we generated will further refine these estimates and serve as a useful benchmark for new algorithms and new sequencing technologies similar to GIAB^59^.

## Data availability

All underlying sequence data from 28 members of the family will be available in dbGaP under accessions numbers: TBD – in process. Importantly, 17 family members (G1-NA12889, G1-NA12890, G1-NA12891, G1-NA12892, G2-NA12877, G2-NA12878, G3-NA12879, G3-NA12881, G3-NA12882, G3-NA12885, G3-NA12886, G4-200080, G4-200081, G4-200082, G4-200085, G4-200086, G4-200087) consented for their data to be publicly accessible similar to the 1000 Genomes Project samples to allow for development of new technologies, study of human variation, research on the biology of DNA, and study of health and disease.

Corresponding data and phased genome assemblies will be made available via the AWS Open Data program: s3://palladium-genomes.pacbcloud.com/DataSharing/TBD The Y-chromosomal assembly for a closely related R1b haplogroup sample HG00731 was downloaded from the Human Genome Structural Variation Consortium IGSR site (https://ftp.1000genomes.ebi.ac.uk/vol1/ftp/data_collections/HGSVC3/working/2023092 7_verkko_batch2/assemblies/HG00731/).

Reference genomes and their annotations used in this study are listed in **Supplementary Table 15.**

## Code availability

Custom code and pipelines used in this study are publicly available via the following GitHub repositories:

Workflows and custom code: https://github.com/orgs/Platinum-Pedigree-Consortium/repositories
TRGT: https://github.com/PacificBiosciences/trgt
TRGT-denovo: https://github.com/PacificBiosciences/trgt-denovo
SVbyEye: https://github.com/daewoooo/SVbyEye

## Acknowledgements

We thank Sonata Jankauskiene, Mianne Lee, and Trang Nguyen for technical assistance with preparation and sequencing of Strand-seq libraries; Christian Steidl for use of the NextSeq550 sequence platform; and Tonia Brown for useful edits in the preparation of this manuscript. Library pools were also sequenced on the Element AVITI at the University of California Davis DNA Technologies Core.

The following cell lines were obtained from the NIGMS Human Genetic Cell Repository at the Coriell Institute for Medical Research: GM12877, GM12878, GM12879, GM12881, GM12882, GM12883, GM12884, GM12885, GM12886, GM12887, GM12889, GM12890, GM12891, GM12892.

This research was supported, in part, by funding from the National Institutes of Health (NIH) grants R01 HG002385 and R01 HG010169 (to E.E.E.) and U24 HG007497 (to E.E.E. and C.Lee). E.E.E. is an investigator of the Howard Hughes Medical Institute. The Strand-seq work was funded in part by a program project grant (#1074) from the Terry Fox Research Foundation and a research grant (#159787) from the Canadian Institutes of Health Research.

This research was further supported by funding to H.D. by 5K99HG012796-02, C.J.S. by R00HG011657, and L.B.J. and W.S.W. being supported by NIH R35GM118335. G.A.L. was supported by NIH GM147352.

This article is subject to HHMI’s Open Access to Publications policy. HHMI lab heads have previously granted a nonexclusive CC BY 4.0 license to the public and a sublicensable license to HHMI in their research articles. Pursuant to those licenses, the author-accepted manuscript of this article can be made freely available under a CC BY 4.0 license immediately upon publication.

## Author contributions

D.P., L.B.J. and E.E.E. conceptualization.

P.H., P.E. and C.Lee. Chromosome Y analysis.

M.D.N., D.P., H.D., T.A.S., P.H., G.A.L. and Z.N.K. DNM analysis.

N.K., W.T.H. and D.P. Generation of *de novo* assemblies and validation.

N.K., W.T.H., W.J.R., J.L., T.Y.L., V.C.T.H. and H.J. Data analysis support.

D.P., K.K.O. and G.A.L. Centromere analysis.

K.G. and C.E.M. Telomere analysis.

D.P., C.N., Z.N.K. and A.G. Meiotic recombination analysis.

H.D., T.A.S., T.M., E.D., T.J.N., M.E.G. and A.R.Q. TR analysis.

C.J.S. and W.S.W. MEI analysis.

K.M.M., K.H., D.D.C., Y.W., J.K., G.H.G., C.F. and C.Lambert. Generated sequencing data.

B.S.P., H.C.H., S.M. and J.D.S. Short-read callsets generation.

D.P., H.D., T.A.S., T.M., G.A.L., P.H., M.D.N. and E.E.E. developed main figures.

D.P., H.D., T.A.S., G.A.L., P.H., M.D.N., Z.N.K. and E.E.E. manuscript writing.

S.L., C.E.M., E.G., P.M.L., D.W.N., L.B.J., A.R.Q., M.A.E. and E.E.E. supervised experiments and analyses.

## METHODS

### Sample and DNA preparation

Family members from G2 and G3 were re-engaged for the purpose of updating informed consent and health history, and for enrolling their children (G4) and the marry-in parent (G3). Archived DNA from G2 and G3 was extracted from whole blood. Newly enrolled family members underwent informed consent and blood was obtained for DNA and cell lines. DNA was extracted from whole blood using the Flexigene system (Qiagen 51206). All samples are broadly consented for scientific purposes, which makes this dataset ideal for future tool development and benchmarking studies.

### Sequence data generation

Sequencing data from orthogonal short- and long-read platforms were generated as follows:

### Illumina data generation

Illumina WGS on G1-G3 was generated as previously described^14^. Illumina WGS on G4 and marry-in spouses for G3 were generated by the Northwest Genomics Center using the TruSeq library prep kit and sequenced to approximately 30x on the NovaSeq 6000 with paired end 150 bp reads.

### PacBio HiFi sequencing

PacBio HiFi data were generated per manufacturer’s recommendations. Briefly, DNA was extracted from blood samples as described or cultured lymphoblasts using the Monarch® HMW DNA Extraction Kit for Cells & Blood (New England Biolabs, T3050L). At all steps, quantification was performed with Qubit dsDNA HS (ThermoFisher, Q32854) measured on DS-11 FX (Denovix) and size distribution checked using FEMTO Pulse (Agilent, M5330AA & FP-1002-0275.) HMW DNA was sheared with Megaruptor 3 (Diagenode, B06010003 & E07010003) using settings 28/30, 28/31, or 27/29 based on initial quality check to target a peak size of ∼22 kbp. After shearing, the DNA was used to generate PacBio HiFi libraries via the SMRTbell prep kit 3.0 (PacBio, 102-182-700). Size selection was performed either with diluted AMPure PB beads per the protocol, or with Pippin HT using a high-pass cutoff between 10-17 kbp based on shear size (Sage Science, HTP0001 & HPE7510). Libraries were sequenced either on the Sequel II platform on SMRT Cells 8M (PacBio, 101-389-001) using Sequel II sequencing chemistry 3.2 (PacBio,102-333-300) with 2-hour pre-extension and 30-hour movies, or on the Revio platform on Revio SMRT Cells (PacBio, 102-202-200) and Revio polymerase kit v1 (PacBio, 102-817-600) with 2-hour pre-extension and 24-hour movies.

### ONT sequencing

To generate UL sequencing reads >100 kbp, we used ONT sequencing. Ultra-high molecular weight gDNA was extracted from the lymphoblastoid cell lines according to a previously published protocol^60^. Briefly, 3-5 x 10^7 cells were lysed in a buffer containing 10 mM Tris-Cl (pH 8.0), 0.1 M EDTA (pH 8.0), 0.5% w/v SDS, and 20mg/mL RNase A for 1 hour at 37°C. 200 ug/mL Proteinase K was added, and the solution was incubated at 50°C for 2 hours. DNA was purified via two rounds of 25:24:1 phenol-chloroform-isoamyl alcohol extraction followed by ethanol precipitation. Precipitated DNA was solubilized in 10 mM Tris (pH 8.0) containing 0.02% Triton X-100 at 4°C for two days.

Libraries were constructed using the Ultra-Long DNA Sequencing Kit (ONT, SQK-ULK001) with modifications to the manufacturer’s protocol: ∼40 ug of DNA was mixed with FRA enzyme and FDB buffer as described in the protocol and incubated for 5 minutes at RT, followed by a 5-minute heat-inactivation at 75°C. RAP enzyme was mixed with the DNA solution and incubated at RT for 1 hour before the clean-up step. Clean-up was performed using the Nanobind UL Library Prep Kit (Circulomics, NB-900-601-01) and eluted in 450 uL EB. 75 uL of library was loaded onto a primed FLO-PRO002 R9.4.1 flow cell for sequencing on the PromethION, with two nuclease washes and reloads after 24 and 48 hours of sequencing. All G1-G3 ONT base calling was done with guppy (v6.3.7).

### Element (AVITI) sequencing

Element WGS data was generated per manufacturer’s recommendations. Briefly, DNA was extracted from whole blood as described above. PCR-free libraries were prepared using mechanical shearing, yielding ∼350 bp fragments, and the Element Elevate library preparation kit (Element Biosciences, 830-00008). Linear libraries were quantified by qPCR and sequenced on AVITI 2 x 150 bp flow cells (Element Biosciences, not yet commercially available). Bases2Fastq Software (Element Biosciences) was used to generate demultiplexed FASTQ files.

### Strand-seq library preparation

Single-cell Strand-seq libraries were prepared using a streamlined version of the established OP-Strand-seq protocol^61^ with the following modifications. Briefly, EBV cells from G1-3 were cultured for 24 hrs in the presence of BrdU and nuclei with BrdU in the G1 phase of the cell cycle were FACS sorted as described^61^. Next, single nuclei were dispensed into individual wells of an open 72 x 72 well nanowell array and treated with heat-labile protease, followed by digestion of DNA with restriction enzymes AluI and HpyCH4V (NEB, Ipswich, MA) instead of micrococcal nuclease (MNase). Next, fragments were A-tailed, ligated to forked adapters, UV-treated, and PCR-amplified with index primers. The use of restriction enzymes results in short, reproducible, blunt-ended DNA fragments (>90% smaller than 1 kbp) that do not require end-repair before adapter ligation, in contrast to the ends of DNA generated by MNase. Omitting end-repair enzymes allows dispensing of index primers in advance of dispensing individual nuclei. The pre-spotted, dried primers survive (and do not interfere) with library preparation steps prior to PCR amplification. Pre-spotting of index primers is more reliable than the transfer of index primers between arrays during library preparation as described^61^. Strand-seq libraries were pooled, cleaned with AMPure XP beads, and library fragments between 300 and 700 base pairs were gel purified prior to PE75 sequencing on either the NextSeq 550 or the AVITI (Element Biosciences, San Diego, CA). **Supplementary Fig. 40** shows examples of Strand-seq libraries made with restriction enzymes.

### Strand-seq data post-processing

The demultiplexed FASTQ files were aligned to both GRCh38 and T2T-CHM13 reference assemblies (see ‘Reference genomes used in this study’) using BWA^62^ (v0.7.17-r1188) for standard library selection. Aligned reads were sorted by genomic position using SAMtools^63^ (v1.10) and duplicate reads were marked using sambamba^64^ (v1.0). Libraries passing quality filters were pre-selected using ASHLEYS^65^. We also evaluated such selected Strand-seq libraries manually and further excluded libraries with an uneven coverage, or an excess of ‘background reads’ (reads mapped in opposing orientation for chromosomes expected to inherit only Crick or Watson strands) as previously described^66^. This is done to ensure accurate inversion detection and phasing. Finally, we selected 60+ libraries (range: 62-90) per sample with a median ∼274K reads with mapping quality ≥10 per library what translates to about 0.67% genome (T2T-CHM13) being covered per library (**Supplementary Fig. 41**).

### Strand-seq inversion detection

Polymorphic inversions were detected by mapping Strand-seq read orientation with respect to the reference genome as previously described^67,68^. Briefly, we ran breakpointR^69^ (v1.15.1) across selected Strand-seq libraries to detect points of strand-state changes^69^. We used these results to generate sample-specific composite files using breakpointR function ‘synchronizeReadDir’ as described previously^67^. Again, we ran breakpointR on such composite files to detect regions where Strand-seq reads map in reverse orientation and are indicative of an inversion. Lastly, we manually evaluated each reported inverted region by inspection of Strand-seq read mapping in UCSC Genome Browser^70^ and removed any low-confidence calls.

### Generation of phased genome assemblies

Phased genome assemblies were generated using two different algorithms, namely Verkko (v1.3.1 and v1.4.1)^16^ and hifiasm (UL) with ONT support (v0.19.5)^17^. Due to active development of the Verkko and hifiasm algorithms, assemblies were generated with two different versions. Phased assemblies for G2-G3 were generated using a combination of HiFi and ONT reads using parental Illumina *k*-mers for phasing. To generate phased genome assemblies of G1, we still used a combination of HiFi and ONT reads with the Verkko pipeline and used Strand-seq to phase assembly graphs^71^. Lastly, G4 samples were assembled using HiFi reads only with hifiasm (v0.19.5)^9^.

NOTE: Trio-based phasing with Verkko assigns maternal to haplotype 1 and paternal to haplotype2. In contrast, for hifiasm assemblies we report switched haplotype labeling such that haplotype 1 is paternal and haplotype 2 is maternal in order to match HPRC standard for hifiasm assemblies.

### Evaluation of phased genome assemblies

To evaluate the base pair and structural accuracy of each phased assembly, we employed a multitude of assembly evaluation tools as well as orthogonal datasets such as PacBio HiFi, ONT, Strand-seq, Illumina, and Element data. Known assembly issues are listed in **Supplementary Table 4**.

### Strand-seq validation

We used Strand-seq data to evaluate directional and structural accuracy of each phased assembly. First, we aligned selected Strand-seq libraries for each sample to the phased *de novo* assembly using BWA^62^ (v0.7.17-r1188). Then we ran breakpointR^69^ (v1.15.1) using aligned BAM files as input. Next, we created directional composite files using breakpointR function ‘createCompositeFiles’ followed by running breakpointR on such composite files using ‘runBreakpointR’ function. This provided us, for any given sample, with regions where strand-state changes across all single-cell Strand-seq libraries. Many such regions point to real heterozygous inversions. However, regions where Strand-seq reads mapped in opposite orientation with respect to surrounding regions are likely caused by misorientation. Also positions where the strand state of Strand-seq reads changes repeatedly in multiple libraries might be a sign of an assembly misjoin and such regions were investigated more closely to rule out any such large structural assembly inconsistencies.

### Read to assembly alignment

To evaluate *de novo* assembly accuracy, we aligned sample-specific PacBio HiFi reads to their corresponding phased genome assemblies using Winnowmap (v2.03) with the following parameters:

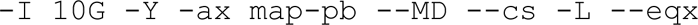

### Flagger validation

Flagger^9^ was used to detect misassemblies using HiFi read alignments to the assemblies and the assemblies aligned to the reference genome (github.com/mrvollger/asm-to-reference-alignment.git). Regions were flagged based on read alignment divergence and specific reference-biased regions. A reference-specific BED file (chm13v2.0.sd.bed) was used setting a maximum read divergence of 2% and specifying reference-biased blocks. These flagged regions were analyzed to identify collapses, false duplications, erroneous regions, and correctly assembled haploid blocks with the expected read coverage.

### NucFreq validation

NucFreq^18^ was used to calculate nucleotide frequencies for HiFi reads aligned using Winnowmap^72^. This was used to identify regions of collapses: where the second-highest nucleotide count exceeded 5, and misassembly: where all nucleotide counts were zero.

### Misjoin evaluation

We used paftools.js script, which is part of minimap2^73^, to detect assembly gaps, inversions, and interchromosomal misjoins within an alignment of each *de novo* assembly to the reference genome. This was done by calling the paftools.js misjoin function.

### Assembly base-pair quality

To evaluate the accuracy of the genome assembly, we employed a pipeline that uses Meryl (v1.0) to count the *k*-mers of length 21 from Illumina reads using the following command:

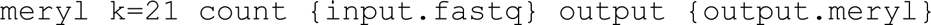

We then used Merqury (v1.1)^74^, which compares the *k*-mers from the sequencing reads against those in the assembled genome and flags discrepancies where *k*-mers are uniquely found only in the assembly. These unique *k*-mers indicate potential base-pair errors. Merqury then calculates the quality value based on the *k*-mer survival rate, estimated from Meryl’s *k*-mer counts, providing a quantitative measure to assess the completeness and correctness of the genome assembly.

### Gene completeness validation

To evaluate the completeness of single-copy genes in our assemblies, we used compleasm^75^ (v0.2.4). See more details at https://github.com/huangnengCSU/compleasm.

### Assembly to reference alignment

All *de novo* assemblies were aligned to both GRCh38 as well as to the complete version of the human reference genome T2T-CHM13 (v2) using minimap2^73^ (v2.24) with the following command:

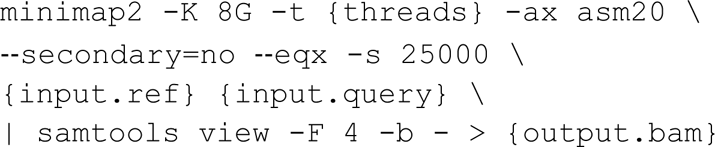

A complete pipeline for this reference alignment is available at GitHub (https://github.com/mrvollger/asm-to-reference-alignment).

We also generated a trimmed version of these alignments using rustybam (v0.1.33) function ‘trim-paf’ to trim redundant alignments that mostly appear at highly identical SDs. With this, we aim to reduce the effect of multiple alignments of a single contig over these duplicated regions.

### Definition of stable diploid regions

For this analysis we use assembly to reference alignments (see ‘Assembly to reference alignment’ section) reported as PAF files. We used trimmed PAF files reported by the rustybam trim-paf function. Stable diploid regions were defined as regions where phased genome assemblies report exactly one contig alignment for haplotype 1 as well as haplotype 2 and are assigned as ‘2n’ regions. Any region with two or more alignments per haplotype is assigned as ‘multi’ alignment. Lastly, regions with only single contig alignment in a single haplotype are assigned as ‘1n’ regions. These reports were generated using the ‘getPloidy’ R function (**Code Availability**).

### Detection and analysis of meiotic recombination breakpoints

We constructed a high-resolution recombination map of this family using three orthogonal approaches that differ either based on underlying sequencing technology or detection algorithm applied to the data. The first approach is based on chromosome-length haplotypes extracted from Strand-seq data using R package StrandPhaseR^22,76^ (https://github.com/daewoooo/StrandPhaseR). The second approach uses inheritance vectors derived from Mendelian consistency of small variants across the family pedigree^13^. Our final approach utilizes trio-based phased genome assemblies followed by small variant calling using PAV and Dipcall to more precisely define the meiotic breakpoints.

### Detection of recombination breakpoints using circular binary segmentation

To map meiotic recombination breakpoints using circular binary segmentation, we used two different datasets. The first dataset represents phased small variants (SNVs and indels) as reported by Strand-seq-based (SSQ) phasing^22,76^. The other is based on small variants reported in trio-based phased assemblies either by PAV^8^ (v2.3.4) or Dipcall^77^ (v0.3). With this approach we set to detect recombination breakpoints as positions where a child’s haplotype switches from matching H1 to H2 of a given parent or vice versa. To detect these positions, we first established what homolog in a child was inherited from either parent by calculating the level of agreement between child’s alleles and homozygous variants in each parent. Next, we compared each child’s homolog to both homologs of the corresponding parent and encoded them as 0 or 1 if they match H1 or H2, respectively. We applied a circular binary segmentation algorithm on such binary vectors by using the R function “fastseg” implemented in R package fastseg^78^ (v1.46.0) with the following parameters: `fastseg(binary.vector, minSeg={}, segMedianT = c(0.8, 0.2))`. In case of sparse Strand-seq haplotypes we set the fastseg parameter `“minSeg”` set to 20 and in case of dense assembly-based haplotypes we used a larger window of 400 and 500 for Dipcall- and PAV-based variant calls to achieve comparable sensitivity in detecting recombination breakpoints. Then the regions with segmentation mean ≤0.25 are marked as H1 while regions with segmentation mean ≥0.75 are assigned as H2. Regions with segmentation mean in between these values were deemed ambiguous and were excluded. In addition, we filtered out regions shorter than 500 kbp and merged consecutive regions assigned the same haplotype (**Code Availability**).

### Detection of meiotic recombination breakpoints using inheritance vectors

DeepVariant calls from HiFi sequencing data from G1, G2, and G3 pedigree members allow us to identify the haplotype of origin for heterozygous loci in G3 and infer the occurrence of a recombination along the chromosome when the haplotype of origin changes between loci. An initial outline of the inheritance vectors was identified by first applying a depth filter to remove variants outside the expected coverage distribution per sample, inheritance was then sketched out via a custom script, requiring a minimum of 10 single-nucleotide polymorphisms (SNPs) supporting a particular haplotype, and manually refined to remove biologically unlikely haplotype blocks, or add additional haplotype blocks, where support existed, and refine haplotype coordinates. Missing recombinations were identified from the occurrence of blocks of pedigree violating variants, matching the location of assembly-based recombination calls. We developed a hidden Markov model framework to identify the most probable sequence of inheritance vectors from SNP sites using the Viterbi algorithm. The transition matrix defines the probability of a given inheritance state transition (recombination). While the emission matrix defines the probability that variant calls at a particular locus accurately describe the inheritance state. The values contained within transition and emission matrices were refined to recapitulate the previously identified inheritance vectors, while correctly identifying missing vectors. The Viterbi algorithm identified 539 recombinations, a maternal recombination rate of 1.29 cM/Mbp and a paternal recombination rate of 0.99 cM/Mbp. Maternal bias was observed in the pedigree, with 57% of recombinations identified in G3 of maternal origin.

### Merging of meiotic recombination maps

Meiotic recombination breakpoints reported by different orthogonal technologies and algorithms (see ‘Detection of meiotic recombination breakpoints’ section) were merged separately for G2 and G3 samples. We started with the G3 recombination map where we used an inheritance-based map as a reference and then looked for support of each reference breakpoint in recombination maps reported based on PAV, Dipcall, and Strand-seq (SSQ) phased variants. A recombination breakpoint was supported if for a given sample and homolog an orthogonal technology reported a breakpoint no further than 1 Mbp from the reference breakpoint. Any recombination breakpoint that is further apart is reported as unique. We repeated this for the G2 recombination map as well. However, in the case of the G2 recombination map we used a PAV-based map as a reference. This is because inheritance-based approaches need three generations in order to map recombination breakpoints in G3. We also report a column called ‘best.range’ that is the narrowest breakpoint across all orthogonal recombination maps that directly overlaps with a given reference breakpoint. Lastly, we report a ‘min.range’ column that represents for any given breakpoint a range with the highest coverage across all orthogonal datasets. Merged recombination breakpoints are reported in **Supplementary Table 8**.

### Meiotic recombination breakpoint enrichment

We tested enrichment of all (n=678) recombination breakpoints detected in G2-G3 with respect to T2T-CHM13 if they cluster towards the ends of the chromosomes depending on parental homolog origin. For this we counted the number of recombination breakpoints in the last 5% of each chromosome end specifically for maternal and paternal breakpoints. Then we shuffled detected recombination breakpoints along each chromosome for 1000 times and redo the counts. For the permutation analysis we used R package regioneR^79^ (v1.32.0) and its function ‘permTest’ with the following parameters:

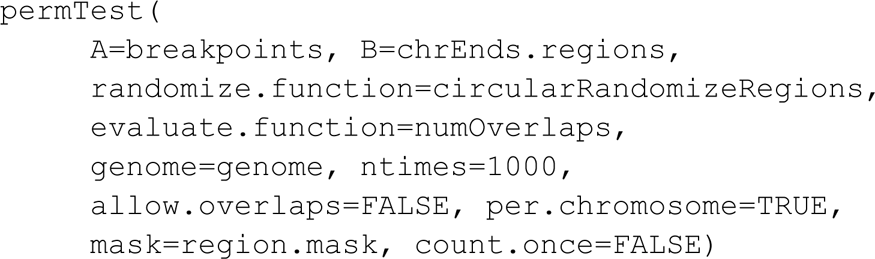

### Refinement of meiotic recombination breakpoints using multiple sequence alignment (MSA)

Up to this point all meiotic recombination breakpoints were called using variation detected with respect to a single linear reference (GRCh38 or T2T-CHM13). To alleviate any possible biases introduced by comparison to a single reference genome, we set out to refine detected recombination breakpoints for each inherited homolog (in child) directly in comparison to parental haplotypes from whom the homolog was inherited from. We start with a set of merged T2T-CHM13 reference breakpoints for G3 only by selecting the ‘best.range’ column (**Supplementary Table 8**). Then for each breakpoint we set a ‘lookup’ region to 750 kbp on each side from the breakpoint boundaries and used SVbyEye (**Code Availability**) function ‘subsetPafAlignments’ to subset PAF alignments of a phased assembly to the reference (T2T-CHM13) to a given region. We follow by extracting the FASTA sequence for a given region from the phased assembly. We did this separately for inherited child homologs (recombined) and corresponding parental haplotypes that belong to a parent from whom the child homolog was inherited from.

Next, we created an MSA for three sequences (child-inherited homolog, parental homolog 1, and parental homolog 2) using the R package DECIPHER^80^ (v2.28.0; function ‘AlignSeqs’). Fasta sequences whose size differ by more than 100 kbp or their nucleotide frequencies differ by more than 10,000 bases are skipped due to increased computational time needed to align such different sequences optimally using DECIPHER. After MSA construction, we selected positions with at least one mismatch and also removed sites where both parental haplotypes carry the same allele. A recombination breakpoint is a region where the inherited child homolog is partly matching alleles coming from parental homologs 1 and 2. We, therefore, skipped analysis of MSAs in which a child’s alleles are more than 99% identical to a single parental homolog. If this filter is passed, we use custom R function ‘getAlleleChangepoints’ (**Code Availability**) to detect changepoints where the child’s inherited haplotype switches from matching alleles coming from parental haplotype 1 to alleles coming from parental haplotype 2. Such MSA-specific changepoints are then reported as a new range where a recombination breakpoint likely occurred. Lastly, we attempt to report reference coordinates of such MSA-specific breakpoints by extracting 1 kbp long *k*-mers from the breakpoint boundaries and matching such *k*-mers against reference sequence (per chromosome) using R package Biostrings (v2.70.2) with its function ‘matchPattern’ and allowing for up to 10 mismatches. A list of refined recombination breakpoints is reported in **Supplementary Table 8**.

### Detection of allelic gene conversion using phased genome assemblies

We set out to detect smaller localized changes in parental allele inheritance using a previously defined recombination map of this family. We did this analysis for all G3 samples in comparison to G2 parents. For this we iterated over each child’s homolog (in each sample) and compared it to both parental homologs from which the child’s homolog was inherited from. We did this by comparing SNV and indel calls obtained from phased genome assemblies between the child and corresponding parent. To consider only reliable variants we kept only those supported by at least two read-based callers (either DeepVariant-HiFi, Clair3-ONT or dragen-Illumina callset). We further kept only variable sites that are heterozygous in the parent and were also called in the child. After such strict variant filtering, we slide by two consecutive child’s variants at a time and compare them to both haplotype 1 and haplotype 2 of the respective parent-of-origin. For this similarity calculation we use the custom R function ‘getHaplotypeSimilarity’ (**Code Availability**). Then for each haplotype segment, defined by recombination breakpoints, we report regions where at least two consecutive variants match the opposing parental haplotype in contrast to the expected parental homolog defined by recombination map. We further merge consecutive regions that are ≤5 kbp apart. For the list of putative gene conversion events, we kept only regions that have not been reported as problematic by Flagger. We also removed regions that are ≤100 kbp from previously defined recombination events and events that overlap centromeric satellite regions and highly identical SDs (≥99% identical). Lastly, we evaluated the list of putative AGC events by visual inspection of phased HiFi reads.

### STR/VNTR analysis

#### Defining the TR catalogs

The command `trf-mod -s 20 −l 160 {reference.fasta}` was used, resulting in a minimum reference locus size of 10 bp and motif sizes of 1 to 2000 bp (https://github.com/lh3/TRF-mod)^81^. Loci within 50 bp were merged, and then any loci >10,000 bp were discarded. The remaining loci were annotated with tr-solve (https://github.com/trgt-paper/tr-solve) to resolve locus structure in compound loci. Only TRs annotated on chromosomes 1-22, X, and Y were considered. The TR catalogs are available on Zenodo DOI: 10.5281/zenodo.13178746.

#### TR genotyping with TRGT

TRGT is a software tool for genotyping TR alleles using PacBio HiFi sequencing reads^28^. Provided with aligned HiFi sequencing reads (in BAM format) and a file that enumerates the genomic locations and motif structures of a collection of TR loci, TRGT will return a VCF file with inferred genotypes at each TR locus. In this analysis, we ran TRGT (v0.7.0-493ef25) on each member of the 1463 pedigree using the TR catalog defined above. TRGT was run using default parameters:

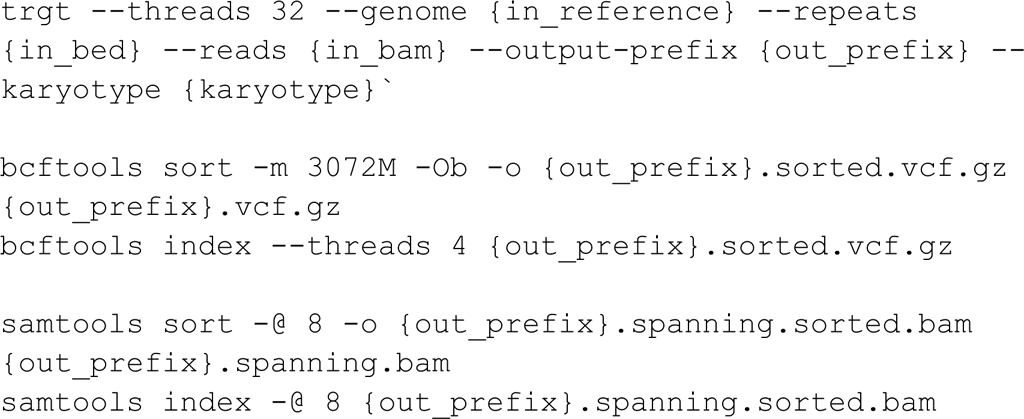

#### Measuring concordant inheritance of TRs

To determine the concordant inheritance of TRs, we calculated the possible Manhattan distances derived from all possible combinations of a proband’s allele length (AL) from TRGT with both the maternal and paternal AL values. We considered a locus concordant if the minimum Manhattan distance from all computed distances was found to be 0, suggesting that a combination of the proband’s AL values matched the parental AL values perfectly. In contrast, if the minimum Manhattan distance was greater than 0, suggesting that all combinations of the proband’s AL values exhibited some deviation from the parental AL values, we regarded the locus as discordant and recorded it as a potential Mendelian inheritance error. For each TR locus, we calculated the number of concordant trios, the number of MIE trios, and the number of trios that had missing values and could not be fully genotyped. Loci with any missing genotypes were excluded when calculating the percent concordance; however, individual complete trios were considered for *de novo* variant calling below.

#### Calling de novo TRs

We focused *de novo* TR calling on G3 for several reasons. First, their G2 parents were sequenced to 99 and 109 HiFi sequencing depths, resulting in a far lower chance of parental allelic dropout than samples with more modest sequencing depths. Second, G1 DNA was derived from cell lines, increasing the risk of artifacts when calling DNMs in G2. And finally, DNMs in the two individuals in G3 with sequenced children in our study can be further assessed by transmission.

We used TRGT-denovo^29^ (v0.1.3), a companion tool to TRGT, to enable in-depth analysis of TR DNMs in family trios using HiFi sequencing data. TRGT-denovo uses consensus allele sequences and genotyping data generated by TRGT, and also incorporates additional evidence from spanning HiFi reads used to predict these allele sequences. Briefly, TRGT-denovo extracts and partitions spanning reads from each family member (mother, father, and child) to their most likely alleles. Parental spanning reads are realigned to each of the two consensus allele sequences in the child, and alignment scores (which summarize the difference between a parental read and a consensus allele sequence) are computed for each read. At every TR locus, each of the two child alleles is independently considered as a putative *de novo* candidate. For each child allele, TRGT-denovo reports the presence or absence of evidence for a *de novo* event, which includes the following: `denovo_coverage` (the number of reads supporting a unique AL in the child that is absent from the parent’s reads); `overlap_coverage` (the number of reads in the parents supporting an AL that is highly similar to the putative *de novo* allele); and magnitude of the putative *de novo* event (expressed as the absolute mean difference of the read alignment scores with *de novo* coverage relative to the closest parental allele).

#### Calculating the size of a de novo TR expansion or contraction

We measured the sizes of *de novo* TR alleles with respect to the parental TR allele that most likely experienced a contraction or expansion event. If TRGT-denovo reported a *de novo* expansion or contraction at a particular locus, we did the following to calculate the size of the event.

Given the ALs reported by TRGT for each member of the trio, we computed the difference in size (which we call a “diff”) between the *de novo* TR allele in the child and all four TR alleles in the child’s parents. For example, if TRGT reported ALs of `100,100` in the father, `50,150` in the mother, and `200,100` in the child, and the allele of length `200` was reported to be *de novo* in the child, the “diffs” would be `100,100` in the father and `150,50` in the mother. If we were able to phase the *de novo* TR allele to a parent-of-origin, we simply identify the minimum “diff” among that parent’s ALs and treat it as the likely expansion/contraction size. Otherwise, we assume that the smallest “diff” across all parental ALs represents the likely *de novo* size.

#### De novo filtering

We applied a series of filters to the candidate TR DNMs (identified by TRGT-denovo) to remove likely false positives. For each *de novo* allele observed in a child, we required the following:

- HiFi sequencing depth in the child, mother, and father ≥10 reads
- the candidate *de novo* AL in the child must be unique

- as in ^33^, we removed candidate *de novo* TR alleles if a) the child’s *de novo* AL matched one of the father’s ALs and the child’s non-*de novo* AL matched one of the mother’s ALs or b) the child’s *de novo* AL matched one of the mother’s ALs and the child’s non-*de novo* AL matched one of the father’s ALs
- the candidate *de novo* allele must represent an expansion or contraction with respect to the parental allele
- at least two HiFi reads supporting the candidate *de novo* allele `(denovo_coverage >= 2)` in the child, and at least 20% of total reads supporting the candidate *de novo* allele `(child_ratio >= 0.2)`
- fewer than 5% of parental reads likely supporting the candidate *de novo* AL in the child

To calculate TR DNM rates in a given individual, we first calculated the total number of TR loci (among the ∼7.8 million loci genotyped using TRGT) that were covered by at least 10 HiFi sequencing reads in each member of the focal individual’s trio (i.e., the focal individual and both of their parents). We then divided the total count of *de novo* TR alleles by the total number of “callable” loci to obtain an overall DNM rate, expressed per locus per generation. Finally, we divided that rate by 2 to produce a mutation rate expressed per locus, per haplotype, per generation. We also estimated DNM rates for each motif size (e.g., a motif size of 1 corresponds to homopolymers, a motif size of 2 to dinucleotides, etc.) using a similar approach; for a given motif size, we counted the number of TR DNMs that occurred at motifs of that size and divided the count by the total number of TR loci of the specified motif size that passed filtering thresholds. We then divided that rate by 2 to produce a mutation rate per locus, per haplotype, per generation.

Prior studies usually measured STR mutation rates at loci that are polymorphic within the cohort of interest. To generate mutation rate estimates that are more consistent with these prior studies, we also calculated the number of STR loci that were polymorphic within the CEPH 1463 cohort. Loci were defined as polymorphic if at least two unique ALs were observed among the CEPH 1463 individuals at a given TR locus. We note that this definition of “polymorphic” STRs is sensitive to both the size of the cohort and the sequencing technology used to genotype STRs. As discussed in prior studies^33^, the number of polymorphic loci is proportional to the size of the cohort. Moreover, by defining loci as polymorphic if we observed more than one unique AL across the cohort, we may erroneously classify loci as polymorphic if HiFi sequencing reads exhibited a substantial amount of “stutter” at those loci, producing variable estimates of STR ALs across individuals. A total of 1,096,430 STRs were polymorphic within the cohort. To calculate mutation rates in each G3 individual, we applied the same coverage quality thresholds as described above.

#### Phasing TRs

The STRs genotyped by TRGT were phased using HiPhase^82^ (v1.0.0-f1bc7a8). We followed HiPhase’s guidelines for jointly phasing small variants, SVs, and TRs by inputting the relevant VCF files from DeepVariant, PBSV, and TRGT into HiPhase, resulting in three phased VCF files for each analyzed sample. We also activated global realignment through the `--global-realignment-cputime` parameter to improve allele assignment accuracy. Note that HiPhase specifically excludes variants that fall entirely within genotyped STRs from the phasing process. This is motivated because STRs often encompass numerous smaller variants.

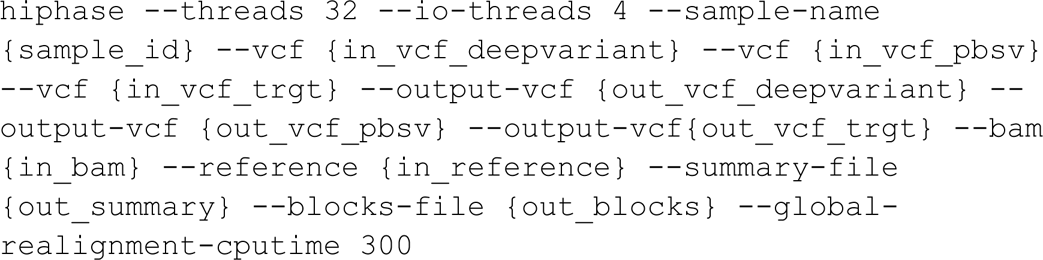

#### Parent-of-origin determination

We used the phased genotypes inferred by HiPhase to determine the likely parent-of-origin for *de novo* TR expansions and contractions. For each phased *de novo* allele that we observed in a child, we examined all informative SNVs in that child’s parents ±500 kbp from the *de novo* allele. We defined informative sites using the following criteria: sites must be biallelic SNVs; total read depth in the mother, father, and child must be at least 10 reads; Phred-scaled genotype quality in the mother, father, and child must be at least 20; the child’s genotype must be heterozygous; and the parents’ genotypes must not be identical-by-state. Using the child’s phased SNV VCF, we then determine whether the child’s REF or ALT allele at the informative site was inherited from either the mother or father. For example, if the mother’s genotype is `0/0`, the father’s genotype is `0/1` (note that the parental genotypes need not be phased), and the child’s genotype is `1|0`, we know that the child’s “first” haplotype was inherited from the father and the “second” haplotype was inherited from the mother. We repeat this process for all informative sites within the ±250 kbp interval. We then find the *N* informative sites that are a) closest to the *de novo* TR allele (either upstream or downstream) while b) supporting a consistent inheritance pattern in the child (i.e., all support the same parent-of-origin for the child’s two haplotypes and c) all reside within the same HiPhase phase block (defined using the `PS` tag in the HiPhase output VCF). Finally, we use the phased TR VCF produced by HiPhase to check whether the *de novo* allele was phased to either the first or second haplotype in the child. We then confirm that the *de novo* allele shares the same `PS` tag as the informative sites identified above and use the *N* informative sites to determine whether the haplotype to which the *de novo* allele was phased was likely inherited from either the mother or the father.

#### Measuring concordance with orthogonal sequencing technology

At each candidate *de novo* TR allele, we calculated concordance between the *de novo* ALs estimated by TRGT and the ALs supported by Element, ONT, or HiFi reads. We restricted our concordance analyses to autosomal TR loci with a single expansion or contraction (i.e., we did not analyze “complex” TR loci harboring multiple unique expansions and/or contractions).

TRGT reports two AL estimates for every member of a trio at an autosomal TR locus, and TRGT-denovo assigns one of these two ALs to be the *de novo* AL in the child. At each TR locus, we calculated the difference between the length of the locus in the reference genome (in base pairs) and each of the two ALs in a given individual. We refer to the difference between the TRGT AL and the reference locus size as the “relative AL.” We then queried BAM files containing Element, Illumina, ONT, or PacBio HiFi reads at each TR locus. Using the `pysam` library (https://github.com/pysam-developers/pysam), we iterated over all reads that completely spanned the TR locus and had a mapping quality of 60. To estimate the AL of a TR expansion/contraction in a read with respect to the reference genome, we counted the number of nucleotides associated with every CIGAR operation that overlapped the TR locus. For example, an Element read might have the following CIGAR string: `100M2D10M6I32M`. For each of the CIGAR operations that overlap the TR locus, we increment a counter by `OP * BP`, where `OP` equals `0` for “match” CIGAR operations, `1` for “insertion” operations, and `-1` for “deletion” operations, and `BP` equals the number of base pairs associated with the given CIGAR operation. Thus, at each TR locus, we generated a distribution of “net CIGAR operations” in each member of the trio.

We used these “net CIGAR operations’’ to validate candidate *de novo* TR alleles in each child. For each *de novo* TR allele, we calculated the number of Element reads in the child that supported the *de novo* AL estimated by TRGT (allowing the Element reads to support the *de novo* AL ±1 bp). We then calculated the number of Element reads in that child’s parents supporting the *de novo* AL. If at least one Element read supported the *de novo* TR AL in the child, and zero Element reads supported the *de novo* TR AL in both parents, we considered the *de novo* TR to be validated.

#### Validating recurrent TR DNMs

To assemble a confident list of candidate recurrent *de novo* TR alleles, we first assembled a list of TR loci where two or more 1463 individuals (in either G2, G3, or G4) harbored evidence for a *de novo* TR allele. For each candidate locus, we then required that all members of the CEPH 1463 pedigree were genotyped for a TR allele at the locus and had at least 10 aligned HiFi reads at the locus. These filters produced a list of 49 candidate loci where we observed evidence of either intragenerational or intergenerational recurrence. We visually inspected HiFi read evidence using the Integrated Genomics Viewer (IGV)^83^, as well as bespoke plots of HiFi CIGAR operations, at each locus to determine whether the candidate *de novo* TR alleles appeared plausible.

### Read-based variant calling

PacBio HiFi data were processed with the human-WGS-WDL (https://github.com/PacificBiosciences/HiFi-human-WGS-WDL/releases/tag/v1.0.3). The pipeline aligns, phases, and calls small variants (using DeepVariant) and SVs (using PBSV). We used the aligned haplotype-tagged HiFi BAMs for all downstream PacBio analysis.

### Clair3

Clair3^84^ (v1.0.7) variant calls were made based on the alignments with default models for PacBio HiFi and ONT (ont_guppy5) data, respectively, with phasing and gVCF generation enabled. Variant calling was conducted on each chromosome individually and concatenated into one VCF. gVCFs were then fed into GLNexus^85^ with a custom configuration file.

PacBio HiFi

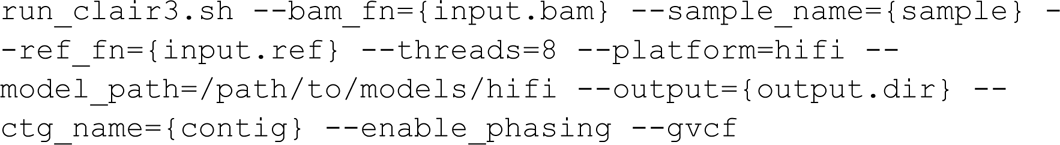

ONT

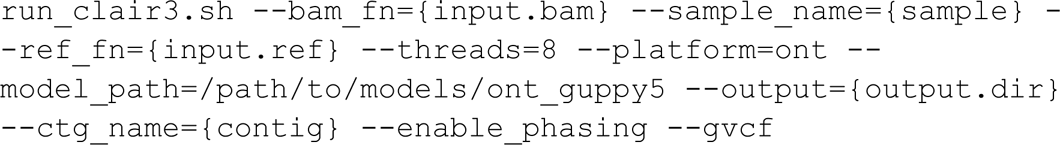

### Generation of truth set of genetic variation using inheritance vectors

We used a previously established framework to define ground truth genetic variation^13^. Our analysis, unlike trio-based filtering, uses all four alleles to detect genotyping errors, whereas in a trio only two alleles are transmitted and observed. By testing the genotype patterns in the third generation against the phased haplotypes of the first generation (A,B,C,D), we can test for the correct transmission of alleles from the second to third generations. We establish a map of the haplotypes across the third generation (inheritance vector) from which we can adjudicate variant calls against. To test for pedigree consistency, we implemented code that uses the inheritance vector as the expected haplotypes and test the possible genotype configurations within the query VCF file. Using the haplotype structure we phase the pedigree consistent variants. These functions are implemented as a single binary tool that requires the inheritance vectors and a standard formatted VCF file (e.g.,):

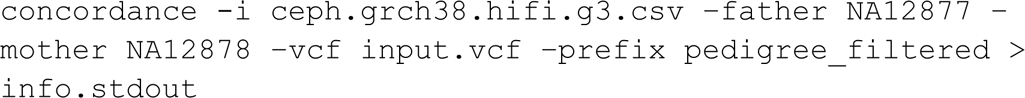

The pedigree filtering and additional steps to build a small variant truth set can be found in the following GitHub repository: https://github.com/Platinum-Pedigree-Consortium/Platinum-Pedigree-Inheritance/tree/main.

### Detection of small *de novo* variants

Following the parameters outlined in Noyes et al.^10^, we called variants in HiFi data aligned to T2T-CHM13 using GATK HaplotypeCaller (v4.3.0.0) and DeepVariant^86^ (v1.4.0) and naively identified variants unique to each G2 and G3 sample^86^. We separated out SNV and indel calls and applied basic quality filters, such as removing clusters of three or more SNVs in a 1 kbp window. We combined this set of variant calls generated by a secondary calling method, (https://github.com/Platinum-Pedigree-Consortium/Platinum-Pedigree-Inheritance/blob/main/analyses/Denovo.md) and subjected all calls to the following validation process.

We validated both SNVs and indels by examining them in HiFi, ONT, and Illumina read data, excluding reads that failed to reach mapping quality (59 for long reads, 0 for short reads) thresholds. Reads with high base quality (>20) and low base quality (<20) at the variant site were counted separately. We retained variants that were present in at least two types of sequencing data for the child, and absent from high base quality parental reads. For SNV calls, we next examined HiFi data for every sample in the pedigree. We determined an SNV was truly *de novo* if it was absent from every family member that was not a direct descendant of the *de novo* sample. Finally, we examined the allele balance of every variant, determined which variants were in TRs, and reevaluated parental read data across all sequencing platforms, removing variants with noisy sequencing data or more than two low-quality parental reads supporting the alternate allele.

### DNM phasing and postzygotic assignment

To determine the parent-of-origin for the *de novo* SNVs, we reexamined the long reads containing the *de novo* allele. First, we used our initial GATK variant calls to identify informative sites in an 80 kbp window around the DNM, selecting any SNPs where one allele could be uniquely assigned to one parent (for example, a site that is homozygous reference in a father and heterozygous in a mother). For every DNM, we evaluated every ONT and HiFi read that aligned to the site of the *de novo* allele and assigned it to either a paternal or maternal haplotype (if informative SNPs were available) by calculating an inheritance score as outlined in Noyes et al.^10^ DNMs that were exclusively assigned to maternal or paternal haplotypes were successfully phased, whereas DNMs on conflicting haplotypes were excluded from our final callset. Unphased variants were determined to be postzygotic in origin (n=7) if their allele balance was not significantly different across platforms (by a chi-squared test) and if their combined allele balance was significantly different from 0.5.

Once we assigned every read to a parental haplotype, we counted the number of maternal and paternal reads that had either the reference or alternate allele. We determined that a DNM was germline in origin if it was present on every read from a given parent’s haplotype. Conversely, if a DNM was present on only a fraction of reads from a parental haplotype, we determined that it was postzygotic in origin.

### Sex chromosome DNM calling and validation

To identify DNMs on the X chromosome, we applied the same strategy as autosomal variants, with one exception: we only used variant calls generated by GATK. For males, we reran GATK in haploid mode, such that it would only identify one genotype on the X chromosome.

To identify DNMs on the Y chromosome, we aligned male HiFi, ONT, and Illumina data to the G1-NA12889 chrY assembly and then called variants using GATK in haploid mode on the aligned HiFi data. We directly compared each male to his father, selecting variants unique to the son. We validated SNVs and indels by examining the father’s HiFi, ONT, and Illumina data and excluded any variants that were present in the parental reads, applying the same logic that we used for autosomal variants.

### Callable genome and mutation rate calculations

We determined the size of the callable genome for each individual based on their HiFi data, using two criteria. First, we reran GATK HaplotypeCaller with the option `“ERC BP_RESOLUTION”` for every *de novo* sample and their parents to generate a genotype at every site in the genome. We excluded any site where both parents were not homozygous for the reference allele. For male sex chromosomes, we only considered the mother’s genotype in the case of the X, and the father’s genotype in the case of the Y. Second, we examined the HiFi data for each sample and their parents and excluded any site where all three members of the trio did not have at least one HiFi read that passed our mapping and base quality thresholds. Any sites that were not excluded were considered to be “callable” with our DNM pipeline. We intersected these sites with annotations to calculate the amount of callable space in a region such as SDs. To calculate the mutation rate on the autosomes in each sample, we divided the number of DNMs in a given region by twice the number of bases deemed to be callable.

### Detection and filtering of *de novo* SVs

We attempted to obtain putative *de novo* SVs from three different sources. The first one is based on reporting *de novo* SVs from read-based callsets (PBSV, Sniffles, Sawfish). The second reports putative *de novo* SVs from variants called in phased genome assemblies. The last utilized pangenome graphs constructed from phased genome assemblies to report *de novo* SVs.

### Assembly-based detection of *de novo* SVs

1. SVPOP (v3.4.0) (https://github.com/EichlerLab/svpop) was used to produce a merged PAV callset across all samples. It merges a single source (single SV caller) across multiple samples. The merge definition used was: “nr::ro:szro:exact:match” The samples were provided in this order (G1-G2-G3): “NA12889”, “NA12890”, “NA12891”, “NA12892”, “NA12877”, “NA12878”, “NA12879”, “NA12881”, “NA12882”, “NA12883”, “NA12884”, “NA12885”, “NA12886”, “NA12887”
2. For each sample in G3, we selected variants unique to that sample alone.
3. To compare variant calls against the previous generation, SVPOP was used again to do a PBSV/PAV intersection. This involved intersecting the PAV calls for G3 with the PBSV calls for G2, comparing each sample in G3 against each sample in G2.
4. The callable BED files from PAV, intersections with G2’s PBSV calls, and the list of putative *de novo* calls went into our validation pipeline.
5. The pipeline:

a. Checks if the putative *de novo* variant was called by PBSV in either parent.
b. Checks if the putative *de novo* variant is seen in HiFi reads in either parent by running subseq (https://github.com/EichlerLab/subseq).
c. Checks if the variant was in a callable region in either parent.
d. Performs an MSA using DECIPHER of the two haplotypes of the sample, and both parents, in the location of the SV with 1000 bp flank on either side.

### Pangenome graph detection on *de novo* SVs

Verkko assemblies were partitioned by chromosome by mapping them against the GRCh38, T2T-CHM13, and HG002 (v1.0.1) human reference genomes using WFMASH (v0.13.1, commit 251f4e1) pangenome aligner. On each set of contigs, we applied PGGB (v0.6.0, commit 87510bc) to build chromosome-level unbiased pangenome variation graphs^4^ with the following parameters: `-s 20k -p 95 -k 47 -V` chm13:100000, grch38:100000. We used Variation graph toolkit^87^ (v1.40.0) to call variants from the graphs with respect to both T2T-CHM13 and GRCh38 reference genomes. Variants were then decomposed by applying VCFBUB (v0.1.0, commit 26a1f0c) to retain those found in top-level bubbles that are anchored on the genome used as reference, and VCFWAVE (v1.0.3) to homogenize SV representation across samples. Subsequently, raw VCF files were used as an input for pedigree-based filtering of putative *de novo* SVs.

### *de novo* SV filtering in SV callsets (PGGB, PAV, PBSV, Sniffles, Sawfish)

*de novo* filtering was done using BCFtools +fill-tags followed by filtering the joint-called VCF for singleton-derived alleles at sites where all samples had a genotype call. By considering all G2/G3 family members (not just trios), we increased *de novo* SV specificity. We used the command line:

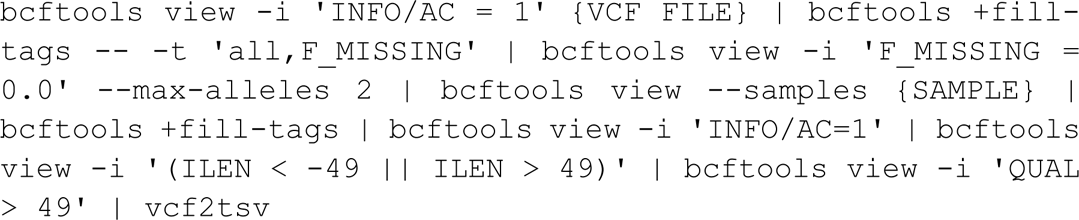

### Evaluation of putative *de novo* SVs

All predicted *de novo* SVs were evaluated by Verkko as well as hifiasm (UL) assemblies. We did this by extracting a sequence around the SV by adding two times the size of the SV on each side. We extracted the sequence from a G3 individual and corresponding G2 parents. Next, we constructed the MSA and visually check if the predicted SV is visible in both Verkko and hifiasm (UL) assemblies.

All predicted *de novo* SVs were subsequently merged into a nonredundant callset that have been further manually validated using manual inspection in the IGV browser^83^. All passed variants were then evaluated in a sense of possible mechanism that could explain each putative *de novo* variant.

### Extracting donor site of *de novo* SVA insertion

We first extracted an inserted SVA element in the *de novo* Verkko assembly of NA12887 (maternal haplotype, haplotype1). Next, we used minimap2^73^ (v2.24) to align this ∼3.4 kbp long piece of DNA to both maternal and paternal Verkko assemblies using the parameters reported below:

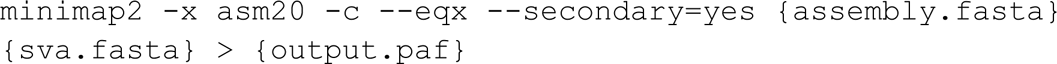

With these parameters we reported all locations of this DNA segment. We defined a putative donor site as an alignment position in maternal haplotype that has nearly perfect match with SVA *de novo* insertion.

### Analysis of centromeric regions

To identify completely and accurately assembled centromeres from each genome assembly, we first aligned the genome assemblies generated via Verkko^16^ or hifiasm (UL)^17^ to the T2T-CHM13 reference genome^1^ using minimap2^73^ and the following parameters: `-a --eqx -x asm20 -s 5000 -I 10G -t {threads}`. Then, we filtered the whole-genome alignments to only those contigs that aligned to the centromeres in the T2T-CHM13 reference genome. We checked if these centromeric contigs spanned the centromeres by checking to see if they contained sequence from the p- and the q-arms in the regions directly adjacent to the centromere. Then, we validated the assembly of the centromeric regions by aligning native PacBio HiFi data from the same source genome to each whole-genome assembly using pbmm2 (v1.1.0; https://github.com/PacificBiosciences/pbmm2) and the following command: align --log-level DEBUG --preset SUBREAD --min-length 5000 -j {threads}, and next assessed the assemblies for uniform read depth across the centromeric regions via NucFreq^18^. We also aligned native ONT data >30 kbp in length from the same source genome to each whole-genome assembly using minimap2^73^ (v2.28) and assessed the assemblies for uniform read depth across the centromeric regions via IGV browser^83^.

To identify *de novo* SVs and SNVs within each centromeric region, we first aligned each child’s genome assembly to the relevant parent’s genome assembly using minimap2^73^ and the following parameters: `-a --eqx -x asm20 -s 5000 -I 10G -t {threads}`. Then, we used the resulting PAF file to identify *de novo* SVs and SNVs using SVbyEye (**Code Availability**), filtering our results to only those centromeres that were completely and accurately assembled. We checked each SV and SNV call with NucFreq^18^, Flagger^9^, and native ONT data to ensure that the underlying data supported each call.

### Analysis of telomeric regions

We processed all G1, G2, and G3 assemblies with Tandem Repeats Finder (TRF)^81^ to determine the existence of the canonical telomeric repeat (p-arm: CCCTAA, q-arm: TTAGGG) within the distal regions of each assembled contig; TRF (v4.09.1) was run with parameters: `‘2 7 7 80 10 50 10 -d -h-ngs’`, recommended for young (in this context, non-deteriorated) repeats as implemented in RepeatMasker (v4.1.6). The assembled contigs, in turn, were aligned to the T2T-CHM13 reference with minimap2^73^ (v2.24) using the *asm20* preset to establish the identities of each sequence (i.e., whether a given contig represented the whole reference chromosome or a part of it, and whether it should be reverse-complemented to represent it canonically). With identities established, TRF annotations were crawled from the outside in (from the 5’ end on p-arms and from the 3’ end on q-arms, with respect to reverse complementarity as reported by minimap2) until the canonical repeat was encountered; incidences of non-canonical interspersed repeats were also retained.

Additionally, PacBio HiFi reads were mapped to the contigs to assess by how many HiFi reads each region of each assembly was supported (coverage depth); distal regions supported by fewer than five HiFi reads were masked. Of the non-acrocentric chromosome ends across all G1, G2, and G3 samples, 74.2% of the Verkko assemblies (893 out of the possible 1,204 across all subjects and haplotypes) were found to terminate in a canonical telomeric repeat (either spanning from the very start or end of the contig, or immediately adjacent to the region masked due to low coverage) with the median length of such repeats being 5,608 bp (**Supplementary Table 3**). Additionally, out of the T2T-CHM13 chromosomes for which both p and q telomeric ends were recovered, 64.6% (221/342) were represented each by a single assembled contig spanning from the p telomere to the q telomere.

The G4 hifiasm assemblies were processed in the same fashion; however, only 56.8% of the telomeric regions (342 out of the possible 602) were recovered (**Supplementary Fig. 3**) with a median length of the canonical repeat being 4,674 bp (**Supplementary Table 3** – same as for G1-G3), and the contiguity was markedly worse: only one chromosome (chr9 in haplotype 1 of subject G4-200101) was verifiably spanned by a single contig (h1tg000017l).

### Y-chromosomal analysis

#### Construction and dating of Y phylogeny

The construction and dating of Y-chromosomal phylogeny for 58 total samples, combining the 14 pedigree males from the current study with 44 individuals, for which long-read-based Y assemblies have previously been published, was done as described previously in detai_l_^48^. In short, all sites were called from the Illumina high-coverage data^14^ of the 14 pedigree males using the approximately 10.4 Mbp of Y-chromosomal sequence previously defined as accessible to short-read sequencing^88^. BCFtools^89,90^ (v1.16) was used with minimum base quality 20, mapping quality 20, and ploidy 1. SNVs within 5 bp of an indel call (SnpGap) and all indels were removed, followed by filtering all calls for a minimum read depth of 3 and a requirement of ≥85% of reads covering the position to support the called genotype. The VCF was merged with a similarly filtered VCF from Hallast et al.^48^ for the 44 individuals using BCFtools, followed by removal of sites with ≥5% of missing calls, that is, missing in more than 3 out of 58 samples, were removed using VCFtools^91^ (v0.1.16). After filtering, a total of 10,404,104 sites remained, including 13,443 variant sites.

The Y haplogroups of each sample were predicted as previously described^92^ and correspond to the International Society of Genetic Genealogy nomenclature (ISOGG, https://isogg.org, v15.73). A coalescence-based method implemented in BEAST^93^ (v1.10.4) was used to estimate the ages of internal nodes. RAxML^94^ (v8.2.10) with the GTRGAMMA substitution model was used to construct a starting maximum-likelihood phylogenetic tree for BEAST. Markov chain Monte Carlo samples were based on 200 million iterations, logging every 1,000 iterations, with the first 10% of iterations discarded as burn-in. A constant-sized coalescent tree prior, the GTR substitution model, accounting for site heterogeneity (gamma), and a strict clock with a substitution rate of 0.76 × 10^−9^ (95% CI = 0.67 × 10^−9^ – 0.86 × 10^−9^) single-nucleotide mutations per bp per year was used^95^. A prior with a normal distribution based on the 95% CI of the substitution rate was applied. A summary tree was produced using Tree-Annotator (v1.10.4) and visualized using the FigTree software (v1.4.4).

#### Identification of sex-chromosome contigs

Detailed analysis of Y-chromosomal DNMs focused on seven males (R1b1a-Z302 Y haplogroup, G1-NA12889, G2-NA12877, G3-NA12882, G3-NA12883, G3-NA12884 and G3-NA12886) for which phased Verkko assemblies were generated. Contigs containing X- and Y-chromosomal sequences were identified and extracted from the whole-genome assemblies as previously described^48^. In addition, the pseudoautosomal regions from the G1 grandmother NA12890 and G2 mother NA12878 genome assemblies were identified by aligning the respective sequences from the T2T-CHM13 reference genome to these assemblies using minimap2^73^ (v2.26).

#### Annotation of Y-chromosomal subregions

The annotation of Y-chromosomal subregions of the Verkko assemblies was performed using both the GRCh38 and T2T-CHM13 Y reference sequences as previously described^48^. The centromeric α-satellite repeats for the purpose of Y subregion annotation were identified using RepeatMasker (v4.1.2-p1) with default parameters. The Yq12 repeat annotations were generated using HMMER^96^ (v3.3.2dev) with published *DYZ1*, *DYZ2*, *DYZ18*, 2k7bp and 3k1bp sequences^48^, followed by manual checking of repeat unit orientation and distance from each other. Dot plots to compare Y-chromosomal sequences were generated using Gepard^97^ (v2.0).

#### Detection and validation of DNMs

Human Y chromosomes vary extensively in the size and composition of repetitive regions^48^, including the T2T-CHM13 Y (haplogroup J1a-L816) and the R1b1a-Z302 haplogroup Y chromosomes carried by the seven pedigree males analyzed in detail here (**Supplementary Figs. 42 and 44**). For this reason, the Y assembly of the G1 grandfather NA12889 was used as a reference for DNM detection (**Supplementary Fig. 45**). The DNMs were called from the Y assemblies of five G2 (NA12877) and G3 (NA12882, NA12883, NA12884, NA12886) males using Dipcall^77^ (v0.3) with the default parameters recommended for male samples. Variants were identified from the male-specific Y regions only, i.e., the pseudoautosomal regions were excluded from this analysis. All identified variants were filtered as follows: any variant calls overlapping with regions flagged by Flagger or NucFreq in either reference or query assembly were filtered out.

For SNVs, the final filtered calls were supported by 100% of HiFi reads (i.e., no reads supported the reference allele in offspring or alternative allele in the father) and ONT reads mapped to both the reference and each individual assembly were checked for support.

For indels (≤50 bp), homopolymer tracts were excluded from the analysis, while the rest of the calls were validated using the read data (HiFi, ONT, Illumina) as follows. Individual reads mapped to the reference (G1 NA12889 Y assembly) and covering the indel call plus 150 bp of flanking sequence were extracted from all samples using subseq (https://github.com/EichlerLab/subseq), followed by alignment using MAFFT (v7.508) with default parameters^98,99^. All alignments were manually checked and any calls where the HiFi data had two or more reads supporting a reference allele and one or more reads supporting an alternate allele were removed. All final SNV and indel calls were additionally supported (if unique mapping to the region was possible) by both Illumina and Element read data mapped to the reference.

For all SV calls, HiFi read depth for reference and alternative alleles were visualized and SVs in regions showing high levels of read depth variation coinciding with clusters of SNVs with >10% of reads supporting an alternative allele removed. HiFi and ONT reads mapped to both the reference and individual assemblies were checked for support.

For all variants, concordance with the expected transmission through generations was confirmed. Additionally, the HiFi data available for three G4 males (200101, 200102 and 200105) were checked for support of the identified variants.

#### Y-chromosomal DNM rate calculation

The assembly-based DNM rates were calculated for each of the five males based on the accessible regions of each individual Y assembly (i.e., any regions flagged by Flagger and/or NucFreq were removed).

### Mobile element analysis

Mobile element analysis was performed on PacBio HiFi reads using xTea^100^ (v0.1.9). Potential non-reference mobile element insertions (MEI) identified with xTea were visualized using IGV to ensure that the insertions were identifiable in the sequencing reads and to determine if any of these events were *de novo.* Using BEDTools^101^, we intersected the non-reference insertions with introns, exons, 5’-UTRs, and 3’-UTRs from T2T-CHM13. To identify potential source elements of the non-reference LINE-1 insertions, we used BLAT^102^ to find the best matching insertion in the T2T-CHM13 reference genome. If there were multiple matches in the reference genome that had the same score, a source element was not called. MEI sequences representing known Alu, L1, and SVA subclasses were obtained from previous work^103^, Dfam^104^, and UCSC Genome Browser^70^. Reference and novel sequences for each MEI class were combined into class-specific files. Sequences were oriented to plus-strand. Highly truncated sequences were removed. MEI sequences were aligned using the MUSCLE^105^ (v3.8.31) aligner. Pairwise distances among MEI sequences were calculated using a Kimera 2-parameter method and then converted to correlations. Principal components (PCs) were obtained by eigenvalue decomposition of the pairwise correlation matrix. The first three PCs were plotted to visualize the relationships among the non-reference MEIs and the known MEI subfamily sequences.

### Ethics declarations

Human subjects: Informed consent was obtained from the CEPH/Utah individuals, and the University of Utah Institutional Review Board approved the study (University of Utah IRB reference IRB_00065564).

