## Supplementary Information for "A familial, telomere-to-telomere reference for human *de novo* mutation and recombination from a four-generation pedigree"

### SUPPLEMENTARY NOTES

##### Y chromosome analysis

The highly contiguous Y assemblies generated here allowed us to investigate all classes of *de novo* mutations (DNMs) for the first time across the entire male-specific Y regions (MSY, i.e., excluding pseudoautosomal regions (PARs)), including the longest heterochromatic block in the human genome, the Yq12 subregion (**Fig. 5a**). Three Y lineages prevalent in populations with European ancestry (Jobling and Tyler-Smith 2003) are carried by the males of the 1463 pedigree: R1b1a-Z302 (n=9, G1-G4), R1b1a-Z326 (n=4, G3-G4), and I1a3a-S2078 (n=1, G1) (**Supplementary Fig. 42, Supplementary Table 13**).

Here, we focused on the nine-member male pedigree carrying the R1b1a-Z302 Y haplogroup (**Fig. 5a, Supplementary Table 13**). DNMs were identified from highly contiguous Verkko Y assemblies generated here for six out of nine males (G1-G3), with the longest Y contigs ranging from 23.1 to 51.2 Mbp. For three males (G1-NA12889, G2-NA12877 and G3-NA12884), representing three generations, the entire MSY (~48.8 Mbp) was contiguously assembled, with assembly breaks only in the PAR(s) (**Supplementary Table 13**). As the T2T-CHM13 chromosome Y reference (J1a-L816 Y lineage (Rhie et al. 2023)) differs extensively from the R1b1a-Z302 Y chromosomes, especially in the repetitive regions (**Supplementary Fig. 43**) (Hallast et al. 2023), we decided to use the G1-NA12889 chromosome Y assembly as a reference for DNM detection. The high quality of the G1-NA12889 Y assembly was confirmed by a comparison to an evolutionarily closely related Y from HG00731 (R1b1a-Z225 Y lineage, TMRCA estimate 5,700 years ago, 95% highest posterior density (HPD) interval = 4,800–6,700 years ago) and to the Y assemblies of his male descendants (**Fig. 5a, Supplementary Figs. 42 and 44**). Additionally, the G1-NA12889 Y assembly is highly similar in size and sequence composition to published assemblies from the R1b lineage Y chromosomes (**Supplementary Table 13**) (Hallast et al. 2023). The accessible *de novo* assembly-based MSY (i.e., excluding PARs and regions flagged as potential misassemblies by either Flagger or NucFreq) for variant calling for the six G1-G3 males ranged from 46.47–48.82 Mbp (mean 48.31 Mbp, median 48.60 Mbp) (**Supplementary Table 14**).

In total, we identified 48 *de novo* SNVs in the MSY across the five G2-G3 males. The majority (45/48) of identified SNVs were located in the Yq12 heterochromatic region, while two were in the Y euchromatic regions and one in the pericentromeric region (**Fig. 5b, Supplementary Table 14**). The average *de novo* SNV rate for euchromatic regions (X-degenerate, X-transposed and ampliconic) across the approximately 22.2 Mbp of accessible length was $1.81 \times 10^{-8}$ mutations per base per generation (95% CI = 0 - $4.89 \times 10^{-8}$, on average 0.4 SNVs/Y transmission), comparable to the *de novo* SNV rate estimates for the same regions from Icelandic patrilineages using Illumina data ($2.87 \times 10^{-8}$ mutations per base per generation, 95% CI = 2.68 - $3.08 \times 10^{-8}$) (Helgason et al. 2015). The *de novo* SNV rate estimate for the accessible ~23.4 Mbp of Yq12 ($3.86 \times 10^{-7}$ mutations per base per generation, 95% CI = 3.02 - $4.71 \times 10^{-7}$) is >20× higher compared to the euchromatic regions, an average of nine SNVs/Y transmission (**Supplementary Table 14**). However, it is worth noting that a substantial proportion of the SNVs in the highly repetitive Yq12 region might arise from mechanisms like interlocus gene conversion. In fact, 13/45 (29%) had 100% identical matches elsewhere in the respective individual's Y assembly (segments of ±200 bp of flanking sequences around the SNVs were used and the number of identical hits ranged from 2–1000, mean 149, median 3). However, this increased to 25/45 or 56% of Yq12 SNVs if flanking regions of ±25 bp around the SNVs were checked. Approximately 23.2/23.8 Mbp of the Yq12 region in the G1-NA12889 Y was annotated as composing of *DYZ1/Hsat3A6* and *DYZ2/Hsat1B* repeats (12.87 Mbp of *DYZ1* and 10.30 Mbp of *DYZ2* repeats), with their length ratio of 0.56/0.44. The SNVs identified in the Yq12 region follow a similar ratio of 0.52/0.48 (23/44 and 21/44) occurring in *DYZ1/DYZ2* repeats, respectively. The transition/transversion ratio across the 45 SNVs identified in the Yq12 is low, 0.73.

The average overall *de novo* SNV rate for the ~48.3 Mbp accessible MSY region was $1.99 \times 10^{-7}$ mutations per base per generation (95% CI = 1.59 - $2.39 \times 10^{-7}$) or 1.45 ×

$10^{-7}$ (95% CI = 1.12 to 1.79 × $10^{-7}$) if the 13 SNVs in Yq12 likely resulting from gene conversions were excluded (**Supplementary Table 14**).

A total of nine *de novo* indels (<50 bp, homopolymers excluded) were identified from the five males, ranging from 1-3 indels/sample (mean 1.8 events/Y transmission) (**Fig. 5b**). Overall, 8/9 were STRs (5/8 were 4-bp and 3/8 were 5-bp repeat expansions/ or contractions) while 1/9 was a 1 bp deletion (not in a homopolymer region); 5/9 indels were located in euchromatic regions (all 5 were validated by the Element data), while 4/9 were in the Yq12 regions where short-read data cannot be mapped unambiguously, overall translating to a mutation rate of 3.76 × $10^{-8}$ mutations per base per generation in the MSY (95% CI = 8.61 × $10^{-9}$ - 6.66 × $10^{-8}$) (**Supplementary Table 14**).

We also identified five *de novo* SVs (≥50 bp), ranging from 2,416 to 4,839 bp in size, an average of one SV per Y transmission (**Fig. 5b**). All identified SVs were located in the Yq12 and result in insertions or deletions of one or two entire *DYZ2* repeat subunits with an average mutation rate of 2.08 × $10^{-8}$ mutations per base per generation (95% CI = 0 - 5.20 × $10^{-8}$) (**Supplementary Table 14**). While it has only very recently become possible to investigate the genetic variation at the Yq12 region at base-pair resolution, a similar rate of approximately one SV at this region per approximately 40 Mbp scanned per meiosis was estimated to occur using restriction digestion followed by fractionation by pulse field gel electrophoresis by Neal Mathias (D.Phil thesis, Mathias, N., 1993, "Y chromosome DNA polymorphisms and human evolution", University of Oxford).

Due to assembly breaks in the PARs, these regions were not used for DNM detection. However, we aimed at locating the recombination breakpoints at the PARs, which is required for proper disjunction of X and Y chromosomes in male meiosis. No recombination events were identified from the Verkko assemblies in the PAR2s of the five R1b1a-Z302 Y males (one G2 and four G3), while one likely recombination event was present in the PAR1 of G3-NA12884 (**Supplementary Fig. 45**).

#### Mobile element insertion (MEI) analysis

Using xTea (v 0.1.9) to analyze the PacBio long reads, we identified non-reference MEI events. We classified all non-reference *Alu,* LINE-1, and SVA insertions. In G1-G3, we found 2161 full-length *Alu* insertions (**Supplementary Fig. 14a**), 398 LINE-1 insertions, and 151 SVA insertions (**Supplementary Fig. 14b**). We identified 112 LINE-1 insertions that were either full-length or near full-length (at least 5500 bp).

To examine the subfamilies of the non-reference MEIs, the sequence for each *Alu* element greater than 240 bp in length was aligned using muscle (v.3.8.31). Genetic similarities among all possible pairs of sequences were calculated, and principal components were obtained by eigen decomposition of the similarity matrix. The relationship between known *Alu* element subfamilies and the non-reference *Alu* element insertion sequences is shown in **Supplementary Figure 46a**. Most of the non-reference *Alu* insertions cluster closely around known *AluYa* and *AluYb* subfamily sequences. This process was repeated for full-length LINE-1 insertions (**Supplementary Fig. 46b**) and SVA insertions (**Supplementary Fig. 46c**). As expected most non-reference LINE-1 insertions cluster around the L1HS subfamily, and most SVA insertions cluster around the SVAE and SVAF subfamilies. Using BLAT, we identified LINE-1 insertions in the reference genome that were most likely to be source elements for each of our non-reference, full-length LINE-1 insertions. For 101 of the 112 insertions, we were able to identify the most likely source element in the reference genome; 79 of these non-reference insertions can be traced back to 20 LINE-1s in the T2T-CHM13 reference genome. The source elements responsible for more than one non-reference insertion and the location of the non-reference insertions are shown in **Supplementary Fig. 46d**. A LINE-1 insertion on chromosome 17 is the most likely source element for 12 of these non-reference insertions, and another on chromosome 16 is the most likely source element for nine non-reference insertions. These findings support the hypothesis that there are only a small number of active LINE-1 loci in each individual. Using BEDTools, we intersected the non-reference MEIs with the 5'- and 3'-UTRs (untranslated regions), introns, and exons of known protein-coding genes in T2T-CHM13. For *Alu* elements, LINE-1, and SVA, most of the insertions fall in intergenic regions. Of those that do overlap genic regions, the majority of the insertions intersect introns, with a small number that fall into 5'- or 3'-UTRs. One *Alu* element retrotransposed into an exon of *PRAMEF4*, a gene involved in cell proliferation, apoptosis, and transcription (**Supplementary Fig. 46e**).

### SUPPLEMENTARY TABLES

**All supplementary tables are available as individual .xlsx files.**

### SUPPLEMENTARY FIGURES

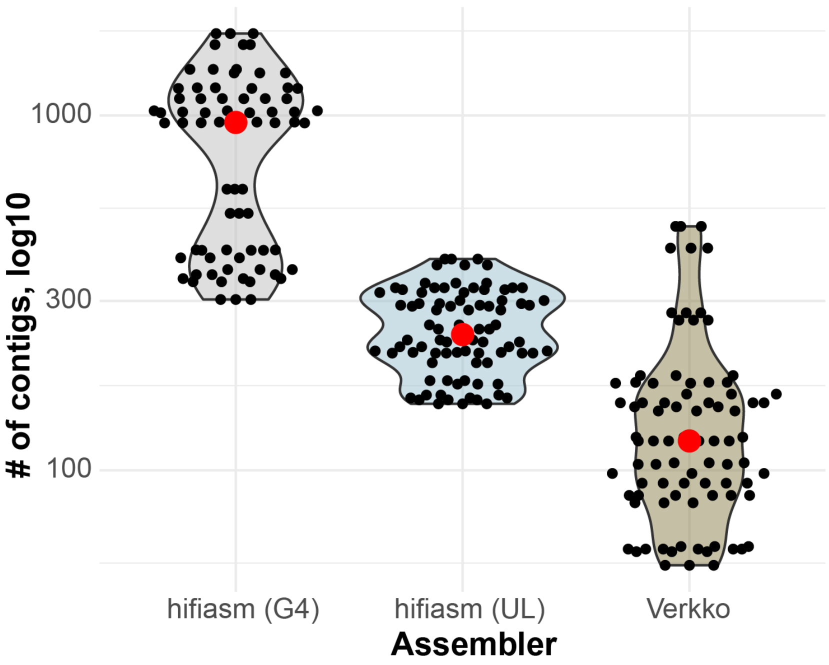

**Supplementary Figure 1: Evaluation of assembly contiguity.**
Distribution of the total contig counts assembled using Verkko (brown), hifiasm (UL) (light blue), and assemblies of G4 samples using hifiasm (light gray).

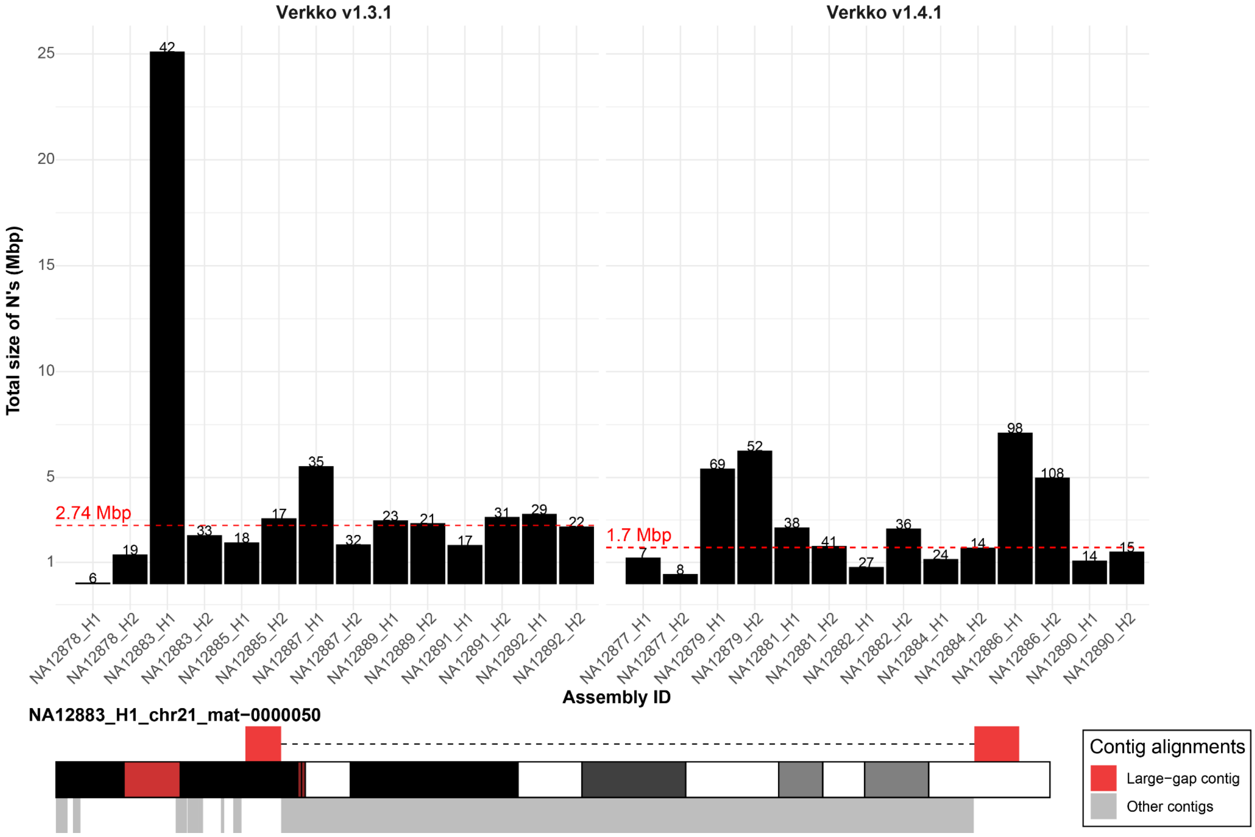

#### Supplementary Figure 2: Evaluation of gaps in scaffolded Verkko assemblies.

**Top**: A barplot showing total length of N's detected in scaffolds reported by Verkko. On top of each bar is the number of separate strings of N's detected in each phased Verkko assembly. Median length of all N's strings in each assembly is shown as a horizontal red dashed line separately for Verkko versions 1.3.1 and 1.4.1. **Bottom**: Explanation for an observed outlier of excess on N's in NA12883_H1 assembly caused by a single contig that maps at opposite ends of chromosome 21 where space in between was filled by string of N's by Verkko. This is likely a bug in the Verkko (v1.3.1) pipeline as this contig was supposed to be divided into two different contigs.

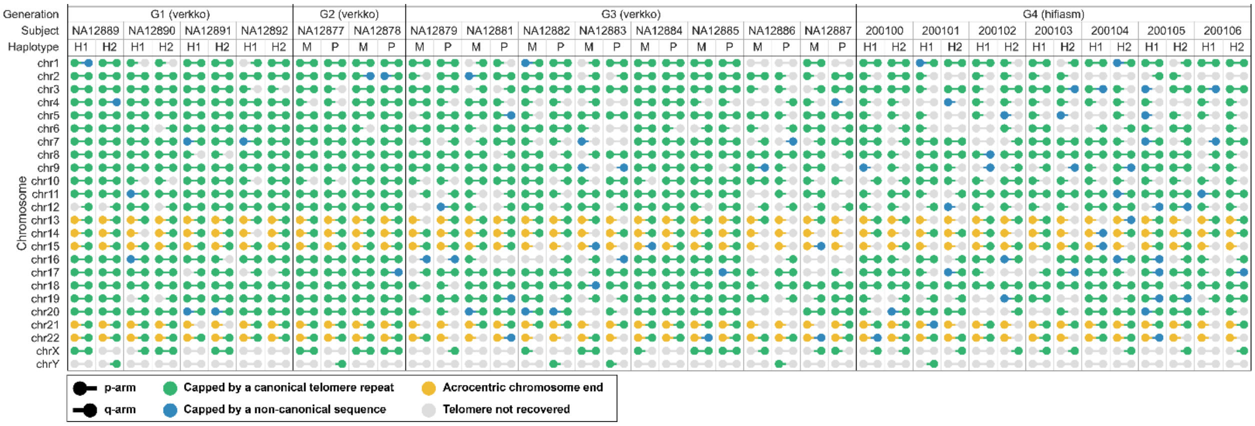

#### Supplementary Figure 3: Telomere completeness in phased genome assemblies.

Evaluation of telomere recovery across G1-G4 assemblies and all chromosomes. *p* and *q* arms are marked separately. See **Extended Data Figure 1d** for the visualization of the chromosomes that are spanned by a single T2T contig in Verkko assemblies for G1-G3; out of all hifiasm assemblies for G4, only chr9 of 200101 haplotype 1 is verifiably contiguous due to the absence of ultra-long ONT reads that have not been generated for the G4 samples.

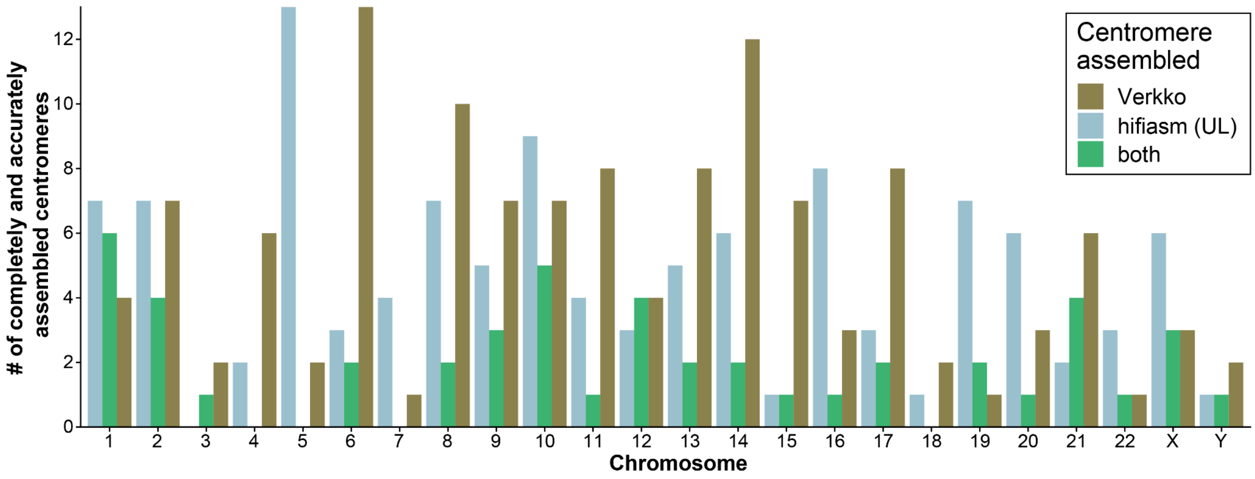

#### Supplementary Figure 4: Centromere completeness in phased genome assemblies.

A barplot showing the number of completely and accurately assembled centromeres by Verkko (brown), hifiasm (UL) (light blue), or both assemblers (green). This plot highlights certain chromosomes (e.g., 5, 6, 17 and 19) as being preferentially assembled by different assembly algorithms.

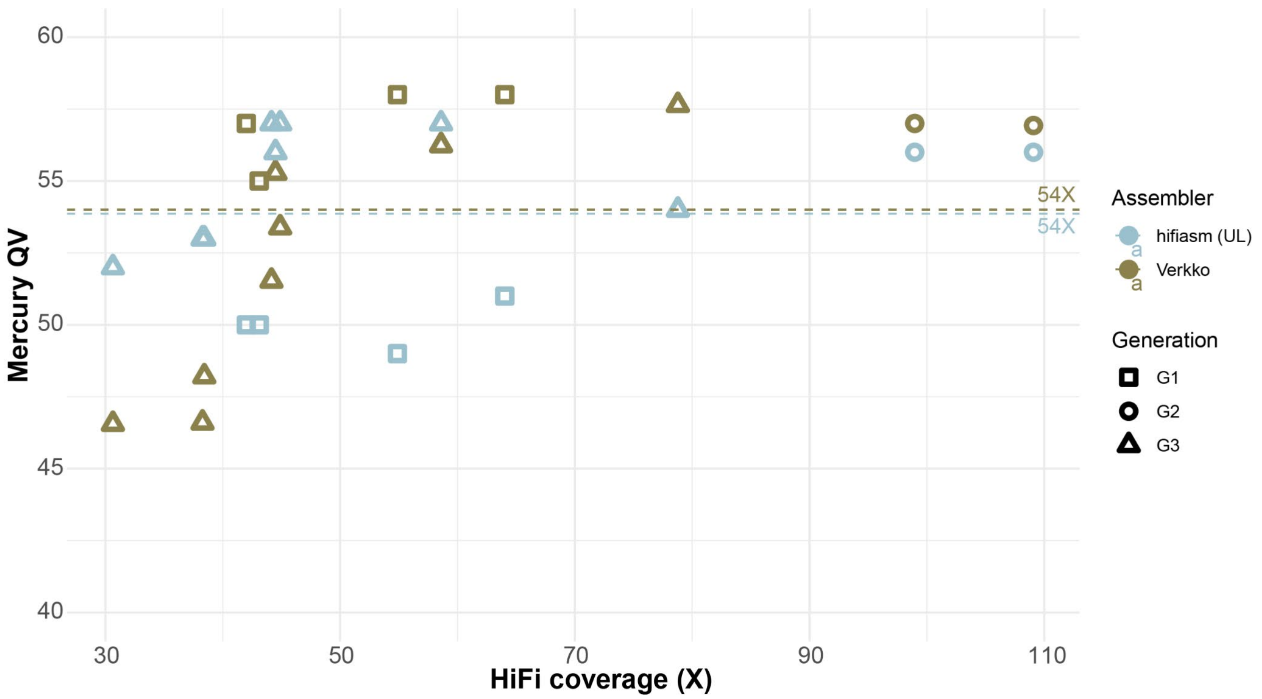

#### Supplementary Figure 5: Evaluation of assembly quality.

Distribution of assembly quality values (QV) for assemblies of G2 and G3 using either Verkko (brown) or hifiasm (UL) (light blue) assembler. Mean QV across haploid assemblies for a given assembler is shown as horizontal dashed line.

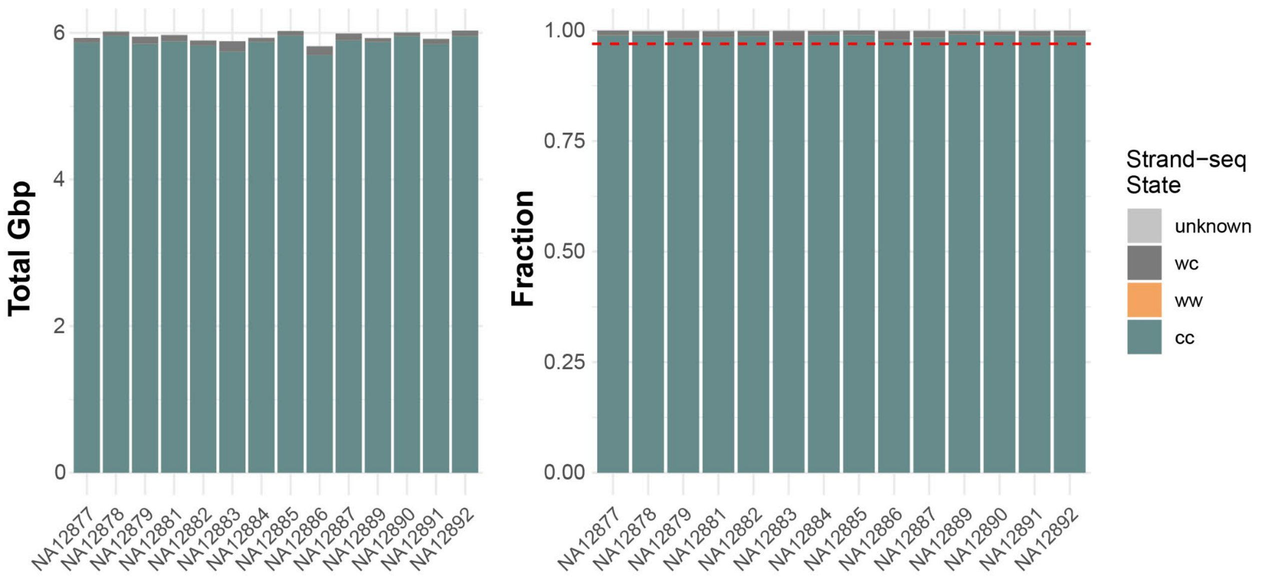

#### Supplementary Figure 6: Evaluation of misoriented regions with Strand-seq.

**Left:** Total number of base pairs (per diploid genome) genotyped as ‘cc’ which means they agree with Strand-seq reference orientation. On the other hand, regions genotyped as ‘ww’ point to possible misorientations. Regions genotyped as ‘wc’ are either caused by the presence of heterozygous inversions or low mappability regions for short Strand-seq reads. ‘Unknown’ region could not be reliably genotyped, most likely due to spurious mapping of short Strand-seq reads. **Right:** The same as the left plot but shown as a fraction of bases of each Strand-seq state. Red horizontal dashed line marks 0.97 value.

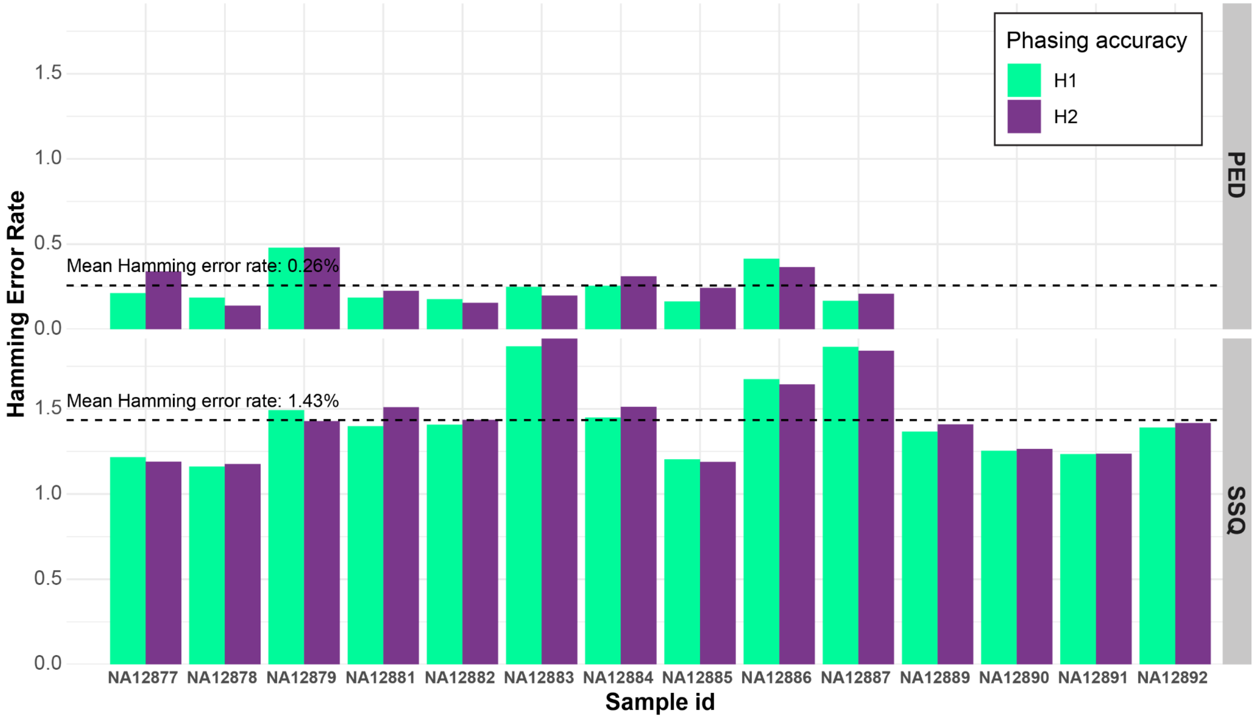

**Supplementary Figure 7: Evaluation of phasing accuracy using Strand-seq.**

A barplot showing the level of phasing disagreement between pedigree-phased genome assemblies (PED) and Strand-seq (SSQ) phasing. Phased variants are compared with respect to the GRCh38 reference. Hamming error rate is reported separately for haplotype1 (H1 - green) and haplotype2 (H2 - purple) for each sample from G2-G3.

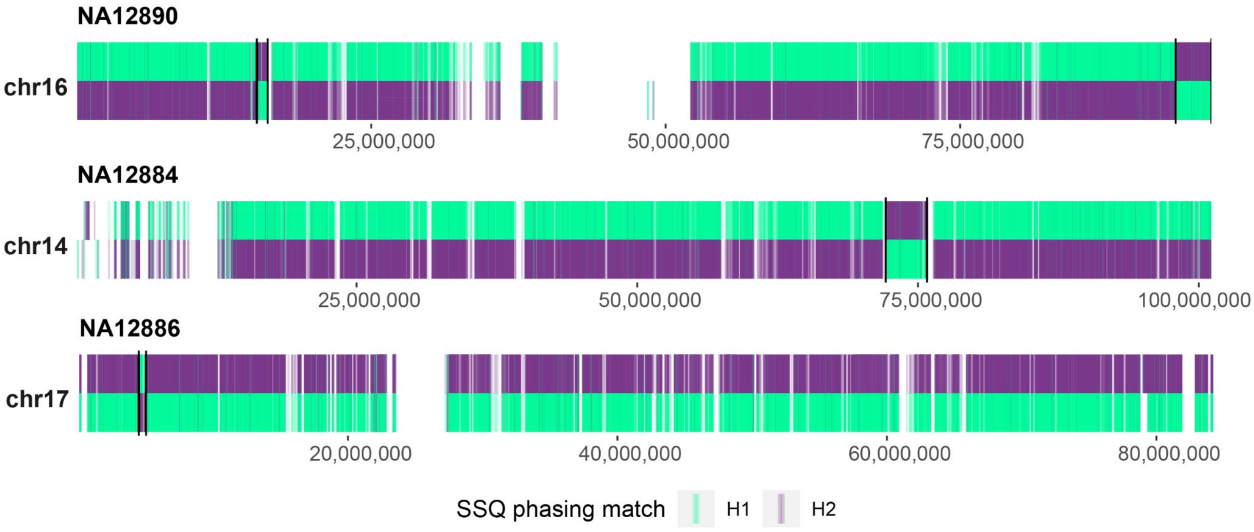

#### Supplementary Figure 8: Phasing errors identified by Strand-seq.

A distribution of heterozygous single-nucleotide variants (SNVs) along three chromosomes (14, 16 and 17) where large-scale haplotype switch errors were detected in Verkko assemblies. Here, variants detected with respect to the T2T-CHM13 reference are compared between assembly-based SNV callset (obtained from PAV) and Strand-seq (SSQ)-based phasing. Assembly-based phased SNV calls are compared to Strand-seq-based phasing and each variant is colored green if it matches haplotype 1 (H1) or haplotype 2 (H2) of the Strand-seq-based phasing. Extended phasing switch errors are visible as regions where H1 changes to match H2 or vice versa (**Supplementary Table 4**).

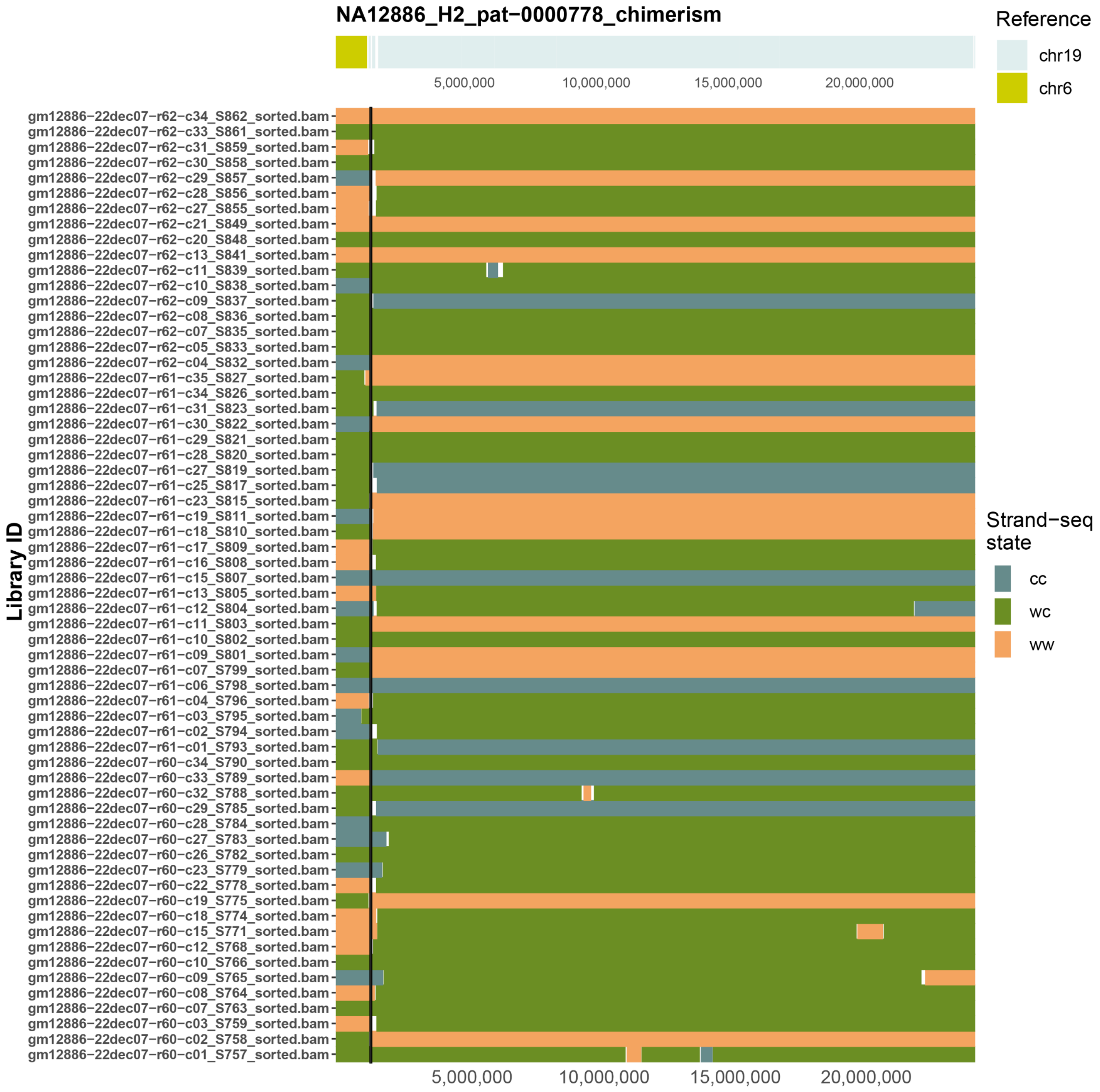

**Supplementary Figure 9: Chimeric contig in Verkko assembly.**

**Top**: Partial mapping of NA12886 paternal contig ‘pat-0000778’ to chromosomes 6 and 19 of the T2T-CHM13 reference. **Bottom**: Horizontal colored bars show Strand-seq strand states for individual single-cell libraries. Each region is genotyped as WC: Watson-Crick (green); WW: Watson-Watson (orange); CC: Crick-Crick (blue). Observed recurrent strand-state change is indicative of a genome misassembly. Recurrent strand-state change in a paternal contig in sample NA12886 points to contig chimerism as parts of the contig can be aligned to both chromosomes 6 and 19. The point of recurrent strand-state change supporting misassembly is highlighted by the vertical line. All chimeric contigs detected in Verkko or hifiasm assemblies are reported in **Supplementary Table 4**.

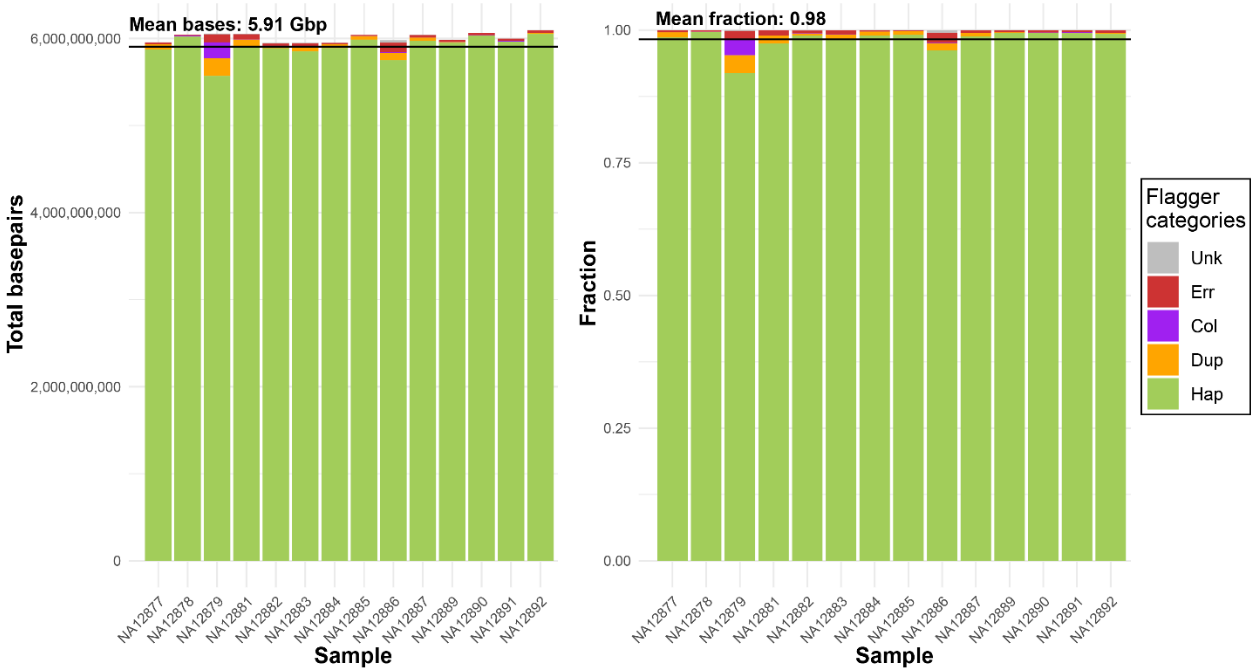

#### Supplementary Figure 10: Flagger summary of possible problematic regions in phased assemblies.

Flagger categorizes the genome assembly into segments based on the mapping of HiFi (high-fidelity) reads back to the assembly. A short definition of these categories follows: **Err** (Erroneous, red) A low-read coverage segment that could be either a misjoin or a region that needs polishing. **Dup** (Duplicated, orange) Likely a false duplication of another region in the genome. **Hap** (Haploid, green) A genomic segment that is correctly assembled and has the expected read coverage. **Col** (Collapsed, purple) Two or more highly similar haplotypes are collapsed into this block. **Unk** (Unknown, gray) A segment that could not be confidently assigned.

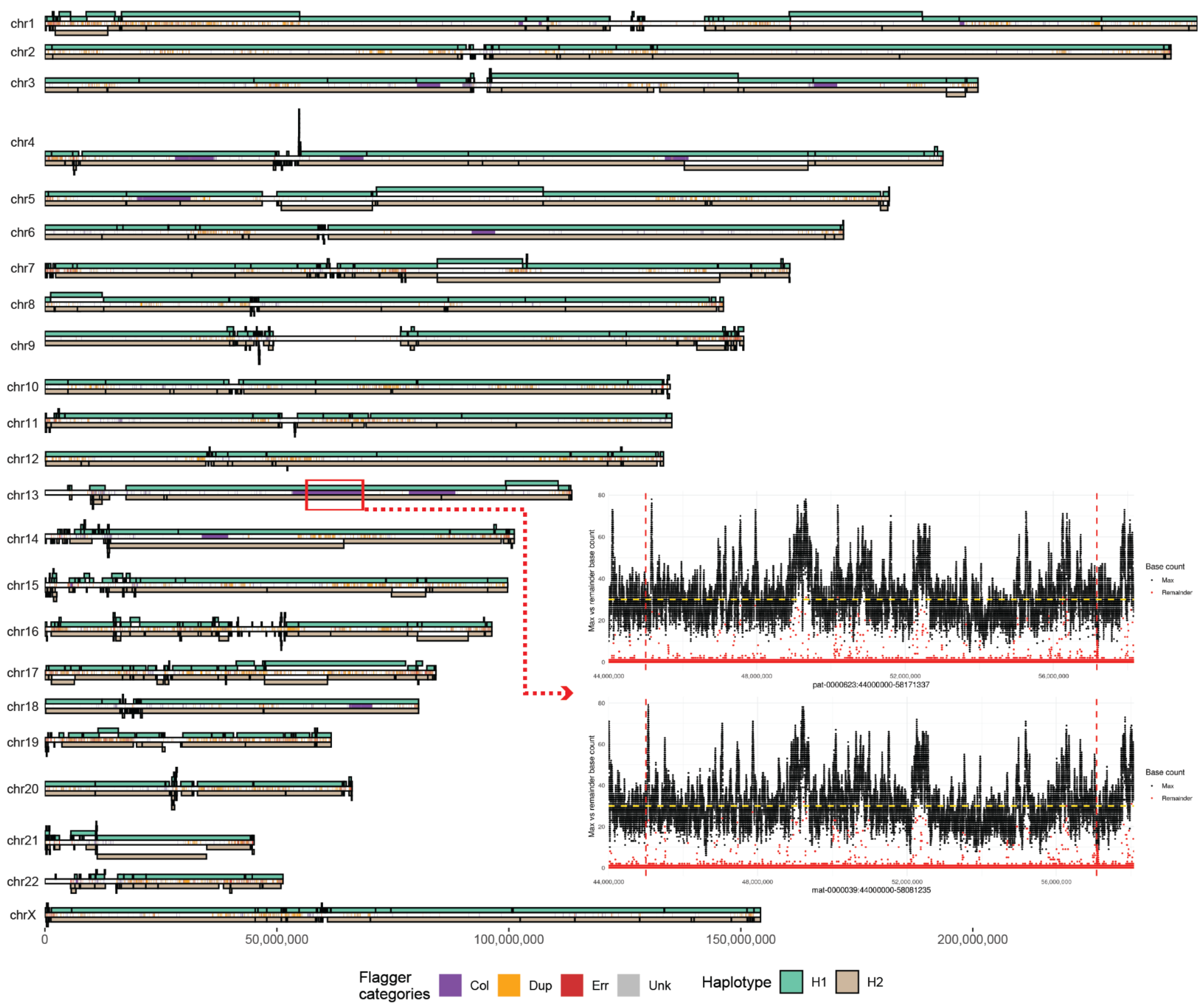

#### Supplementary Figure 11: Flagger evaluation of NA12879.

An ideogram showing alignments of phased assembly to the T2T-CHM13 reference with haplotype 1 contigs (H1 - light green) being shown above the chromosome midline and haplotype 2 (H2 - light brown) contigs below. Positions of regions reported by Flagger as collapsed (‘Col’ - purple), duplicated (‘Dup’ - orange) or erroneous (‘Err’ - red) (see **Supplementary Figure 10** legend for more details) are shown at the midline of each chromosome. **Inset:** A NucFreq visualization of ~12 Mbp region (red rectangle highlight in top plot) reported as collapsed by Flagger for both paternal and maternal haplotype assembly. The horizontal yellow dashed line shows median coverage and vertical red lines show region boundaries reported by Flagger. Black dots show the count of the most abundant base (‘max’) at a given position while red dots show the count of the second most frequent base (‘remainder’). Low frequencies of the second most frequent base along the whole region suggest that a given region is correctly assembled and observed coverage fluctuations inherent to the HiFi data analyzed here.

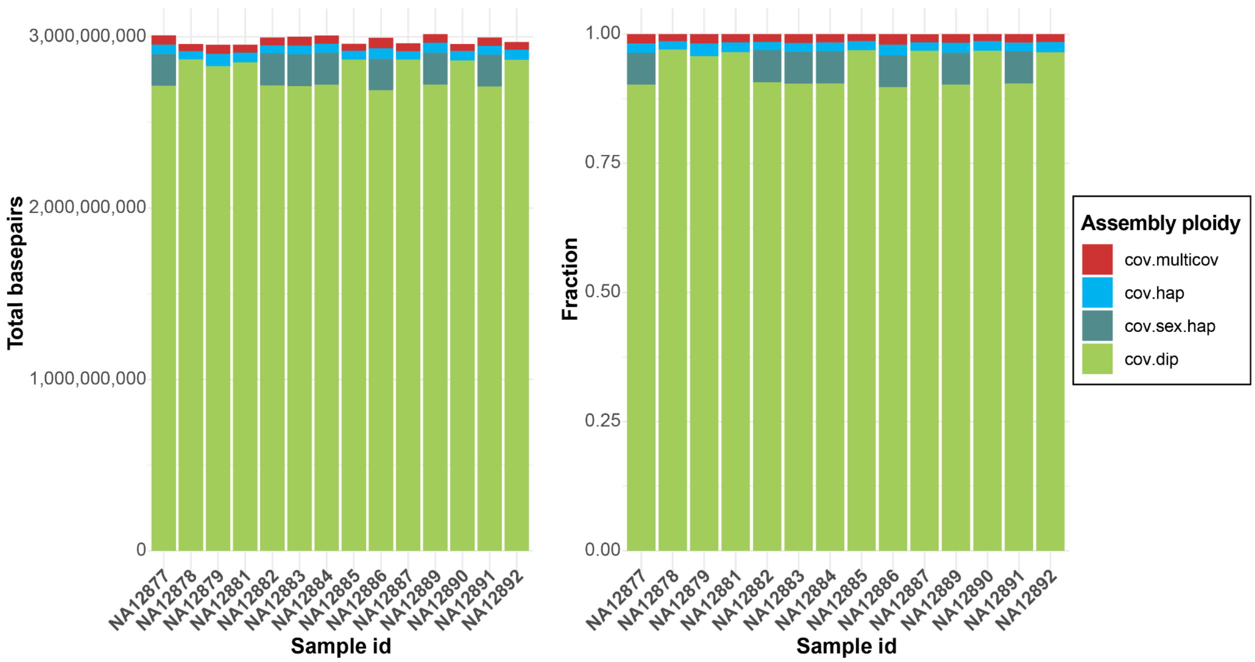

**Supplementary Figure 12: Ploidy summary of aligned Verkko genome assemblies to T2T-CHM13 reference.**

**Left**: A stacked barplot showing the proportion of a number of base pairs reported as having exactly single alignment per haploid assembly, thus being a stable diploid region ('cov.dip' - green). In males, sex chromosomes are expected to be single copy, thus having a single alignment ('cov.sex.hap' - tiel). Regions that have reported more than a single alignment per haploid assembly are marked as multi-coverage regions ('cov.multicov' - red) while those that are missing an alignment in either haplotype are marked as haploid ('cov.hap' - light blue).

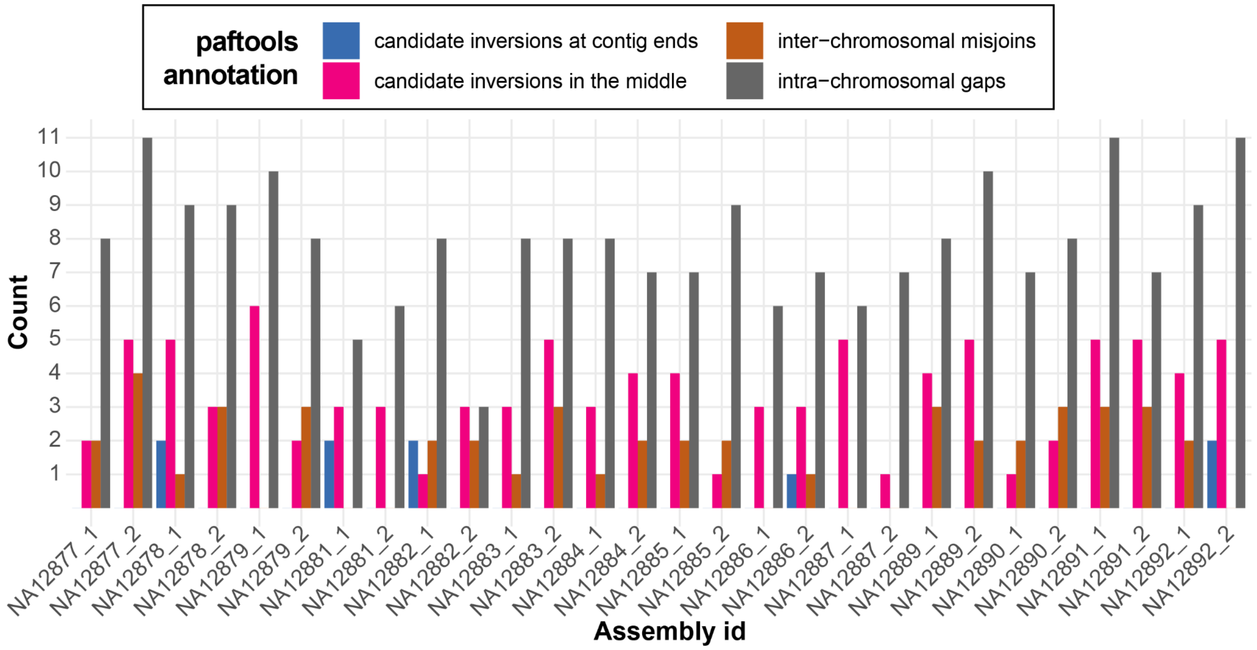

**Supplementary Figure 13: Detected assembly issues with paftools.**

A barplot showing the counts of assembly issues detected by paftools 'misjoin' functionality per haploid assembly.

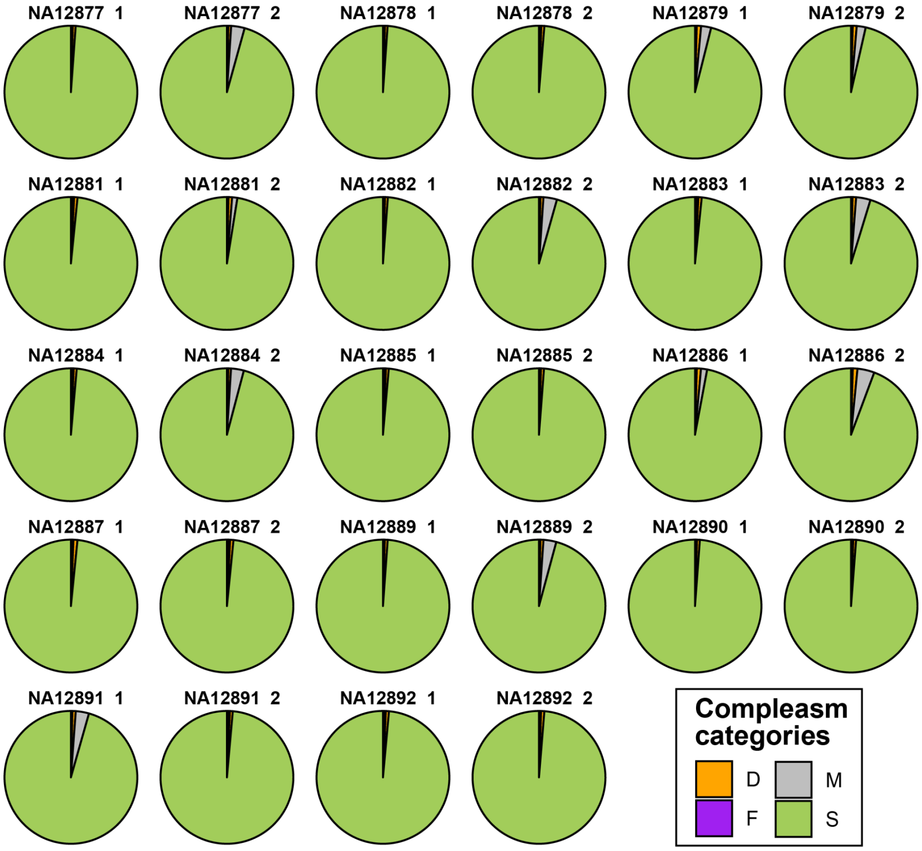

**Supplementary Figure 14: Gene completeness assessment using compleasm.**
Each pie chart shows a fraction of correctly assembled single-copy genes for each phased assembly (1 - haplotype1, 2 - haplotype 2). See below for an explanation of each compleasm (miniBusco) category:

- S (Single-Copy Complete Genes): The BUSCO genes that can be entirely aligned in the assembly, with only one copy present.
- D (Duplicated Complete Genes): The BUSCO genes that can be completely aligned in the assembly, with more than one copy present.
- F (Fragmented Genes, subclass 1): The BUSCO genes where only a portion of the gene is present in the assembly, and the rest of the gene cannot be aligned.
- M (Missing Genes): The BUSCO genes with no alignment present in the assembly.

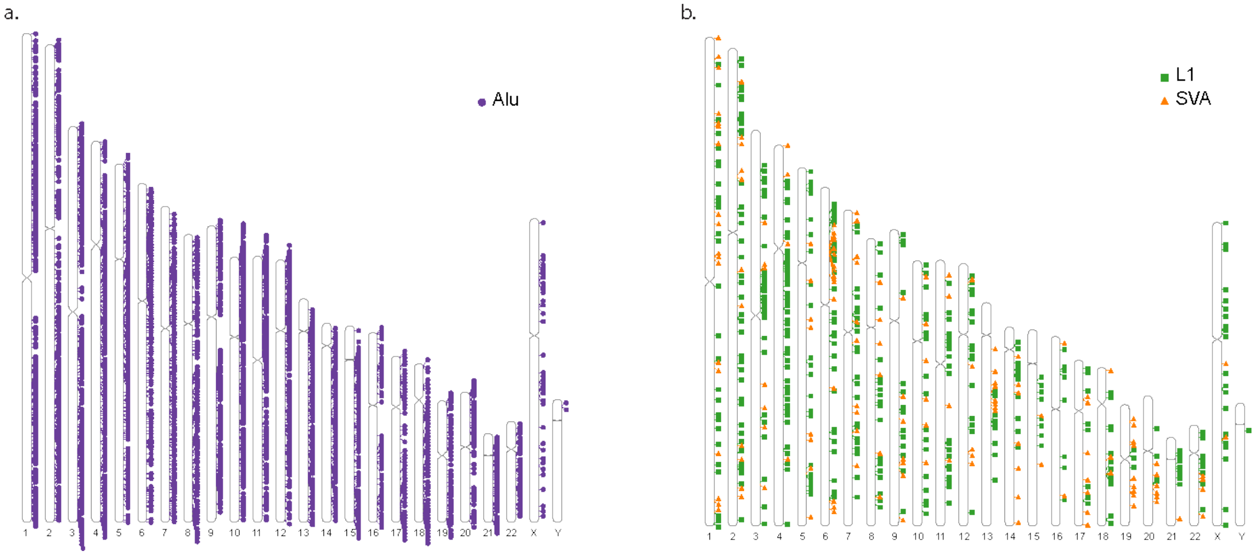

**Supplementary Figure 15: Non-reference mobile element insertion (MEI) analysis.**
**a**) Ideogram of non-reference *Alu* element insertions in T2T-CHM13. **b**) Ideogram of non-reference LINE-1 and SVA insertions in T2T-CHM13. These plots include both full-length and shorter MEI events.

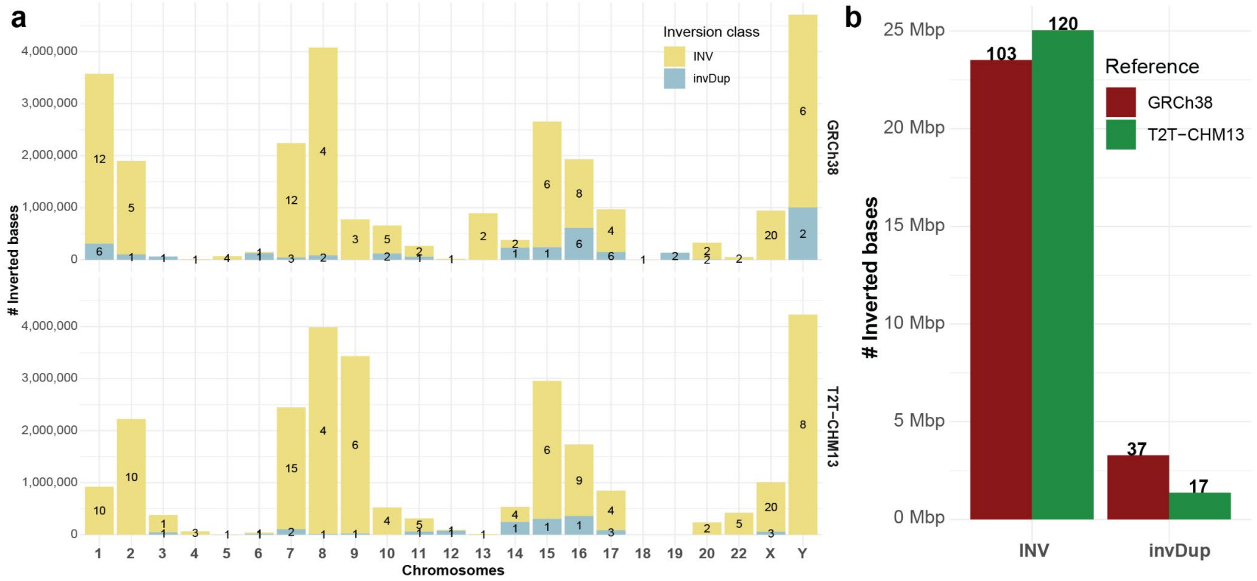

**Supplementary Figure 16: Strand-seq-based inversion callset.**
**a**) A barplot showing the total number of inverted bases with respect to a reference genome (top - GRCh38, bottom - T2T-CHM13). The number in the middle of each stacked bar reports the count of nonredundant inversions contributing to the total count of inverted bases. Inversion calls are stratified based on their class (simple inversions - INV, inverted duplications - invDup). **b**) A barplot showing the total number of simple inversions (INV) and inverted duplications (invDup) with respect to a reference genome (GRCh38 - red, T2T-CHM13 - green).

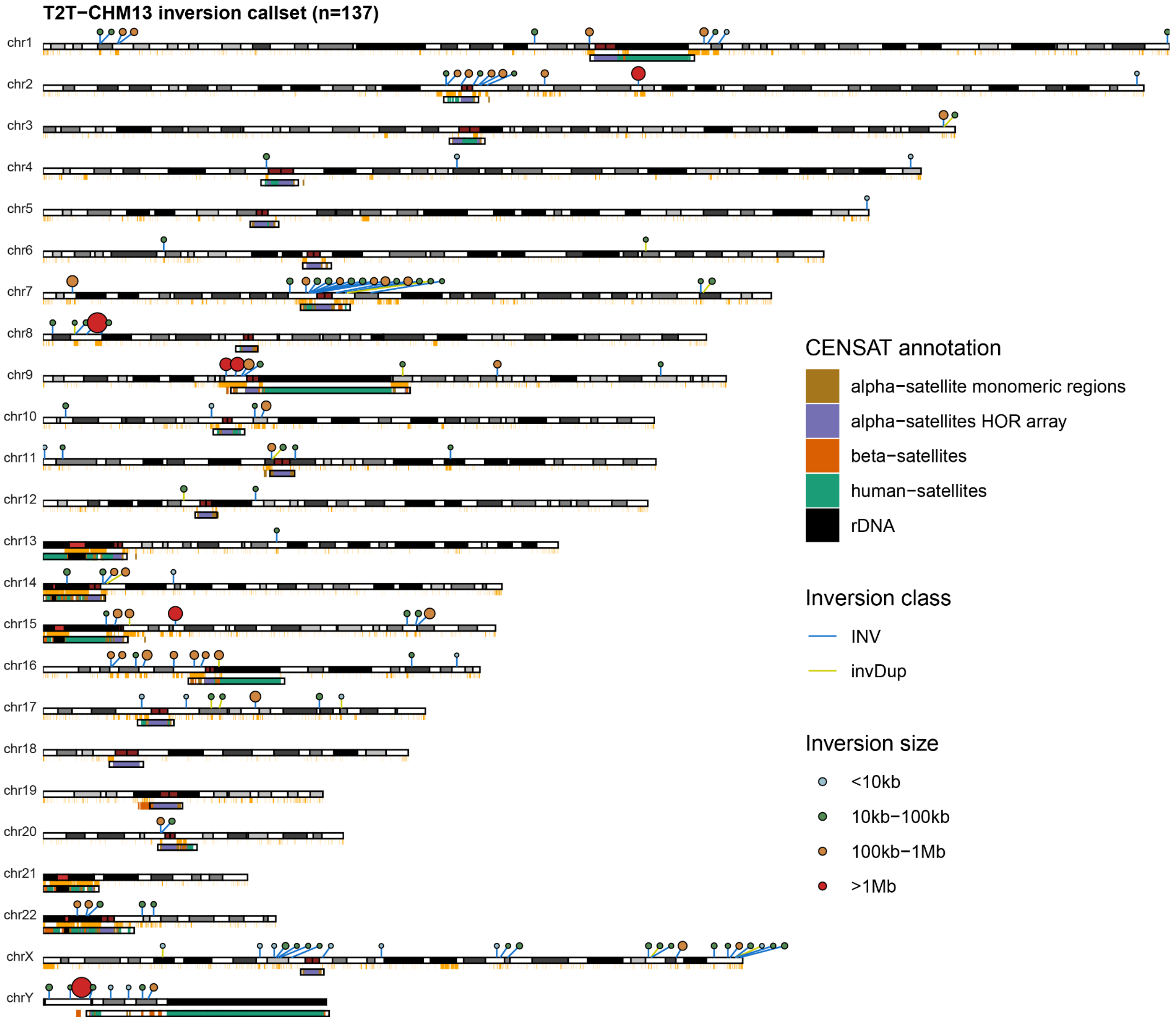

**Supplementary Figure 17: Strand-seq inversion callset with respect to T2T-CHM13.** An ideogram showing the distribution of simple inversions (INV - blue link) and inverted duplications (invDup - yellow link) with respect to the T2T-CHM13 reference. The size and color of each dot reflects the inversion size category. Below each chromosome ideogram there is a segmental duplication (SD) annotation (orange) and centromeric satellite annotation (CENSAT).

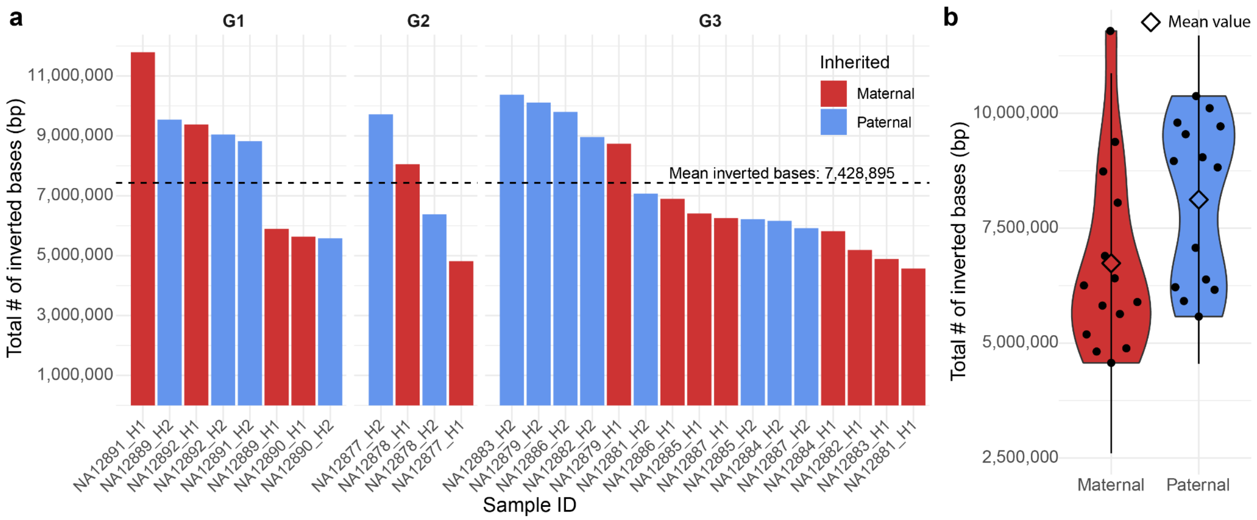

**Supplementary Figure 18: Inverted bases per haploid genome and per generation.**
**a**) A barplot showing the total number of inverted bases per maternal (H1 - maternal; red) and paternal (H2 - paternal; blue) haplotype across G1-G3 samples. The dashed line shows the mean number of inverted bases across all samples and haplotypes. **b**) A violin plot showing the distribution of inverted bases per maternal (H1 - maternal; red) and paternal (H2 - paternal; blue) homolog. The diamond point indicates the mean value for each distribution. There are on average 6.7 Mbp and 8.1 Mbp of inverted bases per maternal and paternal homologs, respectively.

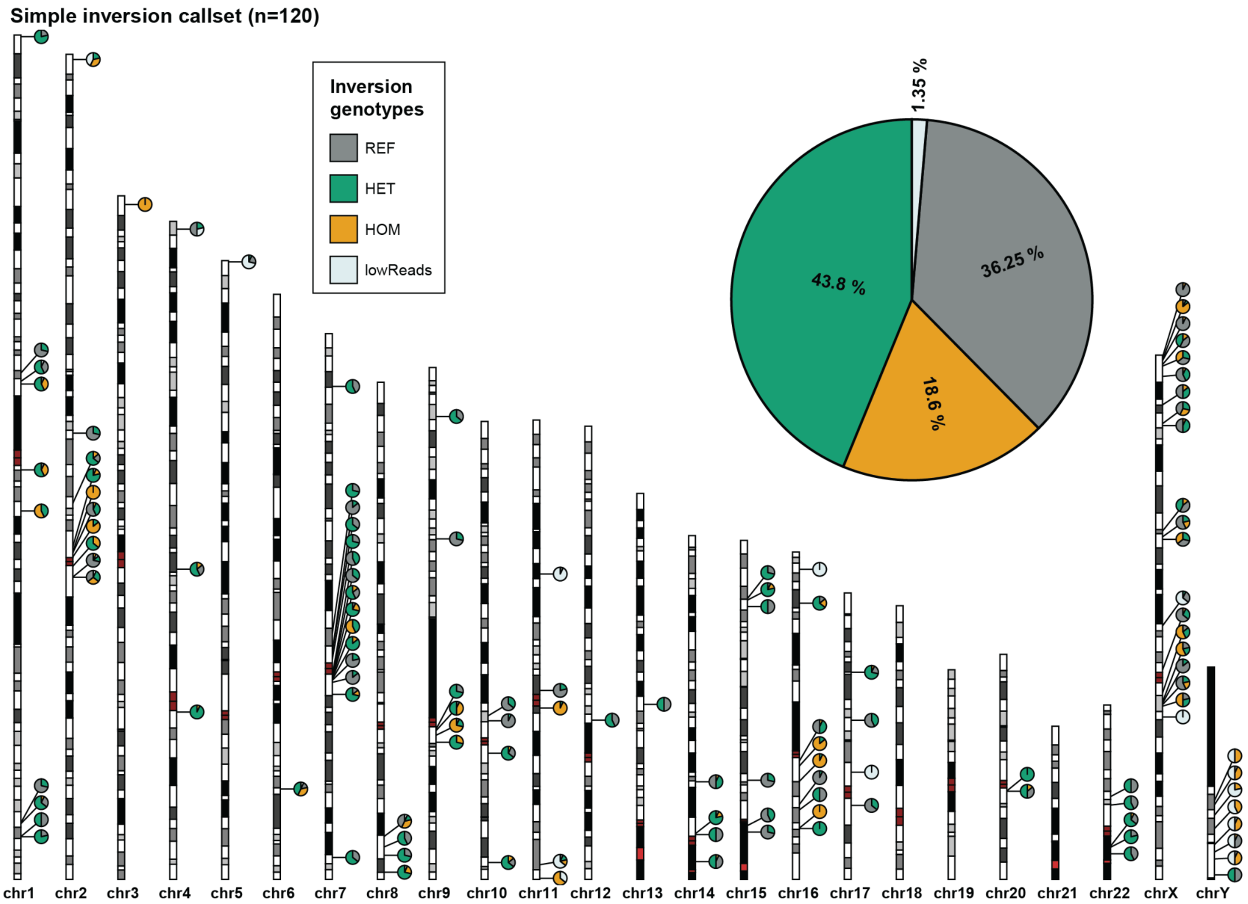

**Supplementary Figure 19: Summary of Strand-seq inversion genotypes.**

An ideogram showing the inversion genotype (‘REF’ - reference orientation, ‘HET’ - heterozygous inversion, ‘HOM’ - homozygous inversion, and ‘lowReads’ - not enough Strand-seq reads to genotype) proportions across G1-G3 for simple inversions (n=120). Each pie chart represents a single inverted region. The large inset pie chart shows the overall proportion of homozygous and heterozygous inversions in this family.

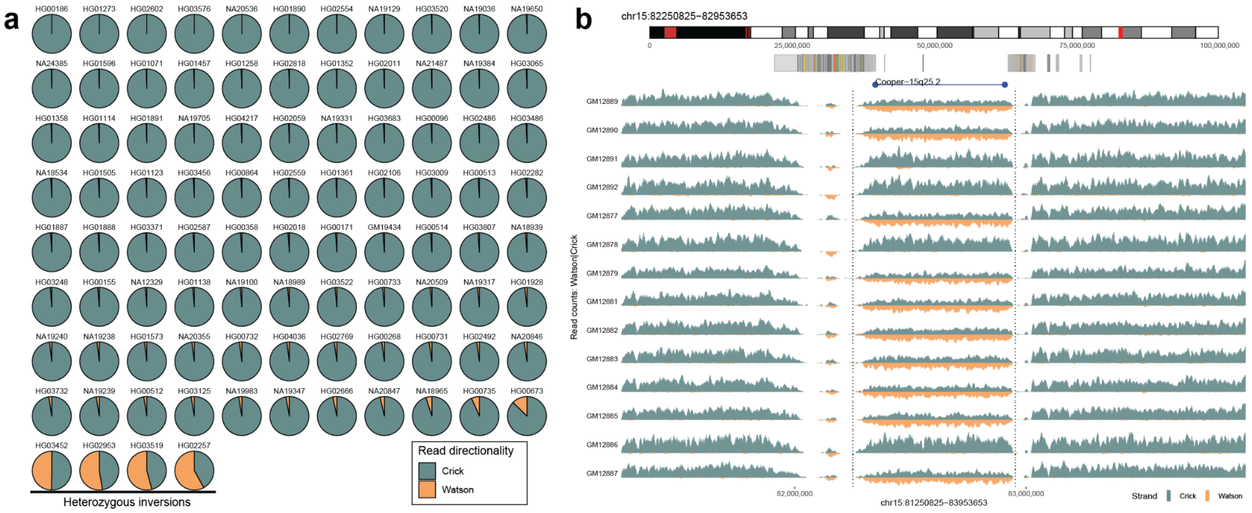

**Supplementary Figure 20: Rare inversion at 15q25.2-25.3.**

**a**) A multiple pie chart plot showing the proportions of Crick (plus read - teal) and Watson (minus reads - orange) aligned to the region of interest across 1000 Genomes Project samples (n=92). **b**) The read-coverage profiles of Strand-seq data over the region of interest summarized as binned (bin size: 10 kbp, step size: 1 kbp) read counts represented as bars above (teal; Crick read counts) and below (orange; Watson read counts) the midline. Dotted lines highlight the inverted region with respect to T2T-CHM13. In this region, equal coverage of Watson and Crick counts represents a heterozygous inversion as only one homologue is inverted with respect to the reference while reads aligned only in Watson orientation represent a homozygous inversion. Above is the chromosome ideogram with the region of interest highlighted in red followed by SD annotation and morbid copy number variant region for Cooper syndrome.

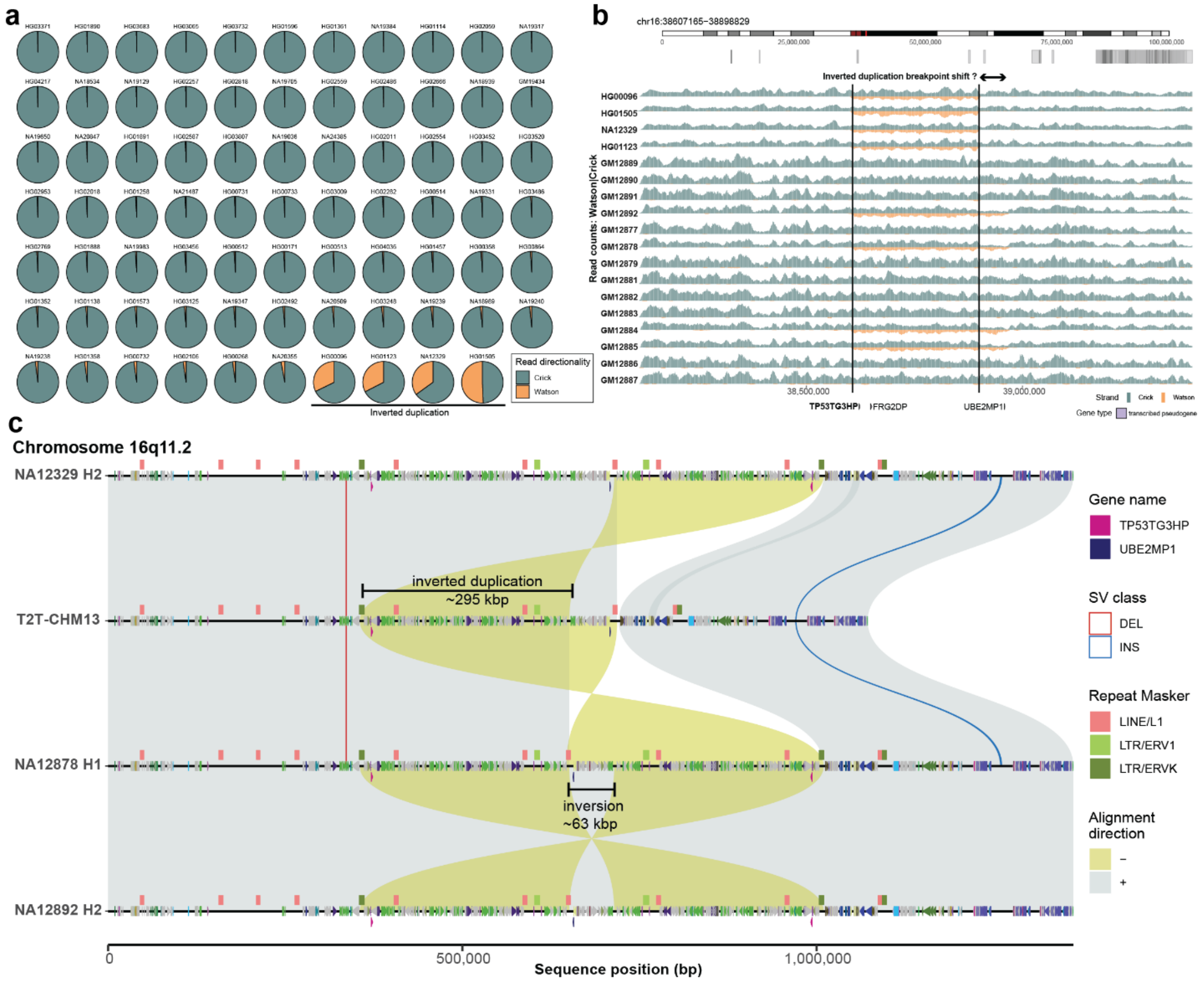

#### Supplementary Figure 21: Rare, inverted duplication at 16q11.2.

**a**) A multiple pie chart plot showing the proportions of Crick (plus read - teal) and Watson (minus reads - orange) aligned to the region of interest across 1000 Genomes Project samples (n=92). **b**) The read-coverage profiles of Strand-seq data over the region of interest (T2T-CHM13; chr16:38250451-39322284) summarized as binned (bin size: 10 kbp, step size: 1 kbp) read counts represented as bars above (teal; Crick read counts) and below (orange; Watson read counts) the midline. Vertical solid lines highlight the duplicated and inverted regions with respect to T2T-CHM13 supported by roughly equal coverage of Watson and Crick read counts with respect to the reference without this inverted duplication. Above is the chromosome ideogram with the region of interest highlighted in red followed by SD annotation. **c**) A visualization of syntenic relationships between three fully assembled haplotypes (NA12329-H2, G2-NA12878-H1, and G1-12892-H2) and the T2T-CHM13 reference based on minimap2 alignments. Direct (+, forward) alignments are shown in gray and reverse (-) alignments in yellow. On top of each haplotype we show the DupMasker (Jiang et al. 2008) annotation as colored arrowheads pointing forward or backward for direct and reverse oriented segments, respectively. Also, there is a RepeatMasker annotation. Structural variants (SVs; ≥50 bp) are shown as red (DEL - deletion) and blue (INS - insertion) lines between aligned haplotypes.

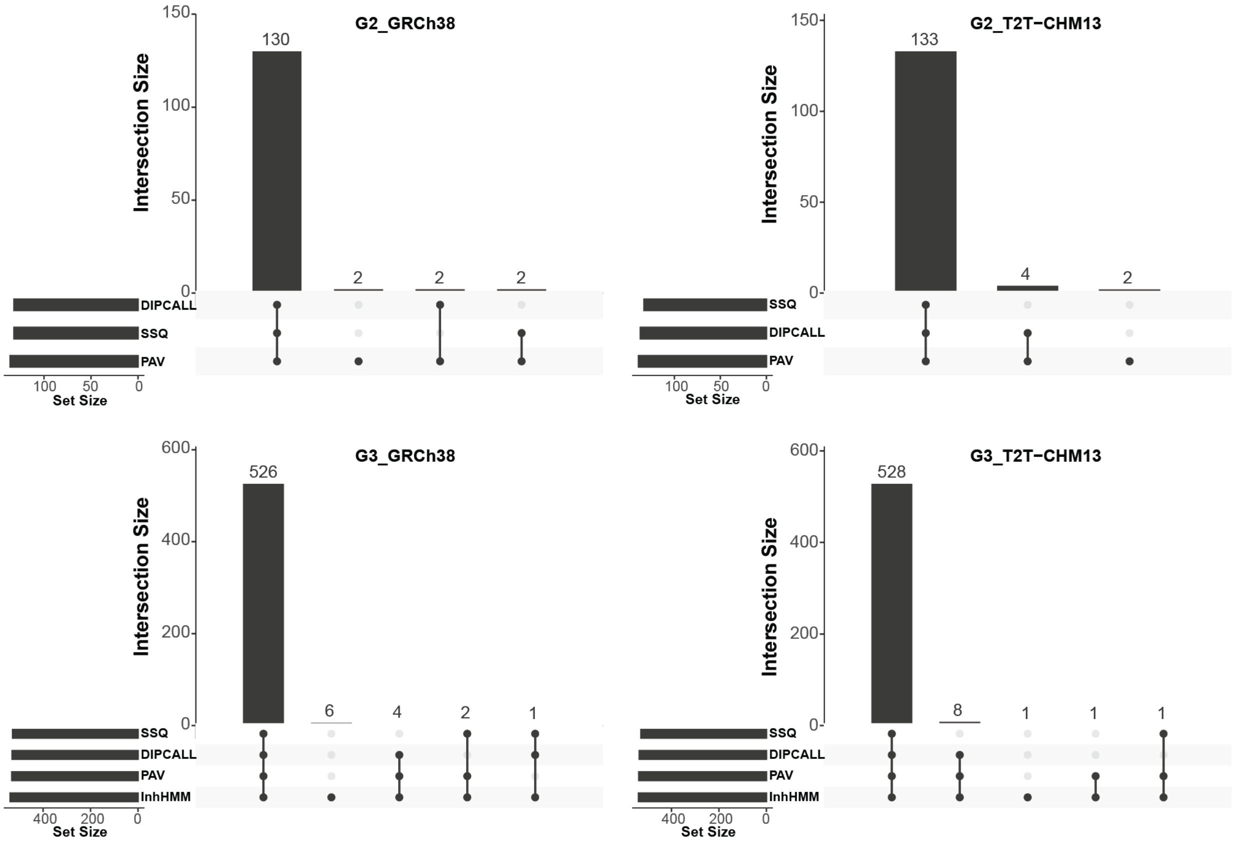

#### Supplementary Figure 22: Recombination breakpoint overlap.

Each upset plot summarizes an overlap between the reference recombination map with respect to supporting orthogonal datasets. In the case of G3, the reference recombination map is based on inheritance vectors (InhHMM) while supporting datasets are based on maps defined with the help of phased genome assemblies (PAV, Dipcall) and Strand-seq (SSQ). In the case of G2, the reference recombination map is based on phased genome assemblies (specifically based on PAV) while supporting datasets are based on maps defined with help of phased genome assemblies (Dipcall) and Strand-seq (SSQ).

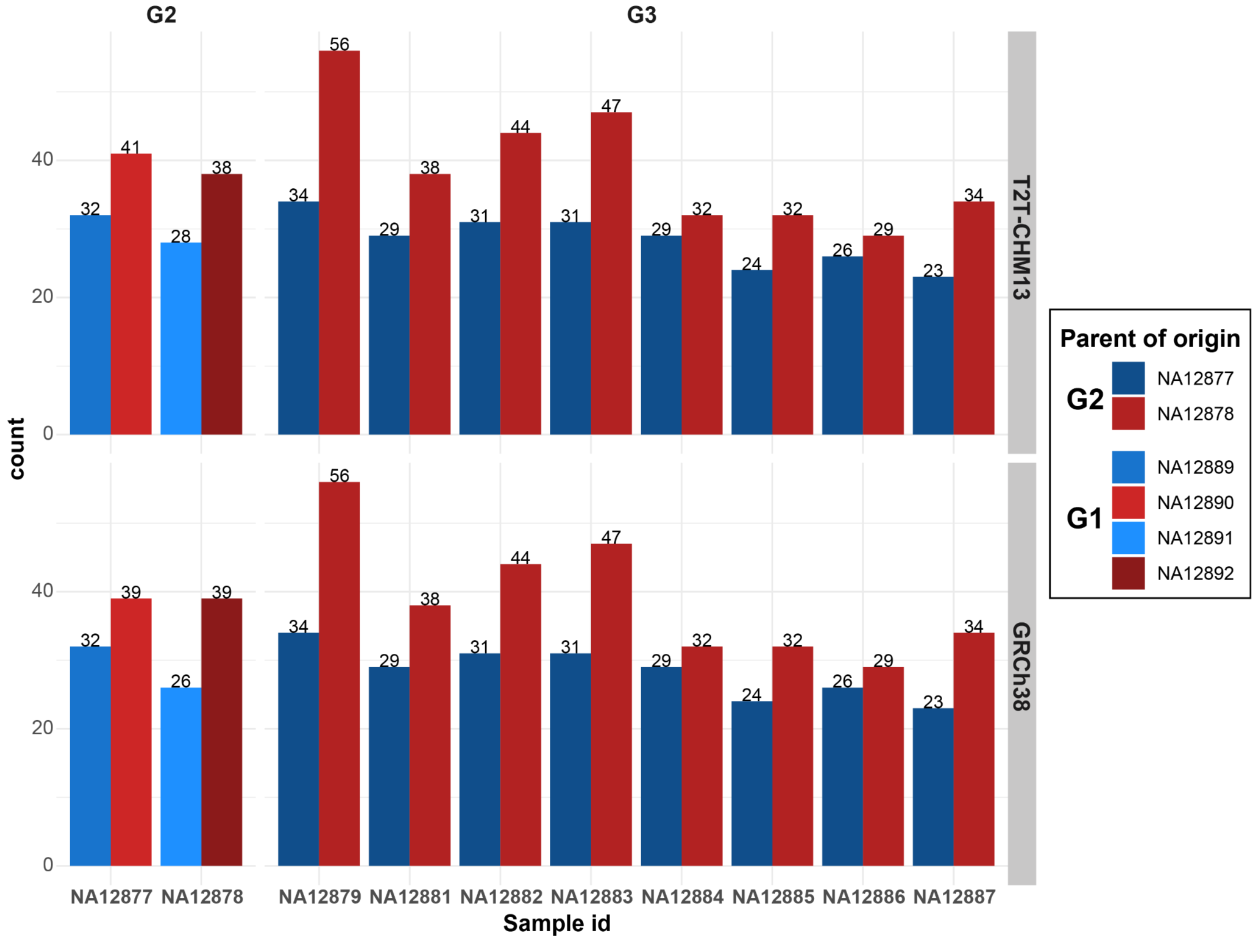

#### Supplementary Figure 23: Male and female recombination breakpoints per sample.

A barplot showing the total number of recombination breakpoints detected in each G2 and G3 sample colored by parent of origin of inherited homologs (red colors - maternal, blue colors - paternal). There are a total of 678 and 675 breakpoints with respect to T2T-CHM13 (top) and GRCh38 (bottom) reference genome, respectively (**Supplementary Table 8**).

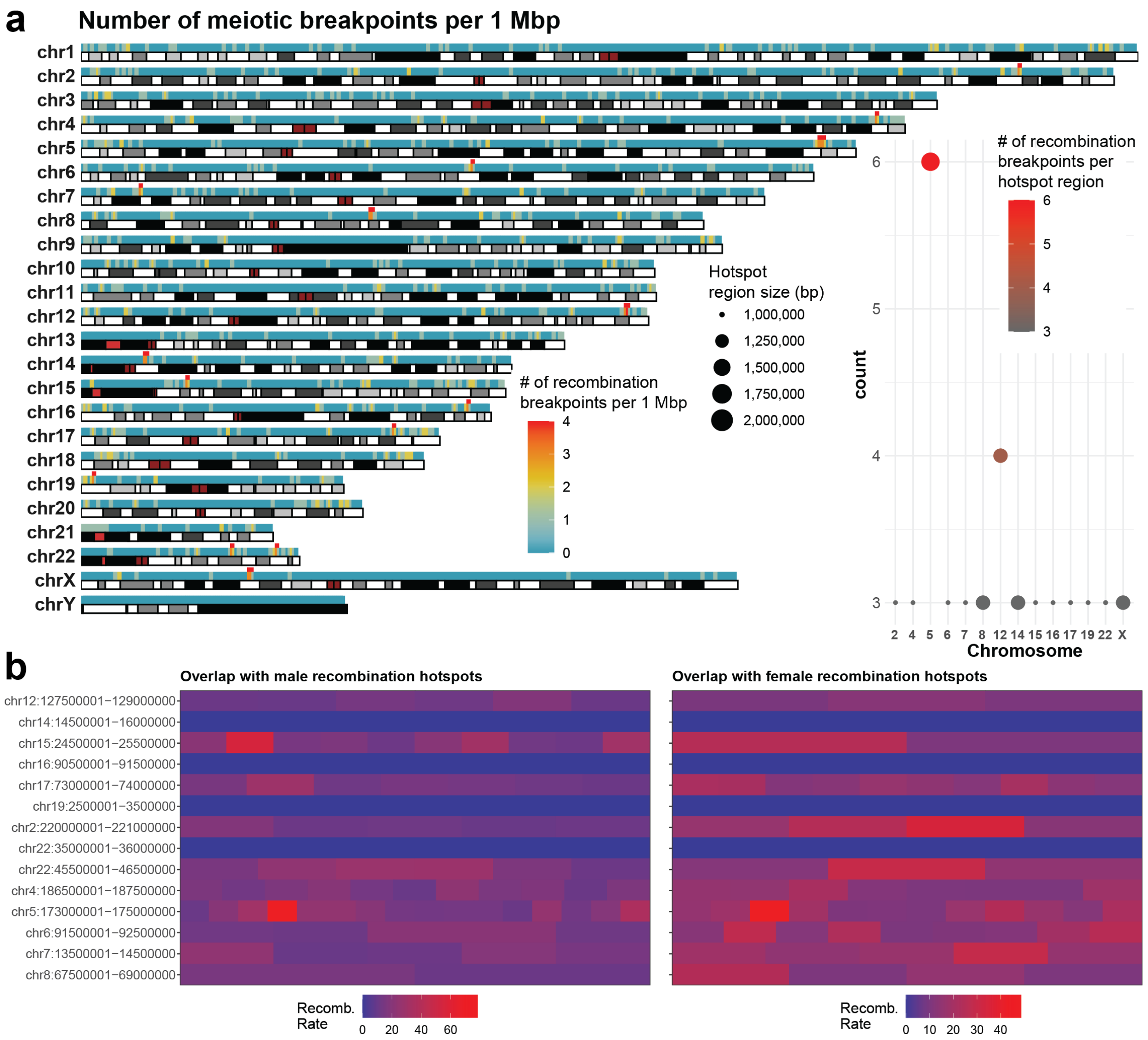

#### Supplementary Figure 24: Recombination breakpoint hotspots (T2T-CHM13).

**a**) An ideogram showing the binned counts of all recombination breakpoints in G2-G3 (n=678) summarized in 1 Mbp long bins sliding by 500 kbp as a heatmap on top of each chromosomal ideogram. Regions with breakpoint count ≥3 were selected as putative recombination breakpoints (n=15) (**Supplementary Table 8**). These are highlighted on top of each chromosomal heatmap as red rectangles. **Inset:** A summary of putative recombination hotspots per chromosome (x-axis) where the color of each point represents the number of meiotic breakpoints in a given region while the size of each point represents the size of the region. Note: Maternal and paternal meiotic breakpoints were not separated in this analysis but rather analyzed as a single set of recombination breakpoints. **b**) A comparison of the overlap of the 14 predicted autosomal recombination hotspots reported for this pedigree with male and female meiotic recombination hotspots from the deCODE project (Kong et al. 2010). Shades of red and blue rectangle depict hot and cold recombination clusters from the deCODE project, respectively.

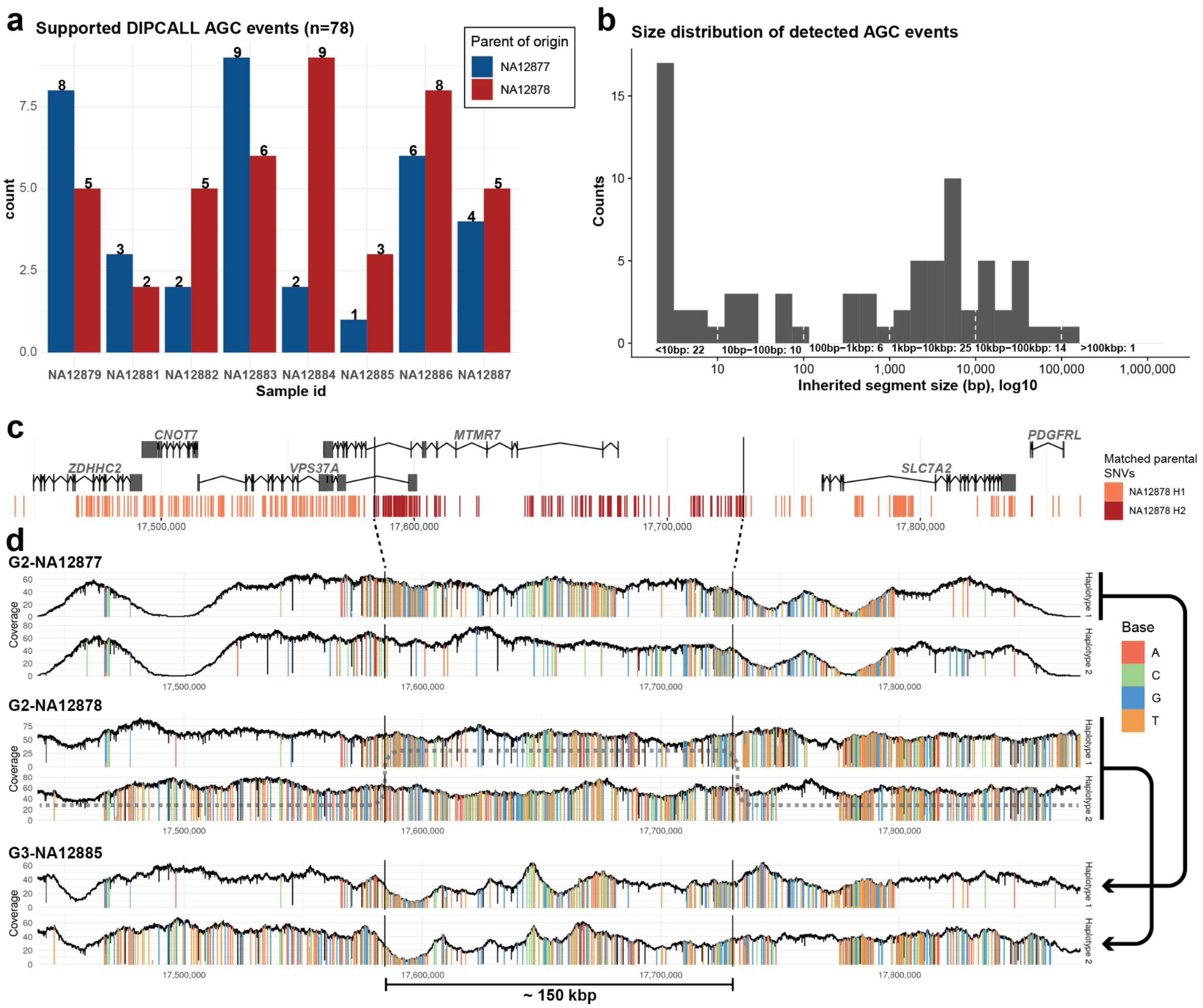

#### Supplementary Figure 25: Putative allelic gene conversion events (T2T-CHM13).

**a**) A barplot showing the total number of allelic gene conversion (AGC) events detected in G2 and G3 samples stratified by parental identity of inherited homologs (red - maternal, blue - paternal). **b**) Size distribution of observed AGC events from panel a. **c**) Overview of protein-coding genes overlapping an example AGC region on chromosome 8 (chr8:17584336-17730132). Below there are visualized SNVs from sample G3-NA12885 colored based on their match to the inherited parental (G2-NA12878) haplotype 1 (light red) or haplotype 2 (dark red). **d**) A mismatch pattern of phased HiFi reads with respect to the T2T-CHM13 reference for both haplotype from parent 1 (G2-NA12877), parent 2 (G2-NA12878), and a G3 sample (NA12885). Arrows on the right side of the plot show what homolog from the parent was inherited based on the matching pattern of observed mismatches. The child's haplotype two is composed of a mixture of alleles from NA12877, which is in line with allelic gene conversion or eventually two double-strand breaks and resolved as cross-over.

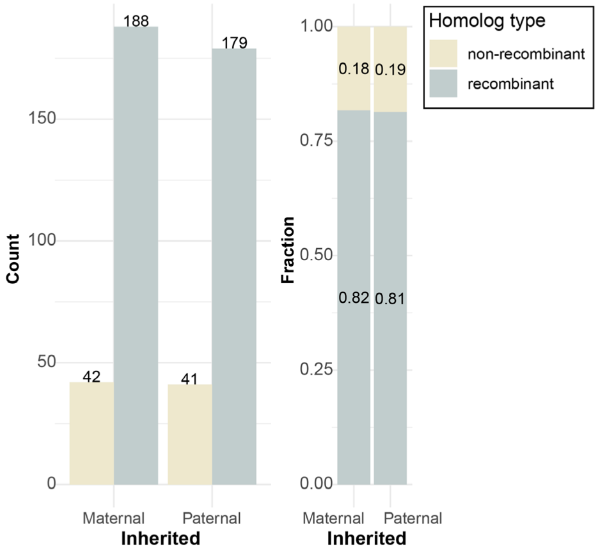

**Supplementary Figure 26: Summary of observed recombinant and nonrecombinant parental alleles.**

**Left:** Counts of recombinant and nonrecombinant homologs shown separately for maternal and paternal homologs. **Right**: Fraction of recombinant and nonrecombinant homologs calculated based on counts from the left plot. Note: Male chromosome X is not considered here.

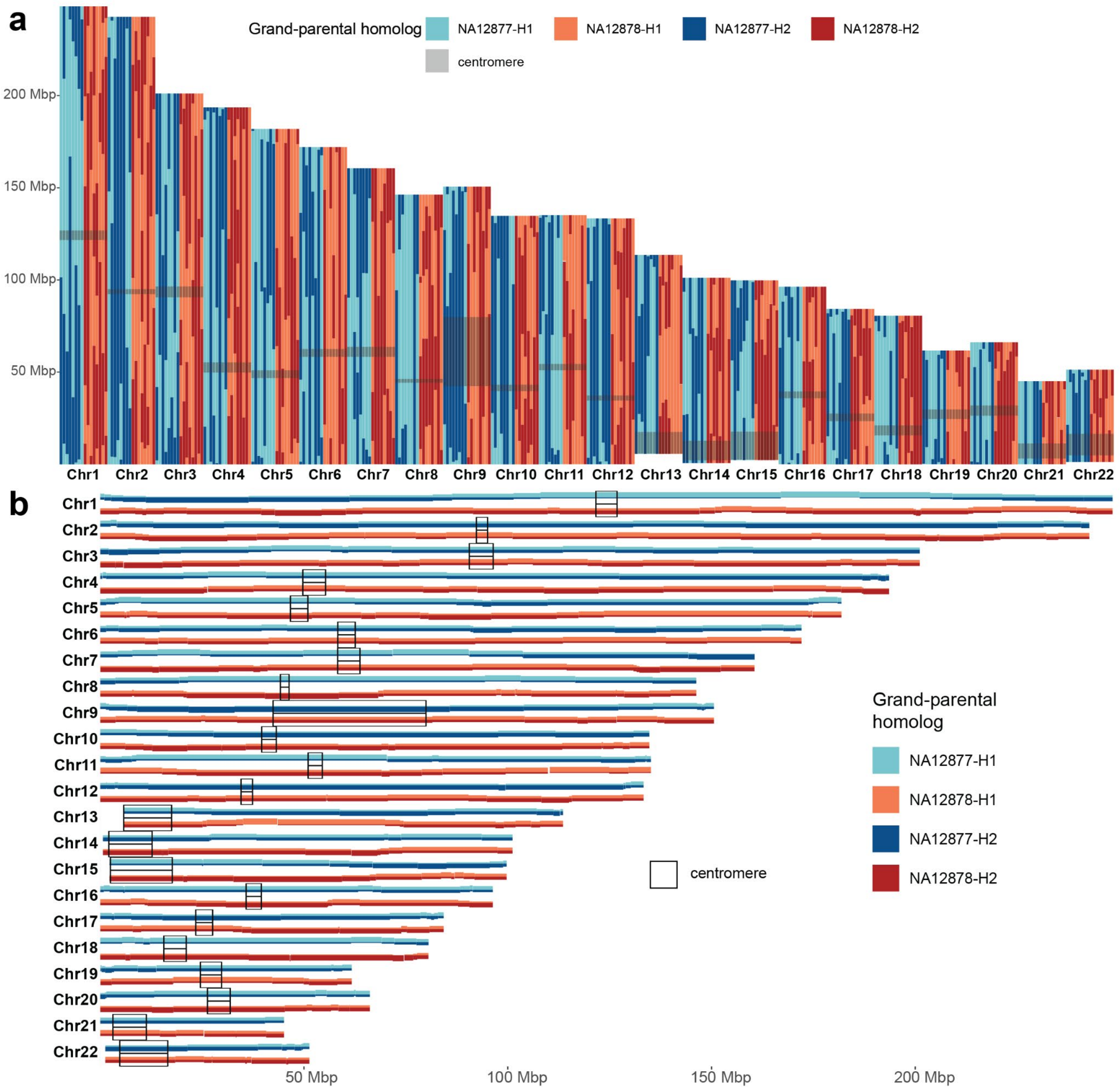

**Supplementary Figure 27: Inheritance of parental segments in G3 samples (T2T-CHM13).**

**a**) Recombination map for all G3 samples (n=8) for all chromosomes. **b**) A collapsed version of the recombination map from above shows the frequency (0 to 8) as to how many times for each genomic segment is inherited from either paternal or maternal haplotype (H1 or H2). Black boxes highlight the positions of centromeres within each chromosome.

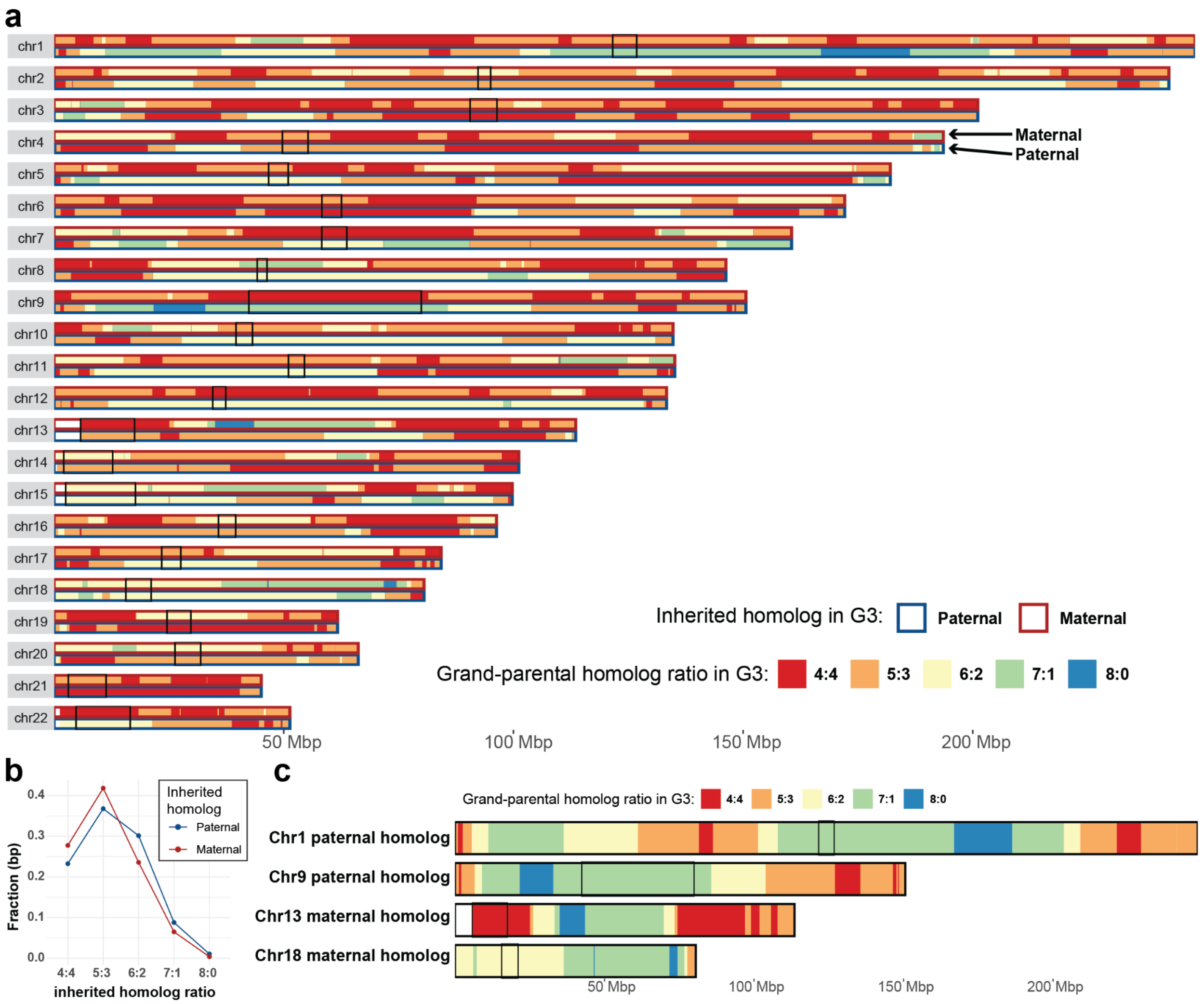

#### Supplementary Figure 28: Inheritance of parental segments in G3 samples (T2T-CHM13).

**a**) Each chromosome is segmented into nonoverlapping segments with specific ratios of maternal (top row) and paternal (bottom row) segments across all G3 samples (n=8). There are five expected ratios (4:4, 5:3/3:5, 6:2/2:6, 7:1/1:7, 8:0,0:8) of inherited parental segments. Centromeric regions are highlighted by black boxes. **b**) A proportion of base pairs belonging to five expected parental segment ratios stratified by homolog of origin (maternal/NA12878 - red, paternal/NA12877 - blue). **c**) Chromosomes that carry regions (blue) inherited from a single grandparental segment from G2. Each color represents a ratio of grandparental homologs inherited in G3 (n=8).

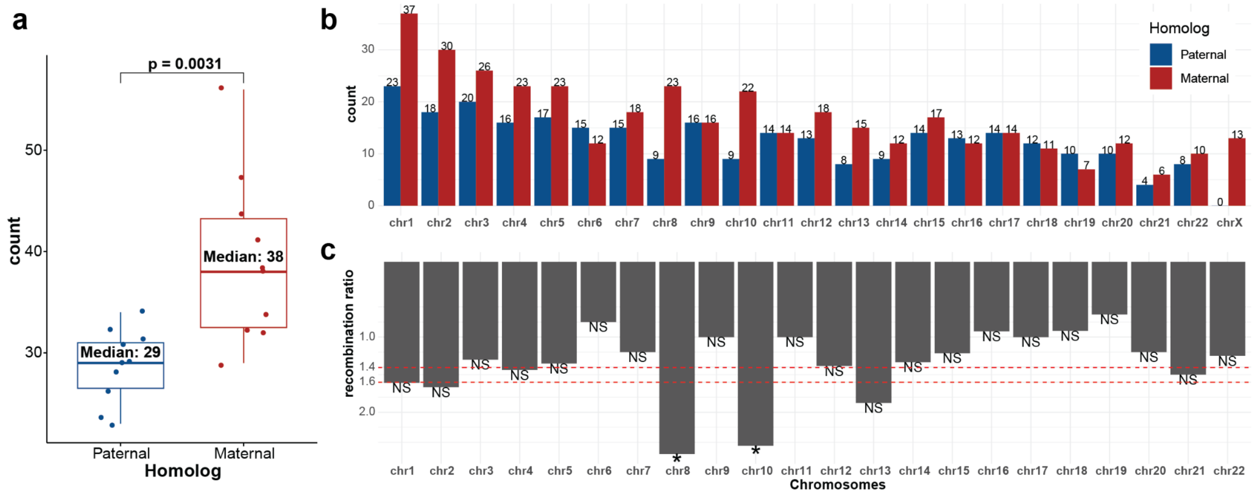

**Supplementary Figure 29: Maternal and paternal recombination counts and their ratios per chromosome.**

**a**) Distribution of total maternal and paternal recombination breakpoints for each G2 and G3 sample (n=10). T-test (two-sided) p-value is shown on top. **b**) A barplot showing a total number of maternal (red) and paternal (blue) recombination breakpoints across G2 and G3 samples (n=10). **c**) A barplot showing the maternal versus paternal breakpoint ratios (based on count from top panel). Significance was evaluated using z-score calculation followed by p-value transformation expecting two-tailed distribution.

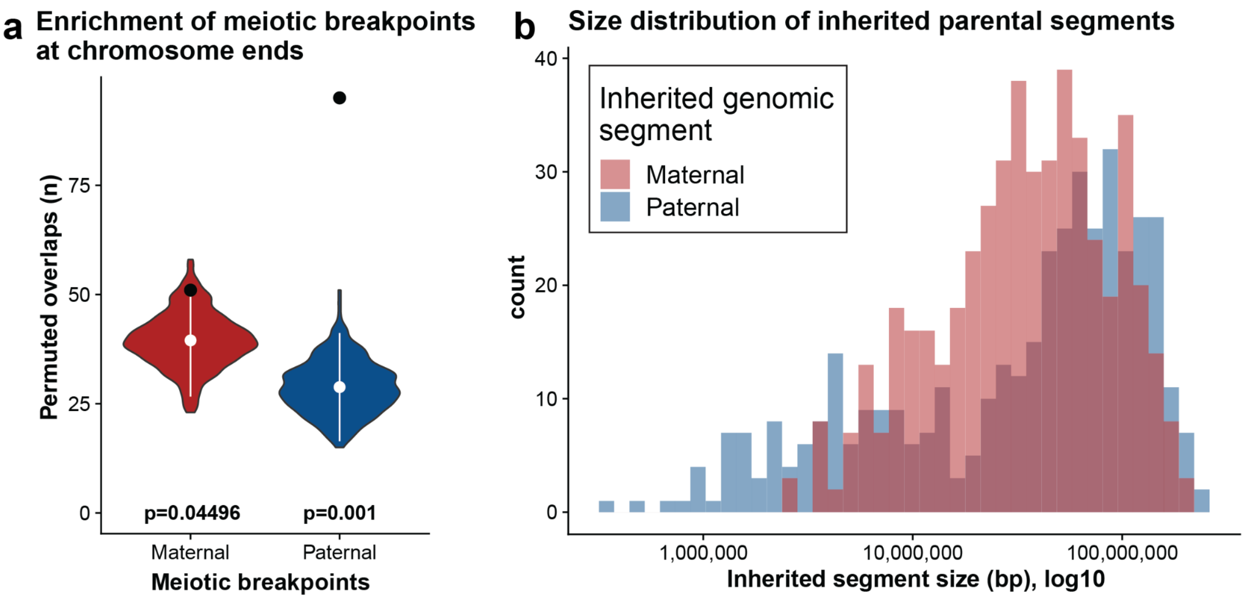

**Supplementary Figure 30: Recombination breakpoint distribution biases (T2T-CHM13).**

**a**) A violin plot showing the distribution of shuffled meiotic recombination counts at ends of each chromosome (last 5% size of each chromosome; **Methods**). The white points show the mean value of the distribution while the black point shows observed counts of the original breakpoint position at the defined chromosome ends. At the bottom of each violin plot there is a p-value (permutation test) associated with the differences between observed and rescheduled counts of recombination breakpoints at chromosome ends. **b**) A histogram showing the distribution of maternally (red) and paternally (blue) inherited DNA segments across all G2 and G3 samples. Note: Segment sizes on x-axis are reported after log10 transformation.

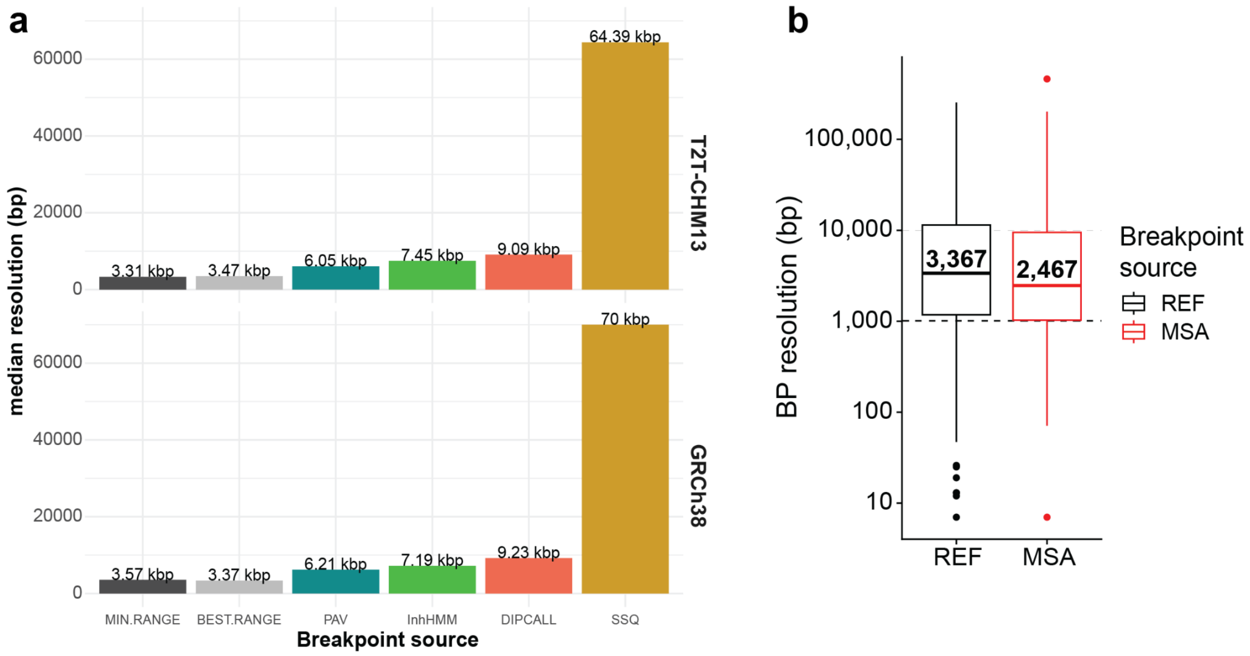

**Supplementary Figure 31: Recombination breakpoint resolution.**

**a**) A barplot showing a median breakpoint resolution separately for recombination maps reported by different orthogonal datasets (assembly-based: PAV and Dipcall; inheritance-vector-based: InhHMM and Strand-seq-based: SSQ). There is also a so-called ‘best.range’—the narrowest breakpoint range among all datasets that overlap the reference breakpoint. Lastly, we report the median value for so-called ‘min.range’—the range with the highest coverage among all datasets. **b**) Summary of refined recombination breakpoints (n=487) using phased genome assemblies and multiple sequence alignment (MSA) between sequence extracted from the parental and inherited homolog in the child as opposed to breakpoints defined with respect to a single reference (REF: T2T-CHM13 reference).

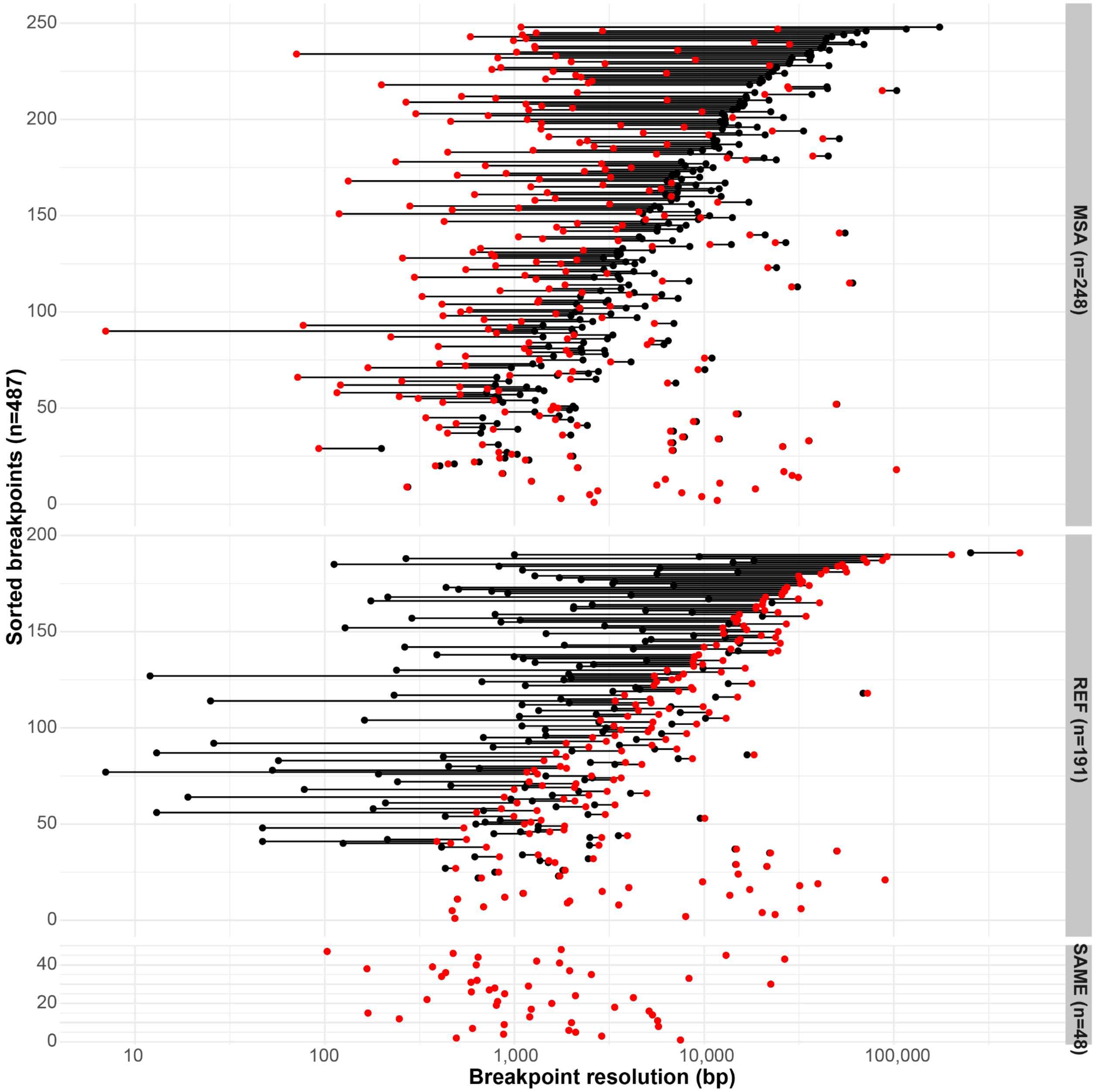

**Supplementary Figure 32: Recombination breakpoint refinement using phased genome assemblies.**

Size distribution of refined recombination breakpoints (n=479) from the total of 539 recombination breakpoints in G3 (with respect to T2T-CHM13). Resolution of original reference breakpoints are marked by black points while the refined breakpoints are marked by red points. Original and refined breakpoints are connected by horizontal lines. Breakpoints with improved resolution after multiple sequence alignment (MSA) analysis are on top (MSA, n=248). Breakpoints in the expanded recombination region after MSA analysis are shown in the middle (REF, n=191). Last, breakpoints with largely unchanged resolution are shown at the bottom (SAME, n=48).

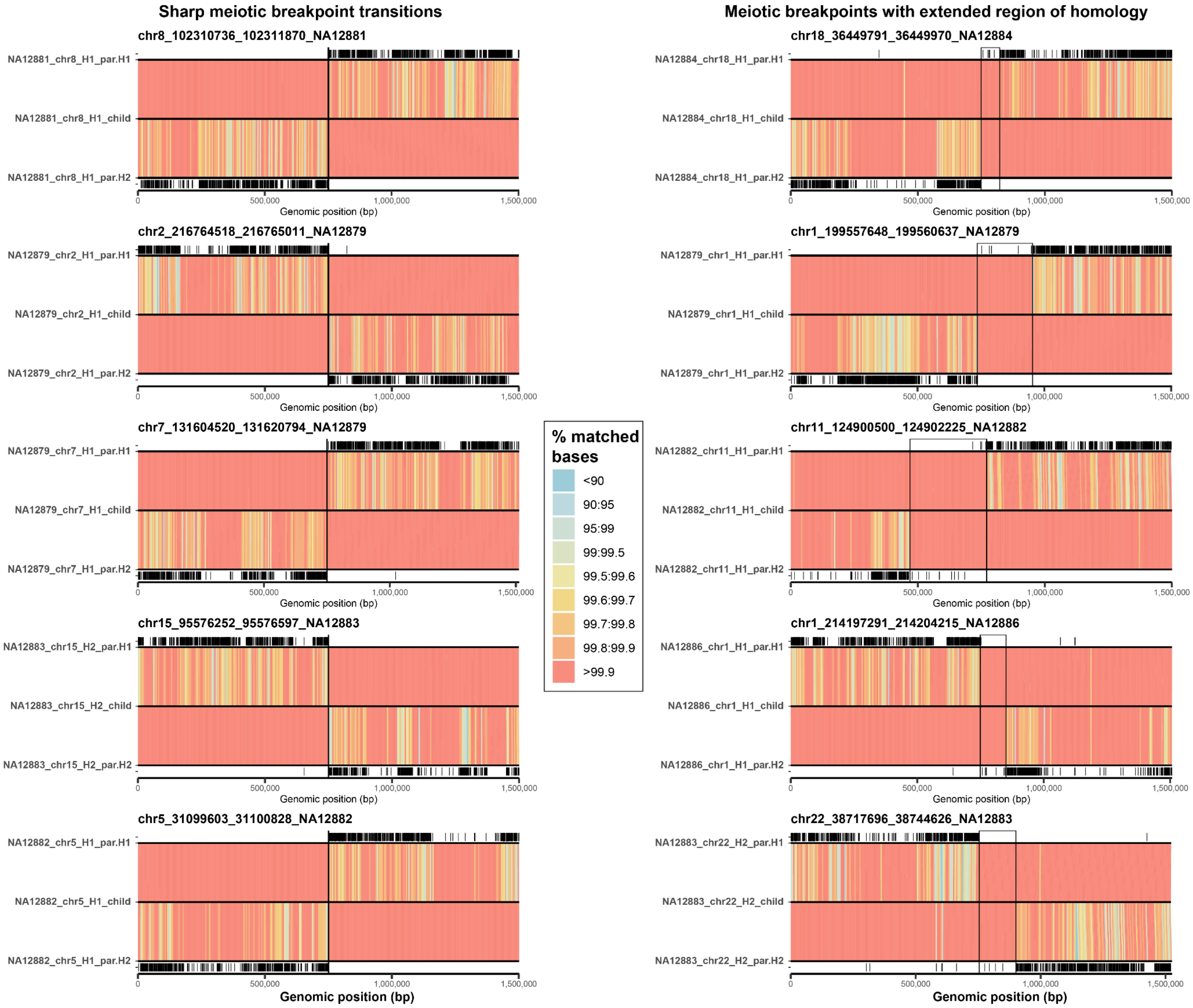

#### Supplementary Figure 33: Sharp and wide transition at recombination breakpoints.

Alignments between the parental and child haplotypes are binned into 5 kbp long bins and colored based on the percentage of matched bases. Black tick marks show positions of mismatches between parental and child haplotypes. We present five examples of sharp recombination breakpoint transitions (left) as well as meiotic breakpoints with extended regions of homology (right). Extended regions of homology at the recombination breakpoints are highlighted by black rectangles.

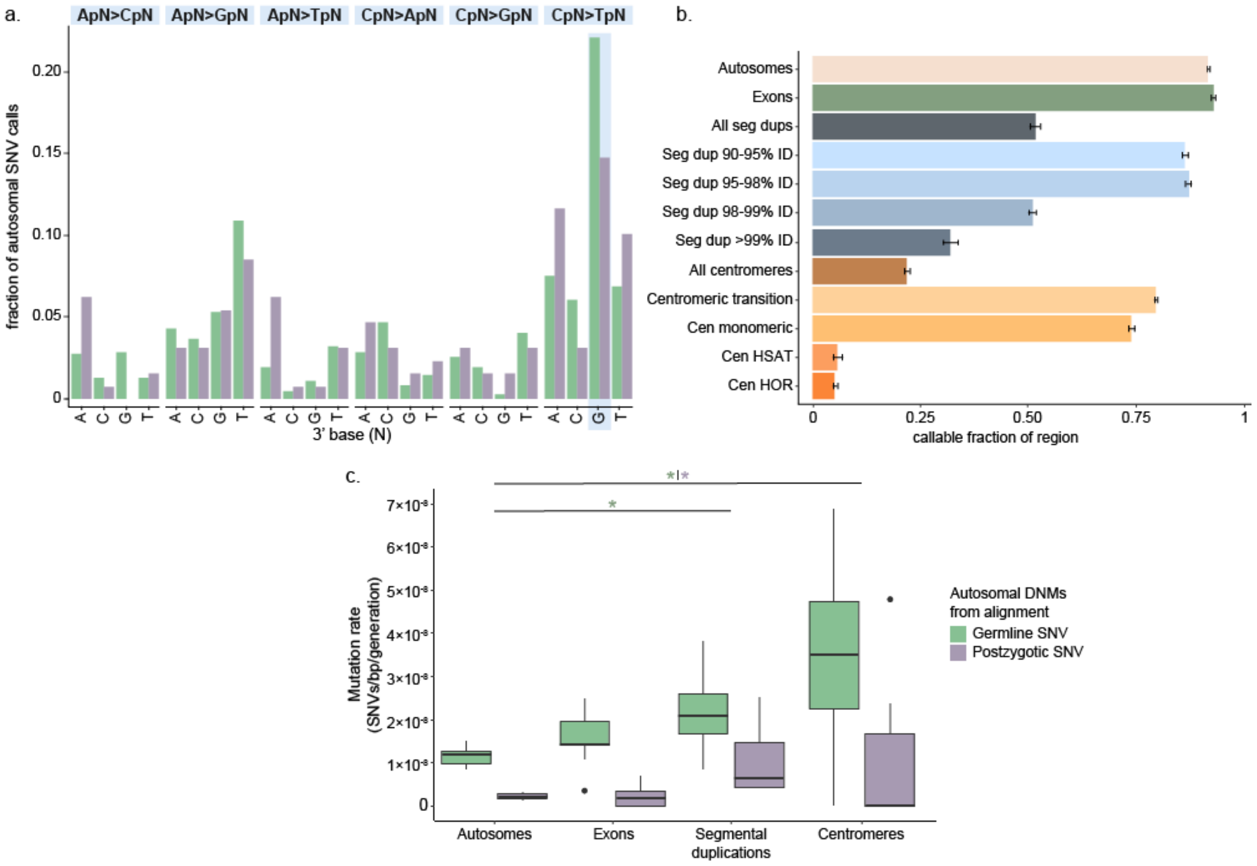

#### Supplementary Figure 34: Germline and postzygotic mutation spectrum and rates.

**a**) Dinucleotide mutation spectrum of germline *de novo* SNVs (green) and postzygotic SNVs (purple). No differences in spectra rose to significance. **b**) Mean amount of callable genome across samples, based on aligned HiFi reads. The black bars represent the first standard deviation. **c**) Per-sample mutation rates across different regions of the autosomes reveal significant enrichment in segmental duplications and centromeres.

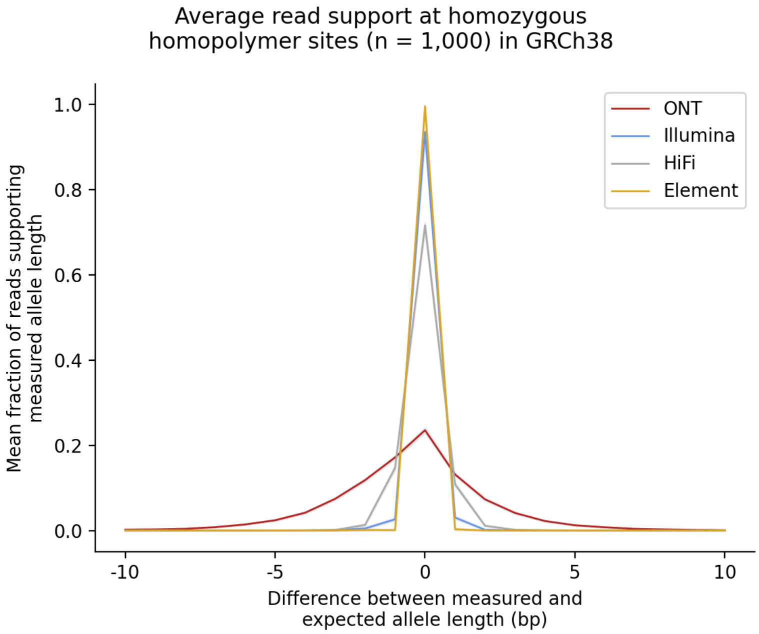

**Supplementary Figure 35: “Stutter” profiles at homozygous homopolymer loci using various sequencing technologies on sample G3-NA12879.**

At each locus, we counted the net CIGAR operations in reads that overlapped the short tandem repeat (STR) locus; we considered the net total of CIGAR operations to be the “measured” allele length of the STR. We then compared the “measured” allele length to the “expected” allele length at each locus derived from TRGT output. An average of 99.5% (bootstrap 95% CI = 99.1-99.8%) of Element reads perfectly support the TRGT allele length followed by 93.5% (95% CI = 92.8 - 94.0) of Illumina reads; 71.6% (95% CI = 70.8 - 72.3) of HiFi reads; 23.5% (95% CI = 22.9 - 24.2) of ONT reads.

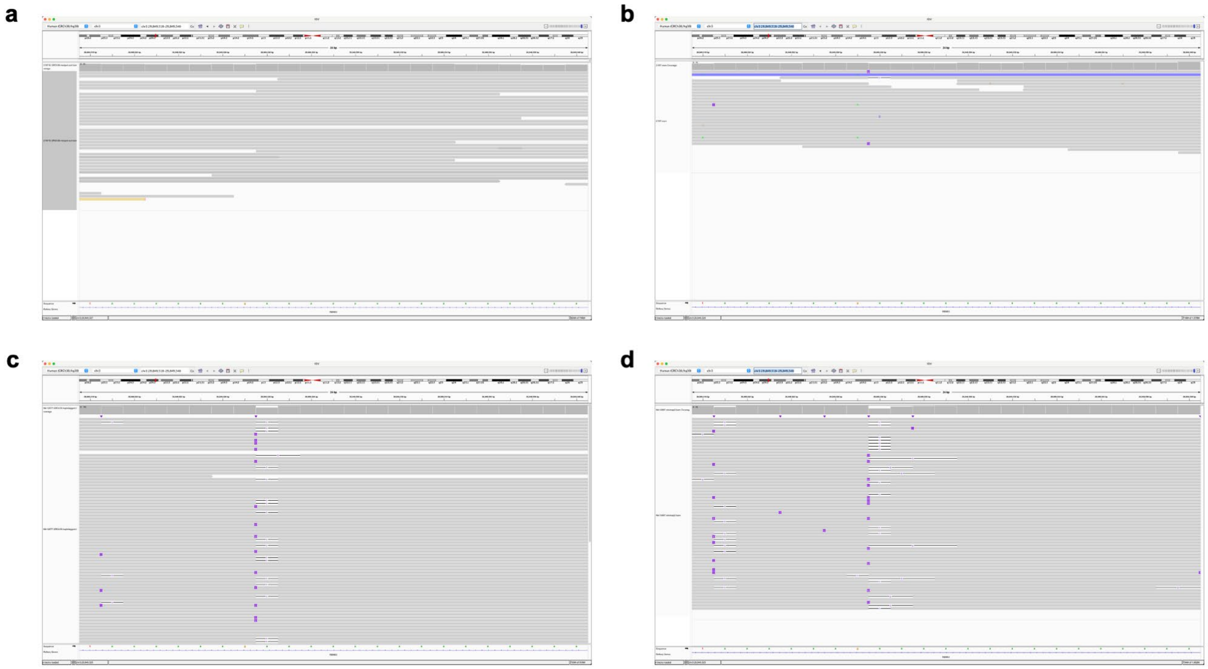

**Supplementary Figure 36: Read evidence from orthogonal technologies at a single homopolymer locus.**

IGV screenshots of read evidence from **a**) Element AVITI, **b**) Illumina NovaSeq, **c**) PacBio HiFi, and **d**) ONT at a 22 bp $(A)_n$ homopolymer locus in GRCh38.

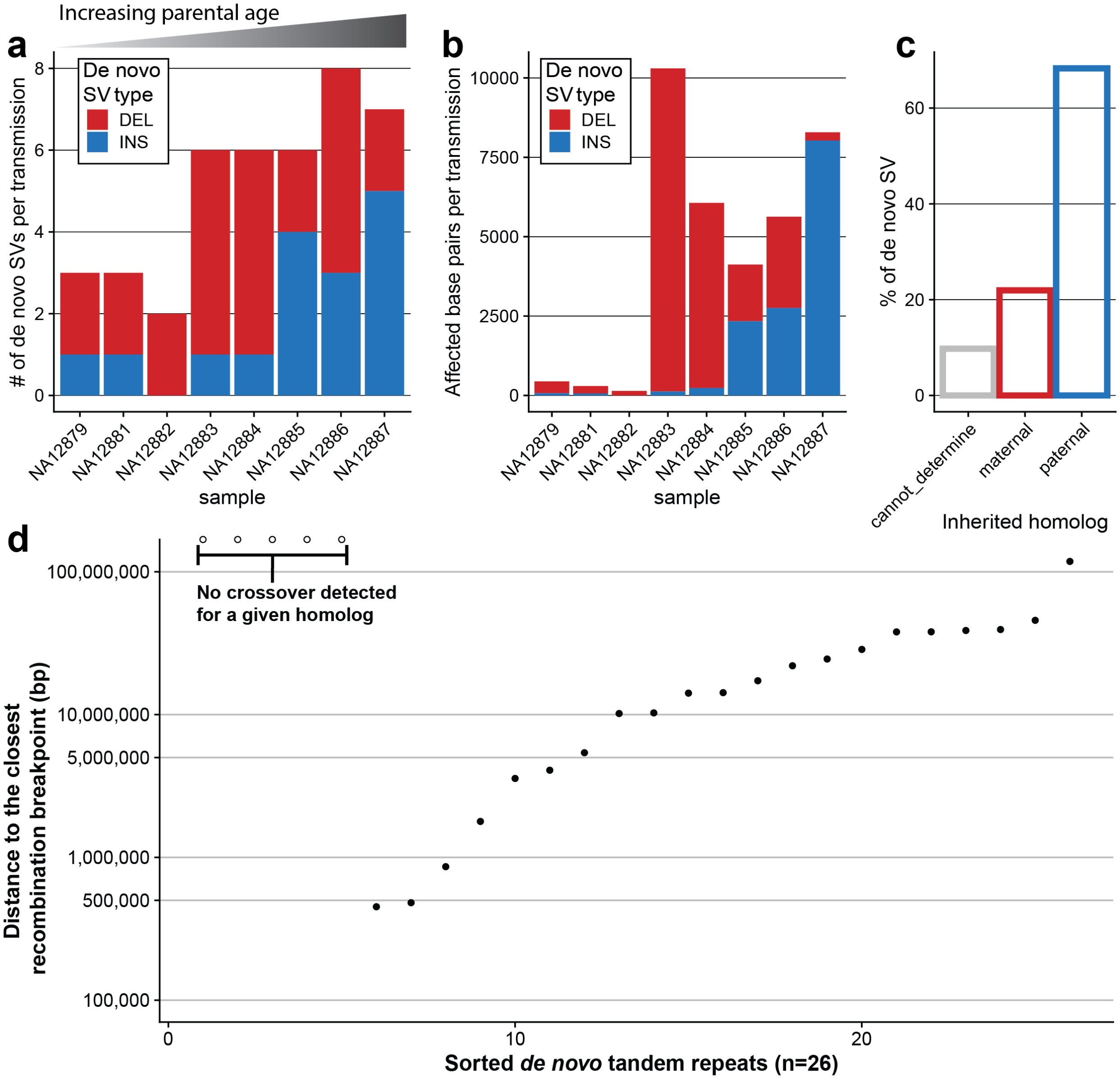

**Supplementary Figure 37: Summary of detected *de novo* SVs (n=41).**

**a**) A barplot showing the total number of *de novo* SVs per G3 sample as stacked insertion (INS) and deletion (DEL) counts. **b**) A barplot showing the total number of bases affected by *de novo* SVs per G3 sample as stacked insertion (INS) and deletion (DEL) base-pair counts. **c**) A barplot showing the percentage of *de novo* SVs inherited from paternal (blue) or maternal (red) homologs. The gray bar shows *de novo* SVs where inheritance could not be reliably determined. **d**) Distribution of distances of *de novo* tandem repeats from detected cross-over in a given sample and chromosome. Empty points mark *de novo* TRs where no crossover was detected for a given chromosome in a given sample.

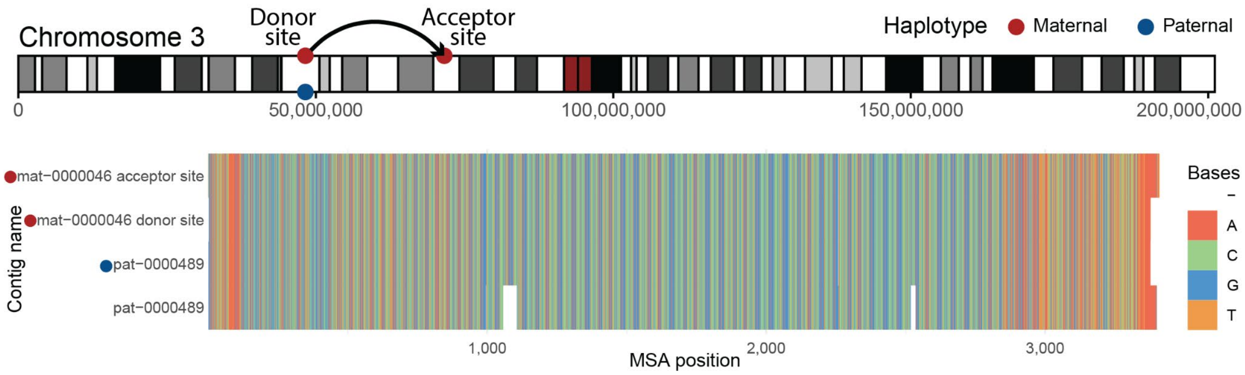

**Supplementary Figure 38: Predicting a donor site of *de novo* SVA insertion.**

**Top**: Predicted donor and acceptor sites of the SVA element on the chr3 ideogram. **Bottom**: An MSA between all SVA insertion positions in the *de novo* assembly of G3-NA12887. All positions of the SVA insertion were defined by mapping the inserted sequence against the *de novo* assembly of G3-NA12887 using minimap2 with the following parameters: `-x asm20 -c --eqx --secondary=yes`. Then the sequence of the SVA element was extracted from both maternal (red) and paternal (blue) assemblies and used to construct the MSA.

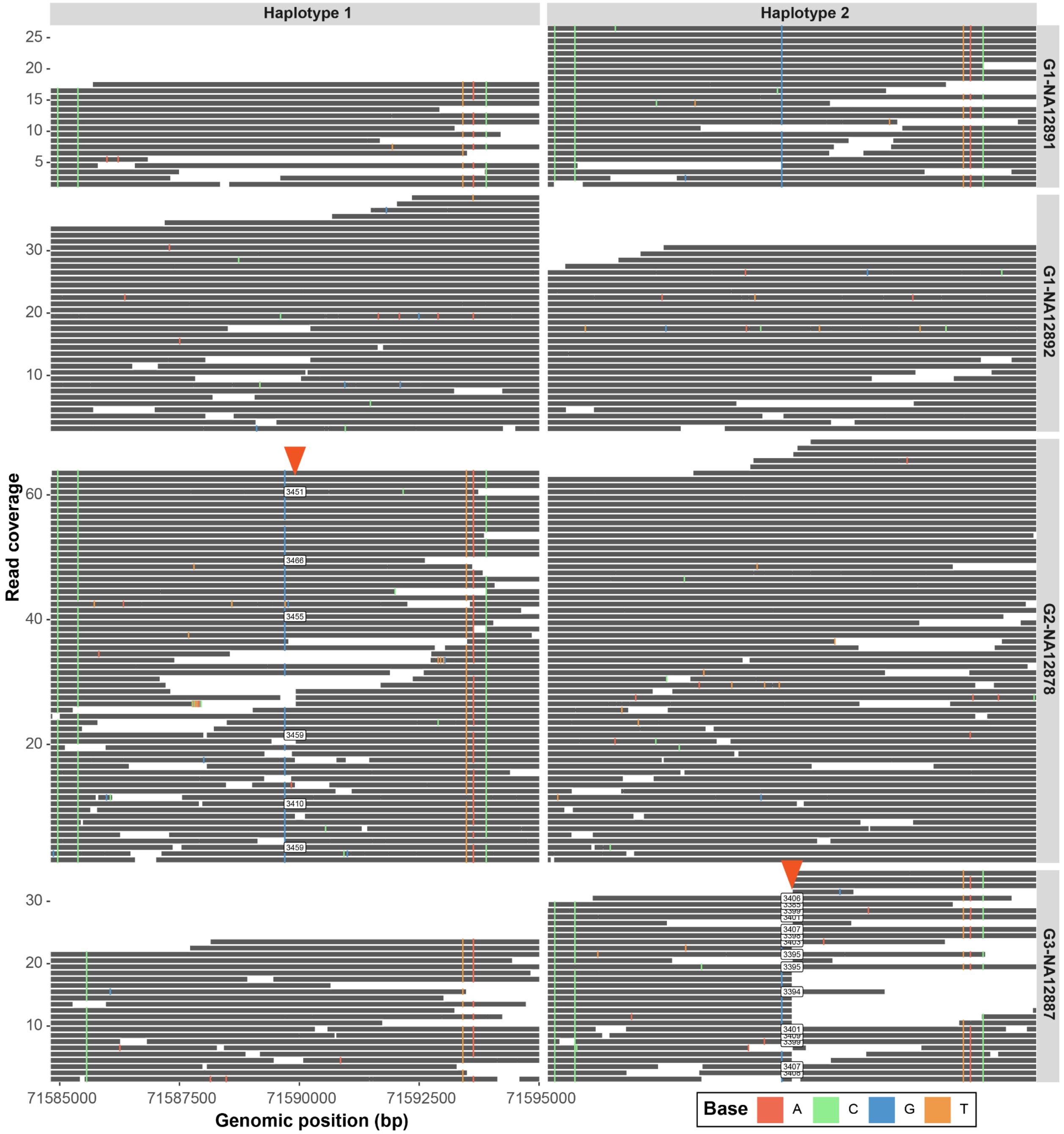

**Supplementary Figure 39: Predicting a donor site of *de novo* SVA insertion.**
Visualization of HiFi reads aligned to the T2T-CHM13 reference for G1-G3 samples (G1 - NA12891, NA12892, G2-NA12878, and G3-NA12887) over the region (chr3:71584799-71595019) where the *de novo* SVA insertion was discovered. Each horizontal gray line represents a single HiFi read. Labels containing a number highlight the position and size of the SVA insertion (red arrowhead) in a given read. HiFi reads are stratified per sample (rows) and per haplotype (columns).

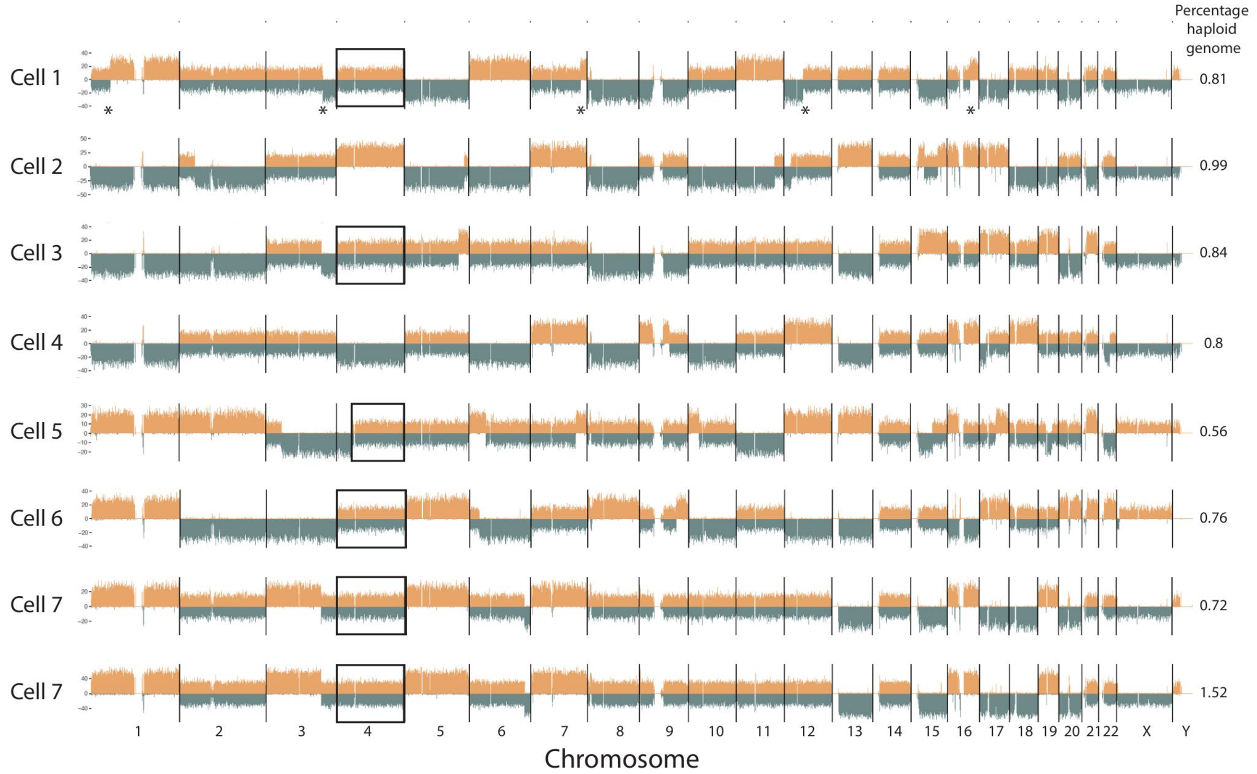

**Supplementary Figure 40: Example of Strand-seq libraries.**

Sequence reads from seven individual cells from GM12877, plotted using BreakpointR (Porubsky et al. 2019), illustrate the power of Strand-seq libraries constructed with restriction enzymes for genome analysis. The number of reads in 0.2 Mbp bins mapping to the plus strand (Crick, top, orange) or the minus strand (Watson, bottom, teal) of the hg38 reference genome are plotted. Asterisks point to sister chromatid exchange events in cell 1. Boxes around chromosome 4 point to cells with suitable data for generating chromosome-length haplotypes for this chromosome. Note that large inversions on chromosomes 8 and 16 are easily recognized in a subset of the cells. For each individual, around 100 Strand-seq libraries were made, of which on average 80% passed ASHLEYS quality control criteria (Gros et al. 2021). Libraries of around 2000 cells were made in two library construction experiments, which included 14 individuals from G1-G3. One of the library pools was sequenced on the AVITI from Element Biosciences (San Diego, CA). Results obtained for the same cell on the NextSeq 550 and the AVITI are shown in the bottom two rows (results with AVITI at the bottom).

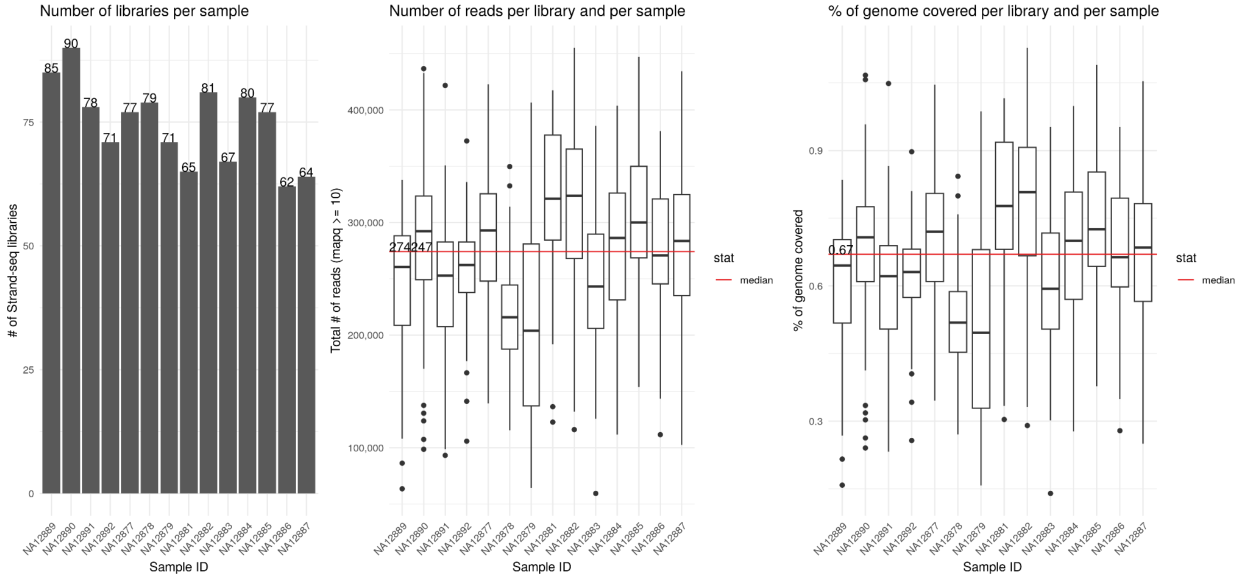

**Supplementary Figure 41: Strand-seq data summary.**

**Left:** Number of selected libraries per sample (G1-G3).

**Middle:** Distribution of mapped reads to the reference (T2T-CHM13) with mapping quality ≥10 per single-cell library and per sample.

**Right:** Distribution of percentage of reference genome (T2T-CHM13) covered by at least one read per single-cell library and per sample.

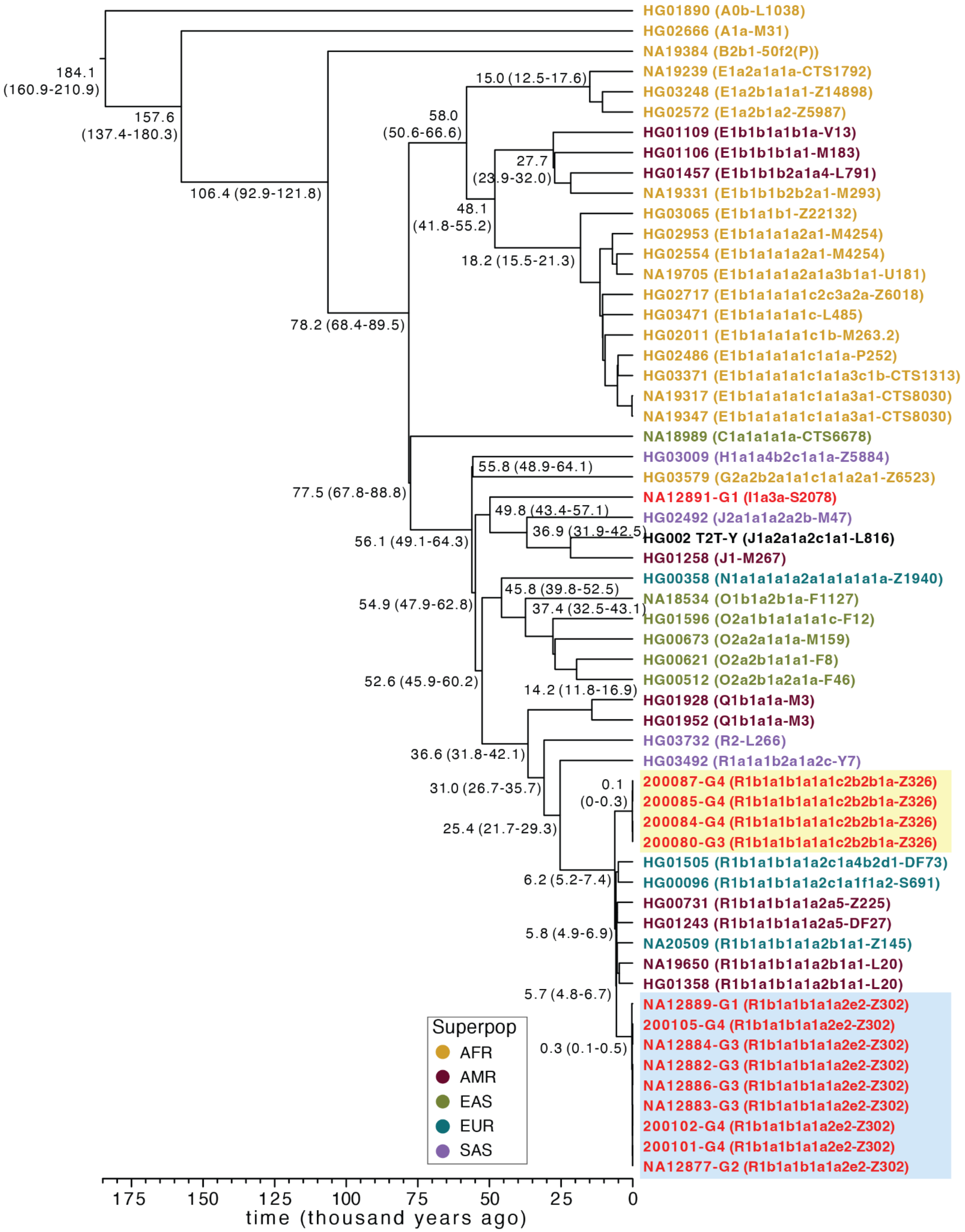

**Supplementary Figure 42: Phylogenetic relationships of long-read Y assemblies and pedigree Y chromosomes.**

Y chromosomes from 14 pedigree males are combined with 44 individuals for which long-read-based Y assemblies have previously been published (Hallast et al. 2023). Split times as estimated according to the BEAST analysis are shown for major splits with 95% highest posterior density (HPD) intervals in brackets. Red text - indicates pedigree males, other colors indicate the 1000 Genomes Project continental groups. Yellow background shows the Y chromosomes of G3 spouse (200080) and his three male offspring. Blue background indicates the nine males with R1b1a-Z302 Y chromosomes analyzed in detail here (**Fig. 5a**).

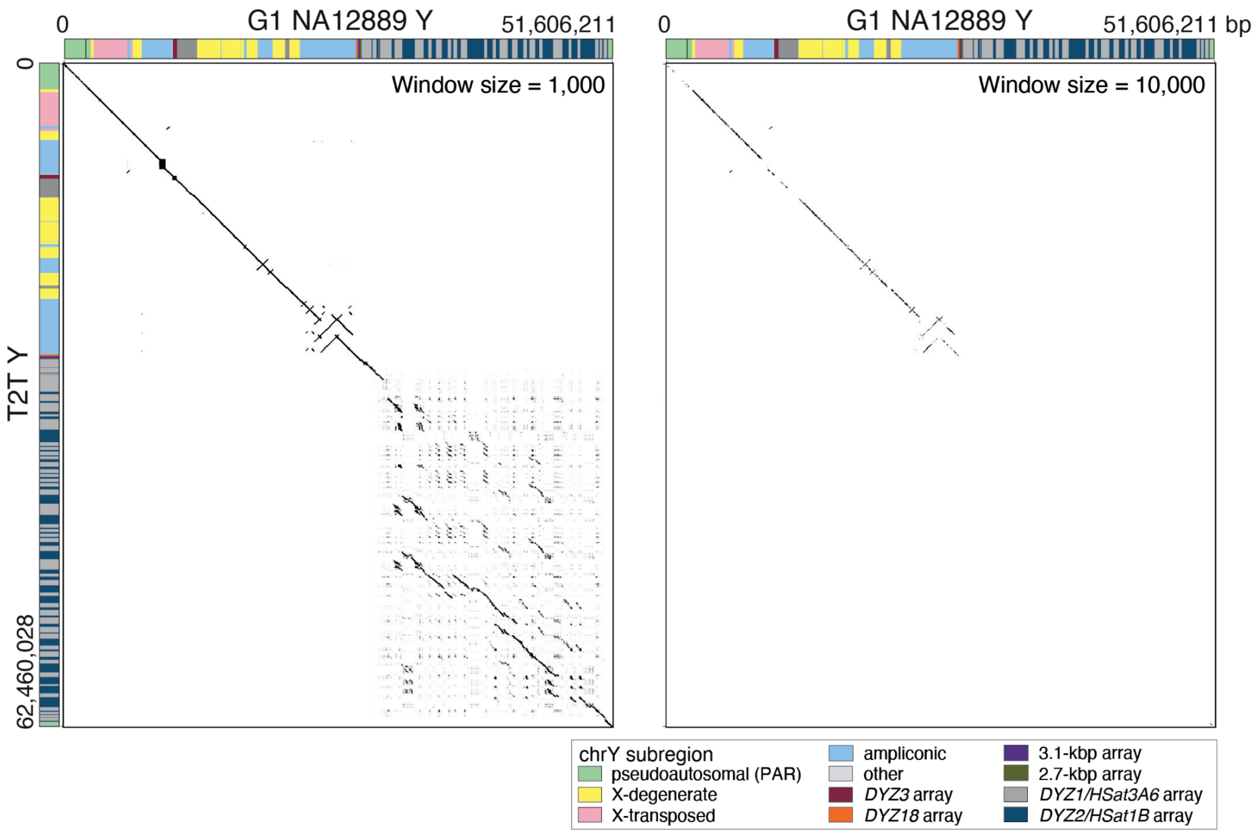

#### Supplementary Figure 43: Comparison of G1-NA12889 and T2T-CHM13 Y chromosome sequences.

Dot plots of sequence similarity between the G1-NA12889 Y assembly and the T2T-CHM13 Y chromosome sequence (Rhie et al. 2023). The T2T Y (J1a-L816 Y haplogroup) last shared a common ancestor with the pedigree R1b1a-Z302 Y chromosome approximately 54,900 years ago (95% HPD interval = 45,900–60,200 years ago; **Supplementary Fig. 42** and differs extensively especially in the repetitive regions of the Y chromosome. Y-chromosomal sequence classes are shown as colored bars, with the assembly break in the PAR1 of NA12889 indicated by a black line. The dot plot is generated with word size of 1,000 bp shown on the left and word size of 10,000 bp on the right.

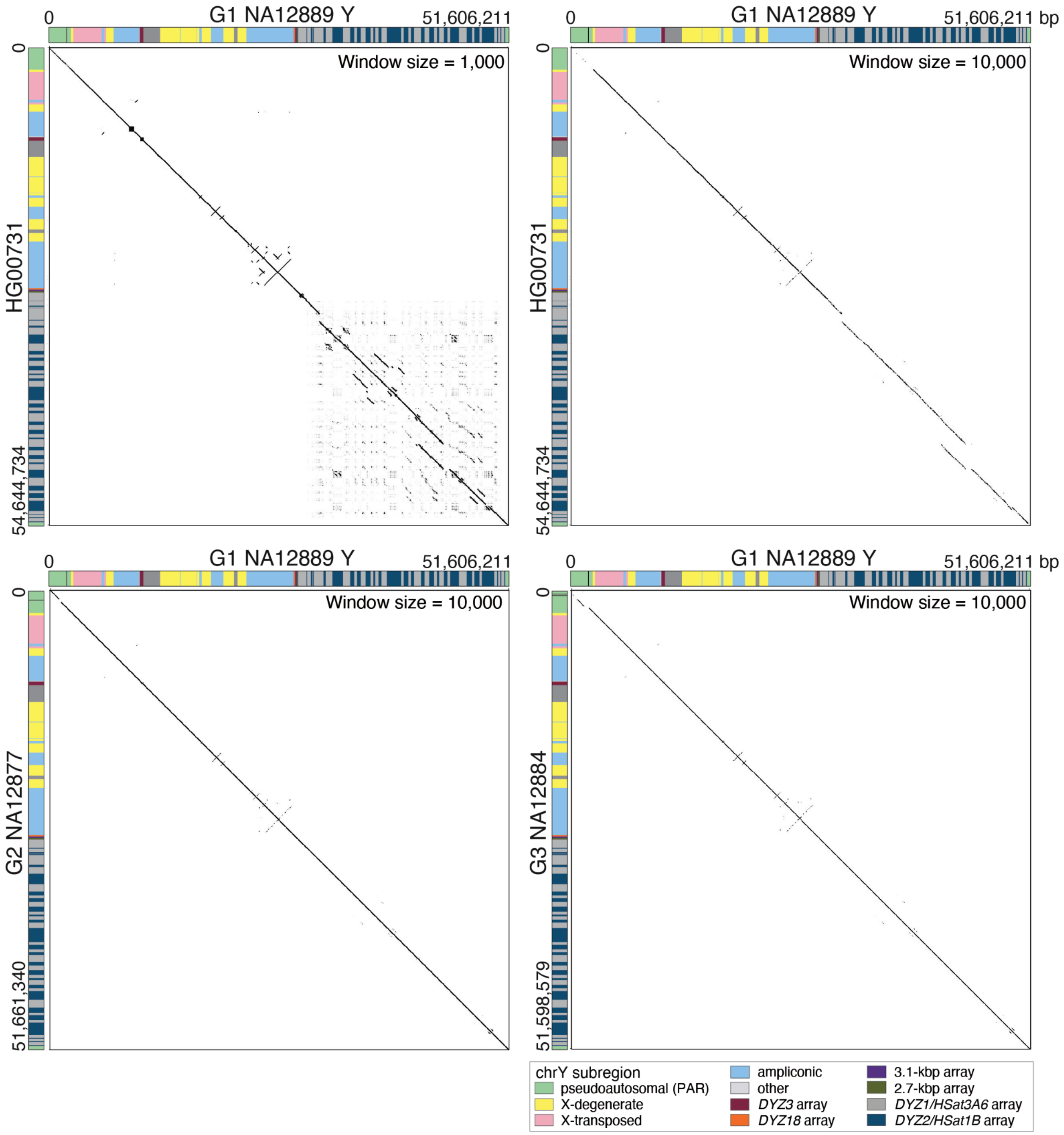

**Supplementary Figure 44: Comparison of chrY assemblies.**

Dot plots of sequence similarity between the G1 NA12889 Y assembly, evolutionarily closely related Y chromosome from HG00731 (top), G2-NA12877 (bottom left), and G3-NA12884 (bottom right) assemblies. The HG00731 Y (R1b1a-Z225 Y haplogroup) last shared a common ancestor with the pedigree R1b1a-Z302 Y chromosome approximately 5,700 years ago, 95% HPD interval = 4,800–6,700 years ago, **Supplementary Fig. 42**. Window sizes of 1,000 and 10,000 bp are shown for HG00731, and 10,000 bp for G2 and G3 males, indicating high levels of sequence similarity between the Y assemblies. Y-chromosomal sequence classes are shown as colored bars, with assembly breaks in the PAR1 indicated by black lines.

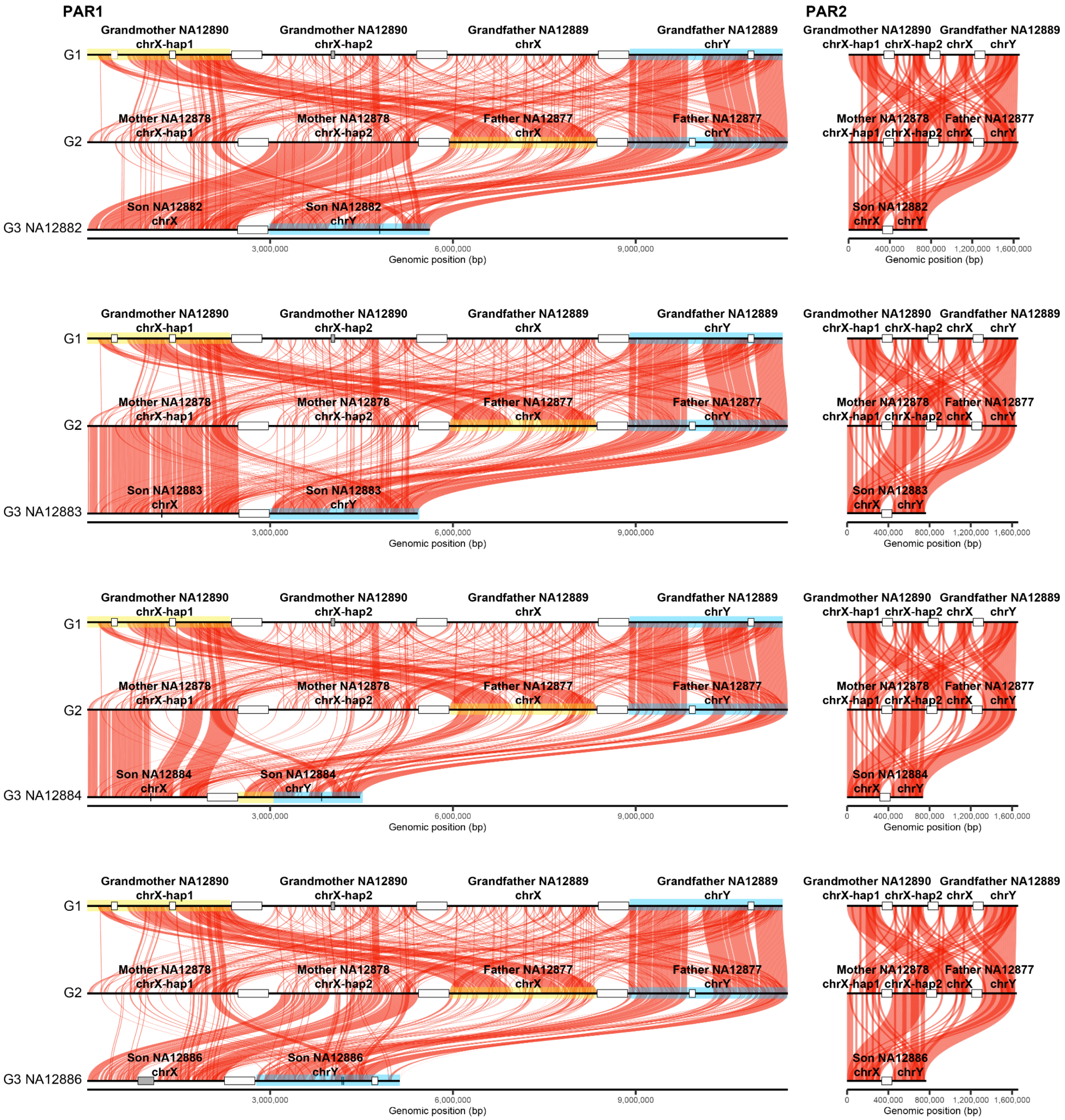

#### Supplementary Figure 45: Assembled chrX and chrY pseudoautosomal regions (PAR) across three generations.

Binned alignments (10 kbp bin size) of ≥99.9% sequence identity are shown for PAR1 (left) and PAR2 (right) across generations. For G1 and G2, maternal sequences for chrX haplotypes (hap) 1 and 2, and paternal chrX and chrY PAR sequences are shown. For each of the G3 males, chrX and chrY PAR sequences are shown. Large white rectangles (equal in size to 500 kbp) separate the haplotypes, while 100 kbp-sized rectangles indicate where joints were made if several contigs represented the region for a specific individual. Gray rectangles indicate blocks of N's in the contigs. Yellow and blue rectangles indicate chrX and chrY PAR1 haplotypes that show evidence of recombination in G3 male NA12884. No recombination events were identified in PAR2. The following contigs were included for PAR1 (in the order as visualized from left to right): G1 NA12890 chrX haplotype 1 - haplotype1−0000079, haplotype1-0000078 and haplotype1-0000010, chrX haplotype 2 - haplotype2-0000100; G1 NA12889 chrX - haplotype1-0000018 and chrY - haplotype2-0000082 and haplotype2-0000081; G2 NA12878 chrX haplotype 1 - mat-0000002, chrX haplotype 2 - pat−0000758; G2 NA12877 chrX - mat−0000005 and chrY - pat−0000406 and pat−0000383; G3 NA12882 chrX - mat−0000046 and chrY - pat−0000587; G3 NA12883 chrX - mat−0000038 and chrY - pat−0000576; G3 NA12884 chrX - mat−0000008 and chrY - pat−0000224; G3 NA12886 chrX - mat−0000010 and chrY - pat−0000781 and pat−0001035; and for PAR2: G1 NA12890 chrX haplotype 1 - haplotype1-0000010, chrX haplotype 2 - haplotype2-0000096, G1 NA12889 chrX - haplotype1-0000018, chrY - haplotype2-0000081; G2 NA12878 chrX haplotype 1 - mat-0000002, chrX haplotype 2 - pat-0000758; G2 NA12877 chrX - mat-0000005, chrY - pat-0000383; G3 NA12882 chrX - mat-0000046 and chrY - pat-0000570; G3 NA12883 chrX - mat-0000045 and chrY - pat-0000567; G3 NA12884 chrX - mat-0000008 and chrY - pat-0000224; G3 NA12886 chrX - mat-0000010 and chrY - pat-0000749.

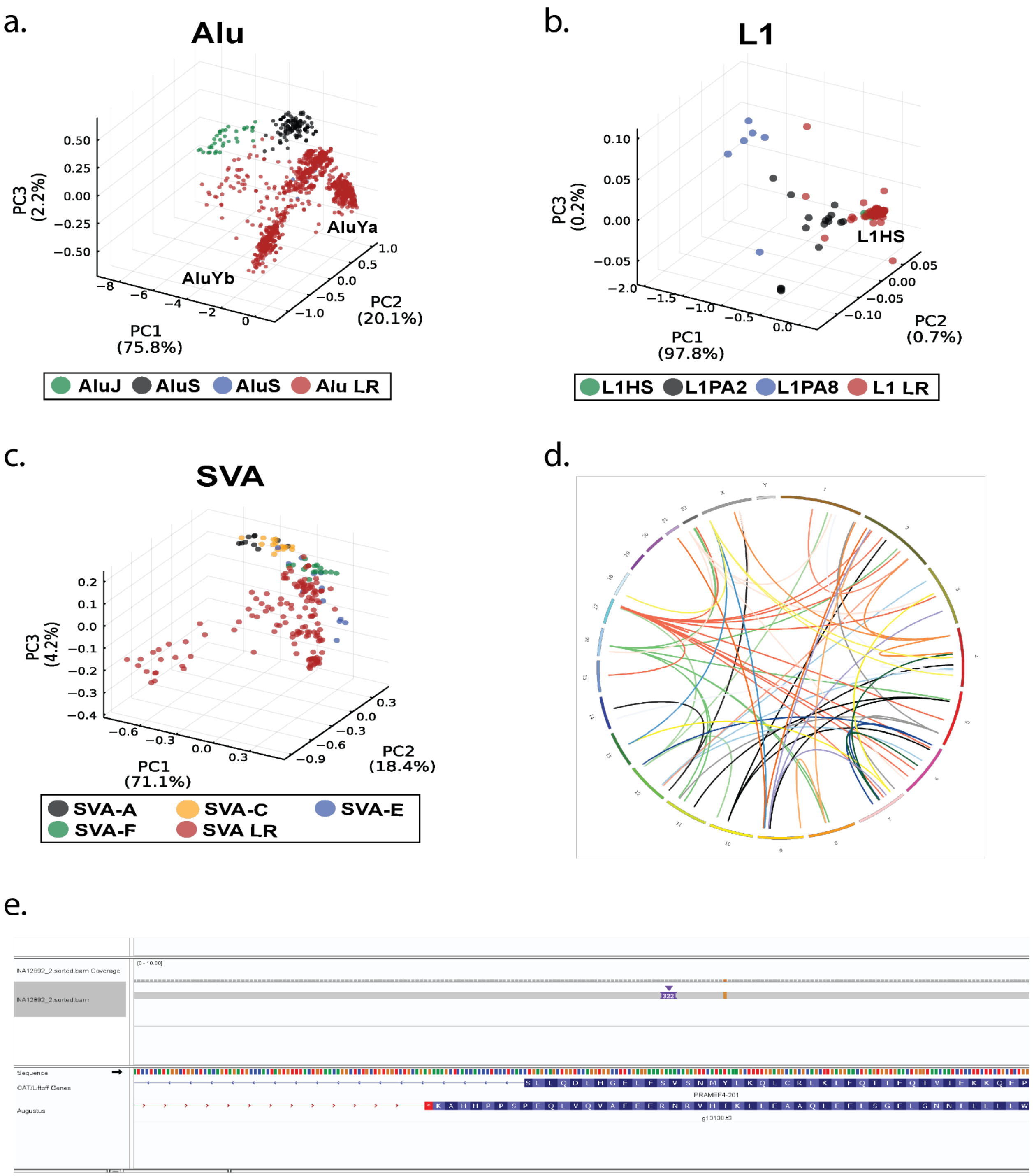

#### Supplementary Figure 46: MEI analysis summary.

**a-c**) Principal component analyses of Alu, L1, and SVA insertions discovered in long-read (LR) sequencing data. For each MEI class, new (non-reference) MEI elements are shown in red. Selected MEIs from known subclasses are shown by additional colors. **a**) 2,158 new Alu insertions cluster predominantly with known AluY elements and show distinct clustering of AluYa and AluYb subfamilies, **b**) 112 new full-length L1s insertions cluster with mostly with L1HS sequences, and **c**) 199 new SVA elements show highest affinity to known SVA-E and SVA-F elements. Locations of selected subfamilies' clusters are shown on the plot for clarity. In general, the new MEIs are representative of MEI subfamilies known to be active in human lineages (e.g., AluY, L1HS, SVA-E/F). Collectively, these MEIs represent ~1.7 Mbp of new non-reference MEI sequence (Alu: 605.4 kbp, L1: 687.1 kbp, and SVA 447.7 kbp) discovered using long-read sequencing technologies. **d**) Circos plot of the source elements responsible for two or more of the non-reference LINE-1 insertions identified in this study. The 20 loci represented in the plot are responsible for approximately 78% of the full-length non-reference LINE-1 insertions that had identifiable source elements. **e**) IGV image of a non-reference *Alu* insertion in an exon of *PRAMEF4* in an assembled genome.
